# Flexible mixture model approaches that accommodate footprint size variability for robust detection of balancing selection

**DOI:** 10.1101/645887

**Authors:** Xiaoheng Cheng, Michael DeGiorgio

## Abstract

Long-term balancing selection typically leaves narrow footprints of increased genetic diversity, and therefore most detection approaches only achieve optimal performances when sufficiently small genomic regions (*i.e*., windows) are examined. Such methods are sensitive to window sizes and suffer substantial losses in power when windows are large. This issue creates a tradeoff between noise and power in empirical applications. Here, we employ mixture models to construct a set of five composite likelihood ratio test statistics, which we collectively term *B* statistics. These statistics are agnostic to window sizes and can operate on diverse forms of input data. Through simulations, we show that they exhibit comparable power to the best-performing current methods, and retain substantially high power regardless of window sizes. They also display considerable robustness to high mutation rates and uneven recombination landscapes, as well as an array of other common confounding scenarios. Moreover, we applied a specific version of the *B* statistics, termed *B*_2_, to a human population-genomic dataset and recovered many top candidates from prior studies, including the then-uncharacterized *STPG2* and *CCDC169-SOHLH2*, both of which are related to gamete functions. We further applied *B*_2_ on a bonobo population-genomic dataset. In addition to the *MHC-DQ* genes, we uncovered several novel candidate genes, such as *KLRD1*, involved in viral defense, and *SCN9A*, associated with pain perception. Finally, we show that our methods can be extended to account for multi-allelic balancing selection, and integrated the set of statistics into open-source software named `BalLeRMix` for future applications by the scientific community.

## Introduction

Balancing selection maintains polymorphism at selected genetic loci, and can operate through a variety of mechanisms (Charlesworth, 2006). In addition to overdominance (Charlesworth and Charlesworth, 2010), other processes such as sexual selection (Cho et al., 2006), periodical environmental shifts (Bergland et al., 2014), pleiotropy (Andrés, 2001; Mitchell-Olds et al., 2007), meiotic drive (Ubeda and Haig, 2004; Charlesworth and Charlesworth, 2010), and negative frequency-dependent selection (Charlesworth and Charlesworth, 2010) can also maintain diversity at underlying loci. Due to the increasing availability of population level genomic data, in which allele frequencies and genomic density of polymorphisms can be assessed in detail, there is an expanding interest in studying balancing selection and detecting its genomic footprints (*e.g*., Andrés et al., 2009; Leffler et al., 2013; DeGiorgio et al., 2014; Gao et al., 2015; Hunter-Zinck and Clark, 2015; Sheehan and Song, 2016; Lonn et al., 2017; Sweeney et al., 2017; Guirao-Rico et al., 2017; Siewert and Voight, 2017, 2018; Bitarello et al., 2018; Ye et al., 2018; Cheng and DeGiorgio, 2019). However, despite multiple efforts to design statistics for identifying balanced loci (*e.g*., DeGiorgio et al., 2014; Siewert and Voight, 2017, 2018; Bitarello et al., 2018; Cheng and DeGiorgio, 2019), performances of existing methods still leave room for improvement.

Early methods applied to this problem evaluated departures from neutral expectations of genetic diversity at a particular genomic region. For example, the Hudson-Kreitman-Aguadé (HKA) test (Hudson et al., 1987) uses a chi-square statistic to assess whether genomic regions have higher density of polymorphic sites when compared to a putative neutral genomic background. In contrast, Tajima’s *D* (Tajima, 1989) measures the distortion of allele frequencies from the neutral site frequency spectrum (SFS) under a model with constant population size. However, these early approaches were not tailored for balancing selection, and have limited power. Recently, novel and more powerful summary statistics (Siewert and Voight, 2017, 2018; Bitarello et al., 2018) and model-based approaches (DeGiorgio et al., 2014; Cheng and DeGiorgio, 2019) have been developed to specifically target regions under balancing selection. In general, the summary statistics capture deviations of allele frequencies from a putative equilibrium frequency of a balanced polymorphism. In particular, the non-central deviation statistic (Bitarello et al., 2018) adopts an assigned value as this putative equilibrium frequency, whereas the *β* and *β*^(2)^ statistics of Siewert and Voight (2017, 2018) use the frequency of the central polymorphic site instead. On the other hand, the *T* statistics of DeGiorgio et al. (2014) and Cheng and DeGiorgio (2019) compare the composite likelihood of the data under an explicit coalescent model of long-term balancing selection (Hudson et al., 1987; Hudson and Kaplan, 1988) to the composite likelihood under the genome-wide distribution of variation, which is taken as neutral.

Nevertheless, all extant approaches are limited by their sensitivity to the size of the region that the statistics are computed on (hereafter referred to as the “window size”). Because the footprints of long-term balancing selection are typically narrow (Hudson and Kaplan, 1988; Charlesworth, 2006), small windows with fixed size comparable to that of the theoretical footprint based on a genome-wide recombination rate estimate are commonly used in practice, especially for summary statistics. However, such small fixed window sizes not only lead to increased noise in the estimation of each statistic, but also render the statistic incapable of adapting to varying footprint sizes across the genome due to factors such as the uneven recombination landscape (Smukowski and Noor, 2011). Though adopting a larger window may reduce noise, true signals will likely be overwhelmed by the surrounding neutral regions, diminishing method power as shown by Cheng and DeGiorgio (2019). Available model-based approaches (DeGiorgio et al., 2014; Cheng and DeGiorgio, 2019) could have been made robust to window sizes if they instead adopted the SFS expected under a neutrally-evolving population of constant size as the null hypothesis, because their model of balancing selection for the alternative hypothesis converges to this constant-size neutral model for large recombination rates. However, this neutral model does not account for demographic factors that can impact the genome-wide distribution of allele frequencies, such as population size changes. To guard against such demographic influences, the model-based *T*_1_ and *T*_2_ statistics (DeGiorgio et al., 2014; Cheng and DeGiorgio, 2019) employ the genome-wide SFS instead, compromising the robustness against large windows. Moreover, Cheng and DeGiorgio (2019) showed that although the power of the *T*_2_ statistic decays much slower than other approaches as window size increases, the loss of power is still substantial.

In this article, we describe a set of composite likelihood ratio test statistics that are based on a mixture model (Figures 1A and B). This framework of nested models allows for robust and flexible detection of balancing selection that can augment the size of genomic regions considered in each test to best fit the data. Dependent on the types of data available, we propose a set of five likelihood ratio test statistics termed *B*_2_, *B*_2,MAF_, *B*_1_, *B*_0_, and *B*_0,MAF_, which respectively accommodate data with substitutions and derived (*B*_2_) or minor (*B*_2,MAF_) allele frequency polymorphisms, with substitutions and polymorphisms with unknown allele frequency (*B*_1_), and with derived (*B*_0_) or minor (*B*_0,MAF_) allele frequency polymorphisms only. We comprehensively evaluated their performances under an array of diverse simulated scenarios, including their powers for balancing selection with varying ages, distinct strengths and equilibrium frequencies, robustness against window sizes, and robustness against confounding factors such as demographic history, recombination rate variation, and mutation rate variation. We also compared and discussed their performances with other leading approaches—namely HKA, *β*, *β*\*, *β*^(2)^, NCD, *T*_1_, and *T*_2_. To gauge the performance of *B* statistics on empirical data, we re-examined contemporary human populations in the 1000 Genomes Project dataset (The 1000 Genomes Project Consortium, 2015) to uncover previously hypothesized candidates. Furthermore, we performed an exploratory whole-genome scan with *B*_2_ on bonobo genomic data (Prado-Martinez et al., 2013) to probe for long-term balancing selection in the other close relative of humans. We further extended our framework to consider multi-allelic balancing selection, and examined the performances of extant methods on cases of multi-locus balancing selection. Lastly, we developed the software `BalLeRMix` (BALancing selection LikElihood Ratio MIXture models) to implement these novel tests for the convenience of the scientific community.

**Figure 1:**
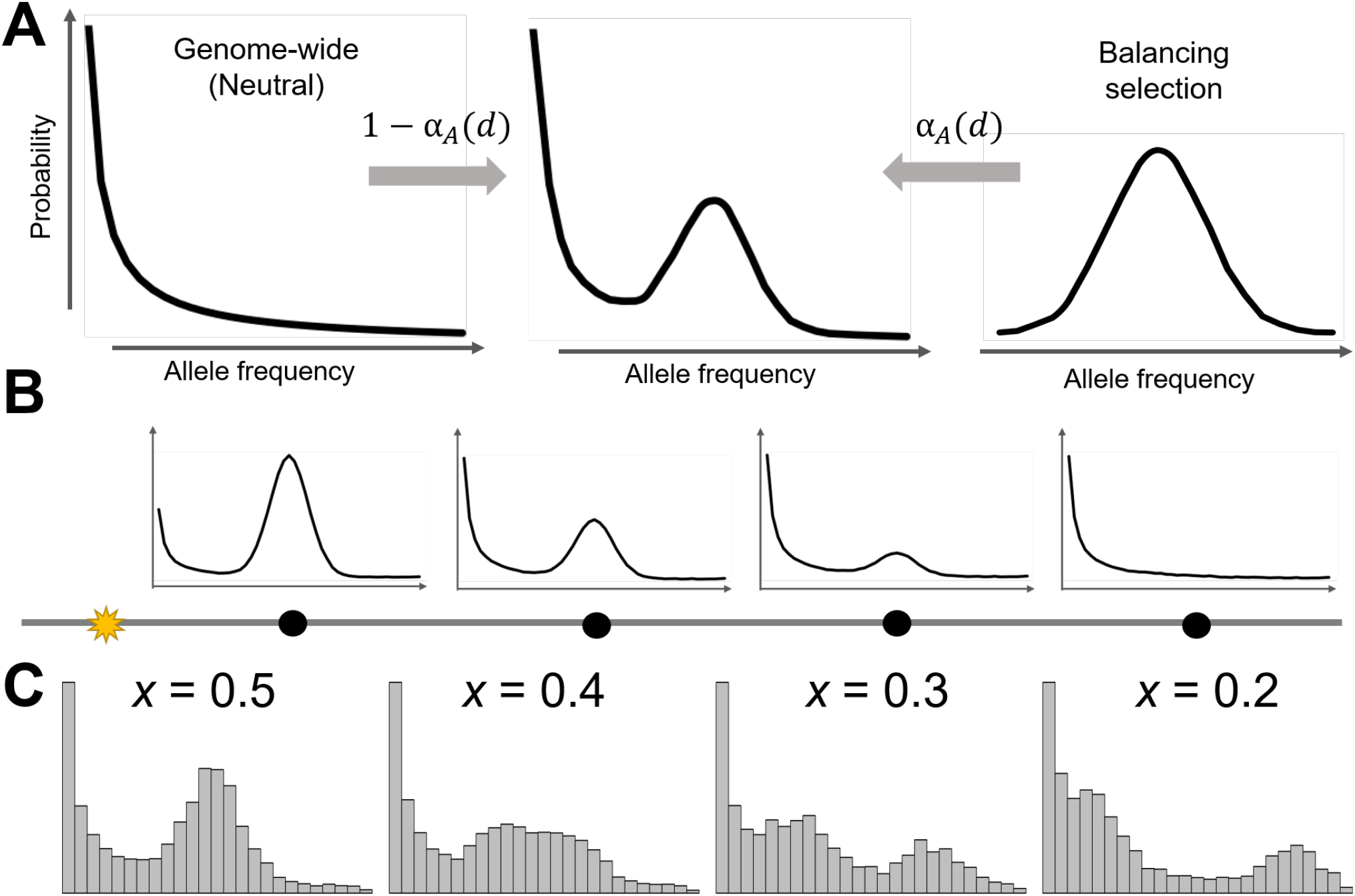
Schematic of the mixture model underlying the *B* statistics. (A) The model for the alternative hypothesis is a mixture of the distribution of allele frequencies under balancing selection at proportion *α_A_*(*d*), modeled by either a binomial or beta-binomial distribution, and the distribution under neutrality at proportion 1 − *α_A_*(*d*), modeled by the genome-wide site frequency spectrum. Here, *α_A_*(*d*) decays as a function of recombination distance *d*, and so sites close to (*i.e*, small *d*) the putative selected site will be modeled mostly by the distribution expected under balancing selection, whereas sites far from (*i.e*., large *d*) the selected site will be modeled mostly by the distribution expected under neutrality. (B) Distributions of allele frequencies at neutral sites (black dots) under the mixture model at varying distances *d* from the putative selected site (yellow star). (C) Distributions of allele frequencies from simulated sequences when balancing selection maintains the equilibrium frequency of *x* = 0.2, 0.3, 0.4, or 0.5.

## Theory

A classical footprint of balancing selection is the increase in the proportion of sites with moderate allele frequencies that are close to the equilibrium frequency at the balanced locus (Kaplan et al., 1988; Siewert and Voight, 2017). Previous modeling attempts (Kaplan et al., 1988; Song and Steinrücken, 2012; DeGiorgio et al., 2014; Cheng and DeGiorgio, 2019) primarily focused on delineating the underlying population-genetic processes, such as through coalescent or diffusion theory. Though these models are able to capture the distortion in the SFS resulting from balancing selection, their intricate mathematical formulations bring challenges to further model extensions to more complicated scenarios as well as the associated computations. As an alternative, it may be appealing to model the effect of balancing selection through statistical approximations of the expected features in the data.

Based on this idea, for a locus under balancing selection that is maintaining a pair of alleles, we can approximate the number of copies, *k*_0_, of the allele balanced at equilibrium frequency *x* ∈ (0,1) observed in *n* samples, as following a binomial distribution with *n* trials and success rate *x*. For a bi-allelic neutral site that is linked to this selected locus, we assume that the *k* derived alleles observed from the *n* samples at this neutral site are all linked to the *k*_0_ alleles balanced at frequency *x*. That is, we assume *k* = *k*_0_, and therefore *k* is also binomially distributed with *n* trials and success rate *x*. Meanwhile, for a neutral site not linked to this selected locus, we assume that *k* follows the distribution expected by the genome-wide SFS. Taken together, the probability *P_n_*(*k*) of observing *k* derived alleles out of *n* sampled alleles at a neutral site can be written as

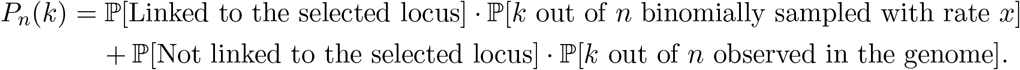

Alternatively, this integration of two conditional probabilities can also be viewed as a mixture model, in which the two mixing components represent probabilities under balancing selection and neutrality (based on the genome-wide empirical distribution), with their respective mixing proportions *α* and 1 − *α* representing the probabilities of being linked to the selected locus or not, respectively. To approximate *α*, we chose to consider the exponential decay function, which has been adopted as a proxy for linkage disequilibrium (*e.g*. Nielsen et al., 2005; Moorjani et al., 2011; Loh et al., 2013). To accommodate the varying rates of linkage decay, we introduce a free parameter *A* > 0 for the statistic to optimize over, which essentially determines the size of the footprint of balancing selection, with smaller values of *A* having wider footprints than larger values. Hence, for a neutral site *d* recombination units away from the selected locus, the probability that it is linked to the selected locus can be approximated by

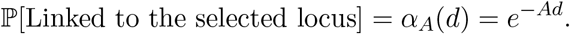

Therefore, for a neutral site *d* recombination units away from the selected locus, we approximate the probability mass function for sampling *k* derived alleles out of *n* sampled alleles as

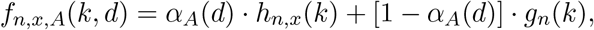

where *h_n,x_*(*k*) denotes the normalized binomial probability of sampling *k* successes out of *n* trials with success rate *x*, and *g_n_*(*k*) is the normalized genome-wide SFS denoting the proportion of sites with *k* derived alleles observed out of *n* sampled alleles. This formulation also applies when *k* represents the number of minor allele copies, for situations in which the ancestral allele cannot be polarized with an outgroup. See subsequent subsection for precise definitions of normalized *h_n,x_*(*k*) and *g_n_*(*k*).

In the following subsections, we describe a set of composite likelihood ratio statistics (*B*_2_, *B*_2,MAF_, *B*_1_, *B*_0_, and *B*_0,MAF_) constructed based on this mixture model approach for identifying loci undergoing bi-allelic balancing selection. We also extended this framework to consider multi-allelic balancing selection, and describe these models in *Supplementary Note 1*. Note that all the composite likelihood ratio statistics considered here assume that balancing selection is acting on a single locus. This set of composite likelihood ratio statistics have been implemented in the open-source software package `BalLeRMix`, which is available at https://github.com/bioXiaoheng/BalLeRMix/tree/master/software.

### Probability distributions given derived allele polymorphisms and substitutions

For *n* sampled alleles at an informative site (*i.e*., polymorphism or substitution), when the ancestral state to each site can be confidently assigned, denote the number of derived alleles as *k, k* = 1,2,…, *n*. Let *ξ_n_*(*k*) be the total number of informative sites across the whole-genome with *k* derived alleles observed out of *n* sampled alleles. The probability of observing such a site is therefore

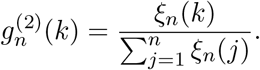

When balancing selection maintains an equilibrium frequency of *x* on the site under selection, the outcomes of observing derived alleles on this site (out of *n* lineages) can be approximated by a binomial distribution of *n* trials with a success probability of *x*. Following this binomial model, the probability of observing the selected site with *k* observed derived alleles is

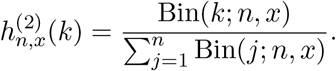

Note the values of *g_n_*(*k*) and *h_n,x_*(*k*) are conditional on the number of sampled alleles *n*, and therefore our model requires that the sample size be made explicit at each informative site. Permitting the sample size to differ across sites is important, as missing genotype calls are often common in empirical studies, with sample sizes naturally varying across the genome.

For an informative site *d* recombination units away from the presumed site under selection, it can either be linked to the derived (with equilibrium frequency *x*) or ancestral (with equilibrium frequency 1 − *x*) haplotype under balancing selection, resulting in a bimodal distribution (Figure 1C). Therefore, the probability of observing *k* derived alleles out of *n* sampled alleles is

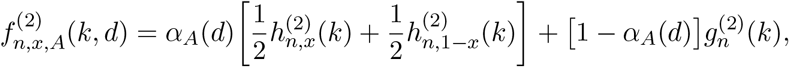

where *α_A_*(*d*) = exp(−*Ad*) and where *A* is a model parameter that determines the size of the genomic footprint of balancing selection. When allele frequency information is unavailable at polymorphic sites, the probability of observing a polymorphic site (*k* ≠ *n*) or substitution (*k* = *n*) would be

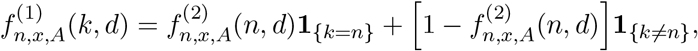

where **1**_{*E*}_ is a dummy variable that takes the value one if the expression *E* is true, and zero otherwise. Similarly, when substitutions are not considered or are missing in the data (*i.e*., only observe derived allele counts *k* = 1, 2,…, *n* − 1), the two mixing components can be normalized as

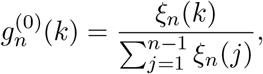

and

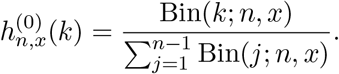

The probability of observing a polymorphic site with *k* derived alleles out of *n* sampled alleles is then

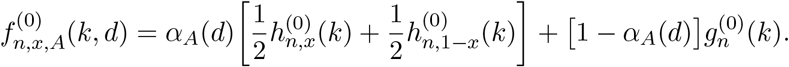

### Probability distributions given minor allele polymorphisms and substitutions

When alleles cannot be confidently polarized, minor allele frequencies are often used instead. For informative sites with *n* sampled alleles, denote the minor allele count as *k, k* =0,1,…, ⌊*n*/2⌋, and the total number of such sites in the genome as *η_n_*(*k*). Substitutions are assigned to *η_n_*(0), as the minor allele count is zero. The probability of observing a site with *k* minor alleles out of *n* sampled alleles in the genome is

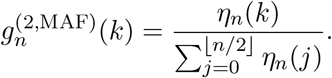

Assume the equilibrium minor allele frequency at the locus undergoing long-term balancing selection is *x* ∈ (0,0.5]. The probability of observing *k* minor alleles out of *n* sampled alleles is then

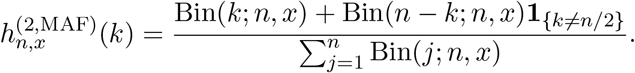

Hence, for an informative site *d* recombination units away from the presumed site under selection, the probability of observing *k* minor alleles out of *n* sampled alleles is

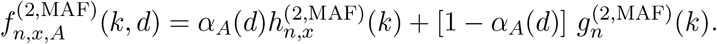

Similarly, when substitutions are not considered or are missing in the data (*i.e*., only observed minor alleles counts *k* = 1, 2,…, ⌊*n*/2⌋), the two mixing components can be normalized as

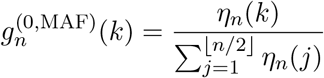

and

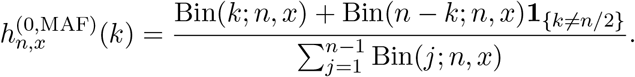

The probability of observing a polymorphic site with *k* minor alleles out of *n* sampled alleles is then

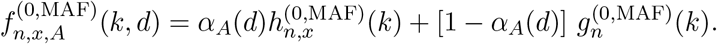

### Composite likelihood ratio tests based on the mixture models

Based on the probability distributions described for the five models, for each model *X* ∈ {“2”,“2,MAF”, “1”, “0”, “0,MAF”}, the composite likelihood of a genomic region with *L* informative sites under the null hypothesis of neutrality is

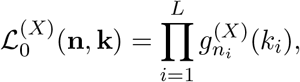

where **n** = [*n*_1_, *n*_2_,…, *n_L_*] and **k** = [*k*_1_, *k*_2_,…, *k_L_*] are the vectors of sample sizes and derived or minor allele counts, respectively, at the *L* informative sites in the genomic region. Recall that the probabilities of sampling a certain number of derived or minor alleles under our model depend on the sample sizes at informative sites, and because sample sizes often vary across the genome due to missing data in empirical studies, we make explicit the sample sizes across all informative sites in the vector **n**. Similarly, the composite likelihood under the alternative hypothesis of model *X* would be

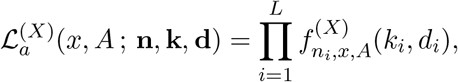

where **d** = [*d*_1_, *d*_2_,…, *d_L_*] is the vector of recombination distances between the test site and each of the *L* informative sites. This likelihood is maximized at

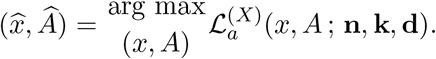

Hence, under model *X* ∈ {“2”,“2,MAF”, “1”, “0”, “0,MAF”}, the log composite likelihood ratio test statistic for the test site is

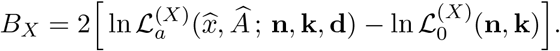

### Interpretation of estimated *A* and *x* parameters

The likelihood for the alternative model is maximized over the parameters *A* and *x*, where, in our formulation for bi-allelic balancing selection in the previous subsections, *x* represents the presumed equilibrium minor allele frequency, and *A* decides the rate of exponential decay for the probability of two sites being linked, which essentially describes the influence of balancing selection on neutral sites of varying distance away from the test site. After optimizing over this parameter space, the parameter values under the optimal likelihood, *Â* and 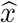, provide information on the nature of detected genomic footprints. The value of 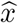 should reflect the enriched minor allele frequency across the region. Note that not all mechanisms for balancing selection will maintain the balanced alleles at fixed frequencies (Asmussen and Basnayake, 1990; Bergland et al., 2014), and so 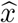 rather represents the value around which our model presumes the allele frequencies across the region are enriched. Therefore, we advise that caution be used when interpreting 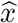 as the equilibrium frequencies without further information about the potential mechanisms that may have acted to maintain the polymorphisms.

Meanwhile, *Â* describes the rate of the exponential decay of the probability *α_A_*(*d*) = exp(−*Ad*) of the two loci being linked, and should intuitively be informative of the impact of balancing selection on nearby neutral sites. The smaller the *Â*, the wider the footprint would be, and likely the younger the balanced polymorphism. However, multi-locus balancing selection can also give rise to wide footprints (Barton and Navarro, 2002; Navarro and Barton, 2002; Tennessen, 2018), which could induce small *Â* values. Furthermore, a large *A* reduces the number of informative sites that yield meaningful likelihood ratios, and can thus also occur when data in the examined area fit the alternative model poorly. Therefore, we advise only comparing the *Â* values among regions with reasonably high composite likelihood ratios, and that caution be used when making inferences from these values as they do not map to an explicit evolutionary model.

## Results

### Performances on simulated data

We simulated 50 kilobase (kb) long sequences using SLiM3.2 (Haller and Messer, 2019), under the three-species demographic model (Figure S1) inspired by the demographic history of great apes (see *Methods*), and extensively evaluated the performances of all five *B* statistic variants. We also compared the *B* statistics to the summary statistics *β, β*\*, HKA, NCD2, and *β*^(2)^, which are respectively analogues to *B*_0_, *B*_0,MAF_, *B*_1_, *B*_2,MaF_, and *B*_2_, and to the likelihood statistics *T*_1_ and *T*_2_, which are respectively analogues to *B*_1_ and *B*_2_.

### Robust high power under varying window sizes

We first examined the robustness of the *B* statistics to overly large window sizes, under a scenario of strong heterozygote advantage (selective coefficient *s* = 0.01 with dominance coefficient *h* = 20) acting on a mutation that arose 7.5 × 10^4^ generations prior to sampling, with all sites flanking the selected locus evolving neutrally. Because `BetaScan` (Siewert and Voight, 2017, 2018) (which implements the standardized and nonstandardized *β*, *β*\*, and *β*^(2)^ statistics, among which we only consider the standardized) operates on windows of fixed physical length, we adopted window sizes of 1, 1.5, 2.5, 3, 5, 10, 15, 20, and 25 kb for all summary statistics and *B* statistics. The *T* statistics were applied on windows with matching expected numbers of informative sites. *Supplementary Note* 2 details the calculation for matching the number of informative sites to physical length of a genomic region.

To reduce potential stochastic fluctuations in the number of true positives when the false positive rate is controlled at a low level, we examined the area under a partial curve with no greater than a 5% false positive rate (hereafter referred to as “partial AUC”). As shown in Figure 2, under optimal window sizes for most other statistics, all variants of *B* statistics display substantial partial AUCs comparable to that of the respective *T* statistic variant, which has outperformed other equivalent summary statistics in most previous simulation studies (DeGiorgio et al., 2014; Siewert and Voight, 2017, 2018; Bitarello et al., 2018; Cheng and DeGiorgio, 2019). Most remarkably, as the window size increases, while all other statistics exhibit drastic decays in power, the powers of all variants of the *B* statistic only show minor decreases. In fact, when comparing the powers under 25 kb windows against those under optimal window sizes for each statistic, the powers of all statistics drop more than twice as much as *B*_1_ and *B*_2_ (Figure 2B). In comparison with each method’s optimal performance, most statistics (except all *B* statistics and *T*_2_, the model-based analog of *B*_2_) lose more than 80% of their optimal power under the largest window size examined (Figure 2C). Although *T*_2_ still retains considerably higher partial AUC compared to all other extant methods, it still decreases to a value substantially lower than that of *B*_2_. Such robustness of *B* statistics to large windows is reasonable and expected, because the probability distribution of allele frequencies at sites far enough from the test site will match the genome-wide SFS, thereby contributing little to the overall likelihood ratio.

**Figure 2:**
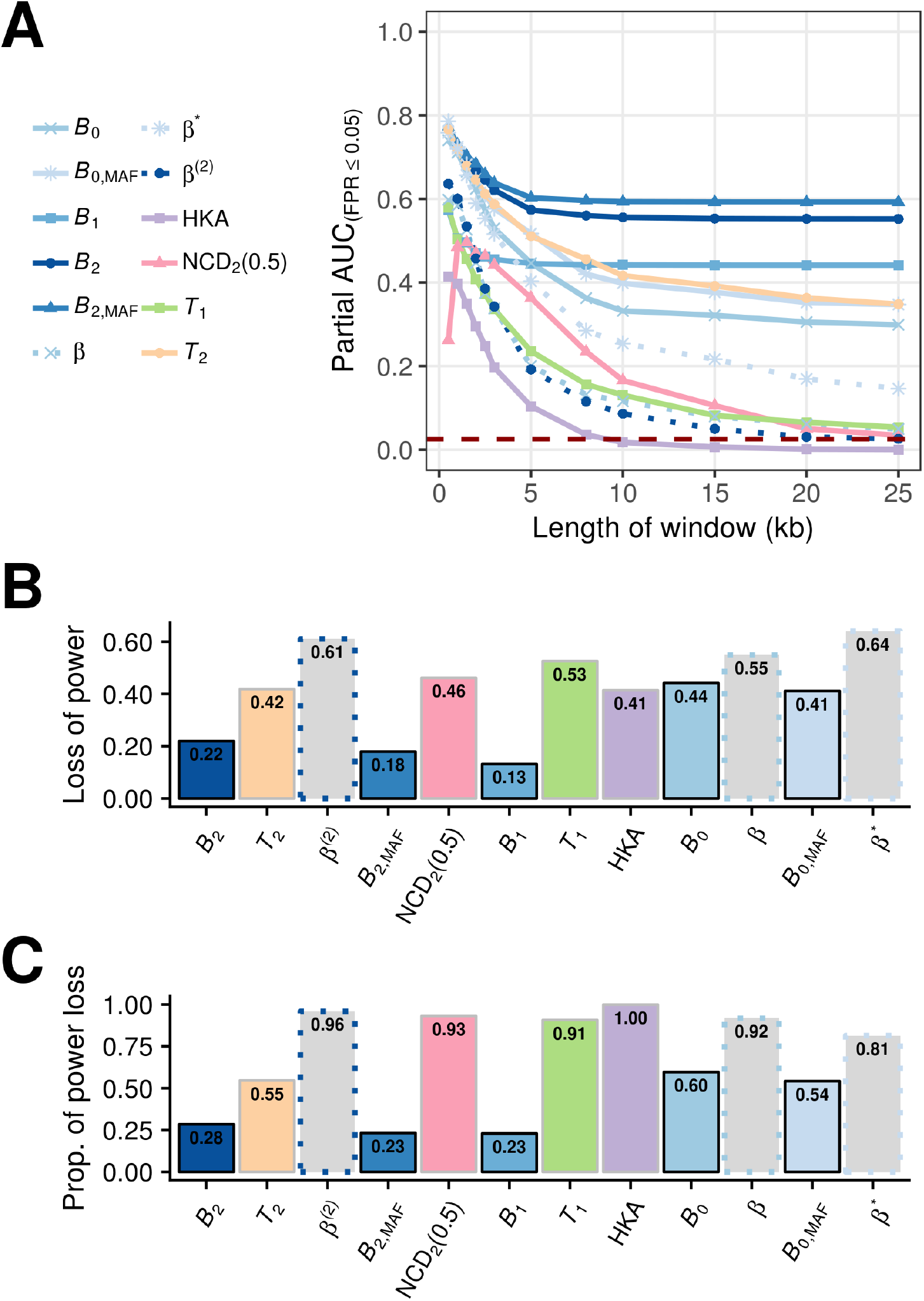
Partial area under the curve (AUC) conditioned on false positive rates (FPRs) less than or equal to 5% (defined such that the maximal value is 1) as a function of window size measured in kilobases (kb) for *B* statistics (varying shades of blue), *β* statistics (dotted line with varying shades of blue), *T*_2_ (orange), *T*_1_ (green), HKA (purple), and NCD_2_(0.5) (pink), under a scenario in which a mutation undergoing ancient balancing selection (selective coefficient *s* = 0.01 and dominance coefficient *h* = 20) arose 15 million years ago (assuming a generation time of 20 years). Statistics that consider the same input type share the same point shape. The dark red dashed line marks the level of partial AUC expected at the *y*=*x* line, or the baseline of randomly choosing between balancing selection and neutrality. (B) The amount of partial AUC lost, and (C) the proportion of the AUC loss as compared with the optimal value for each statistic when the window size increased from the optimum to 25 kb (*e.g*., largest evaluated).

Among all statistics evaluated, we found that those considering polymorphism data only (*i.e*., *B*_0_ variants and *β* variants) demonstrated relatively poor robustness to increases in window size. This result indicates that the detectable footprint of balancing selection in polymorphism data by itself may decay faster than other types of information, and that incorporating substitution data may help improve robustness to large windows.

Considering that the powers of all *B* statistics stabilize at a fixed level as the window size increases (Figure 2), we permit the *B* statistics to employ all informative sites on a chromosome. However, to reduce computational load, we only consider sites with mixing proportion *α_A_*(*d*) ≥ 10^−8^ for each value of *A* considered during optimization, which does not create discernible differences in performance from when all data are considered (Figure S2). However, to ensure that other methods still display considerable power for their comparisons, we applied the summary statistics with their optimal window sizes of one kb, and *T* statistics with numbers of informative sites expected in a one kb window (see *Methods*), unless otherwise stated.

### High power for detecting balancing selection of varying age and selective strength

Next, we explored the powers of *B* statistics when the selective strength *s*, equilibrium frequency (controlled by the dominance parameter *h*), and the age of balancing selection vary. Specifically, we examined scenarios where the selective coefficients were moderate (*s* = 0.01, Figures 3A, C, D, and E) or weak (*s* = 10^−3^, Figure 3B), and when the equilibrium frequency of the minor allele is 0.5 (*h* = 20, Figures 3A and B), 0.4 (*h* = 3, Figure 3C), 0.3 (*h* = 1.75, Figure 3D), or 0.2 (*h* = 1.33, Figure 3E). Across all scenarios considered, *T*_2_ and *β*\* show the highest power for old balancing selection. The best-performing *B* variants, *B*_2_ and *B*_2,MAF_, display high power as well, and are often comparable to that of the *β*^(2)^ statistic. The power of *B*_1_ is also similar to HKA, which is its summary statistic analogue. Furthermore, we noticed that *B* statistics exhibit superior power for younger balanced alleles, particularly when balancing selection is more recent than 2 × 10^5^ generations, and when the equilibrium frequency does not equal to 0.5 (Figure S3). For older selected polymorphisms, although several statistics outperform *B* statistics, it is important to point out that all previous methods were provided optimal window sizes, whereas *B* statistics were set to use all sites with considerable *α_A_*(*d*), under which they show lower power than when window sizes are optimized (Figure 2). This difference in performance between previous methods applied with their optimal window sizes and *B* statistics can also explain the seemingly inferior performance of the two Bo variants when compared with the analogous *β* statistics, as the *B*_0_ variants lose more power than other *B* variants when computed on extended windows. When applied with the same window size, however, Bo outperforms *β* by a large margin (Figure 2). Nevertheless, these results give us confidence that *B* statistics have generally high power to detect young and old balancing selection, even when adopting large windows.

**Figure 3:**
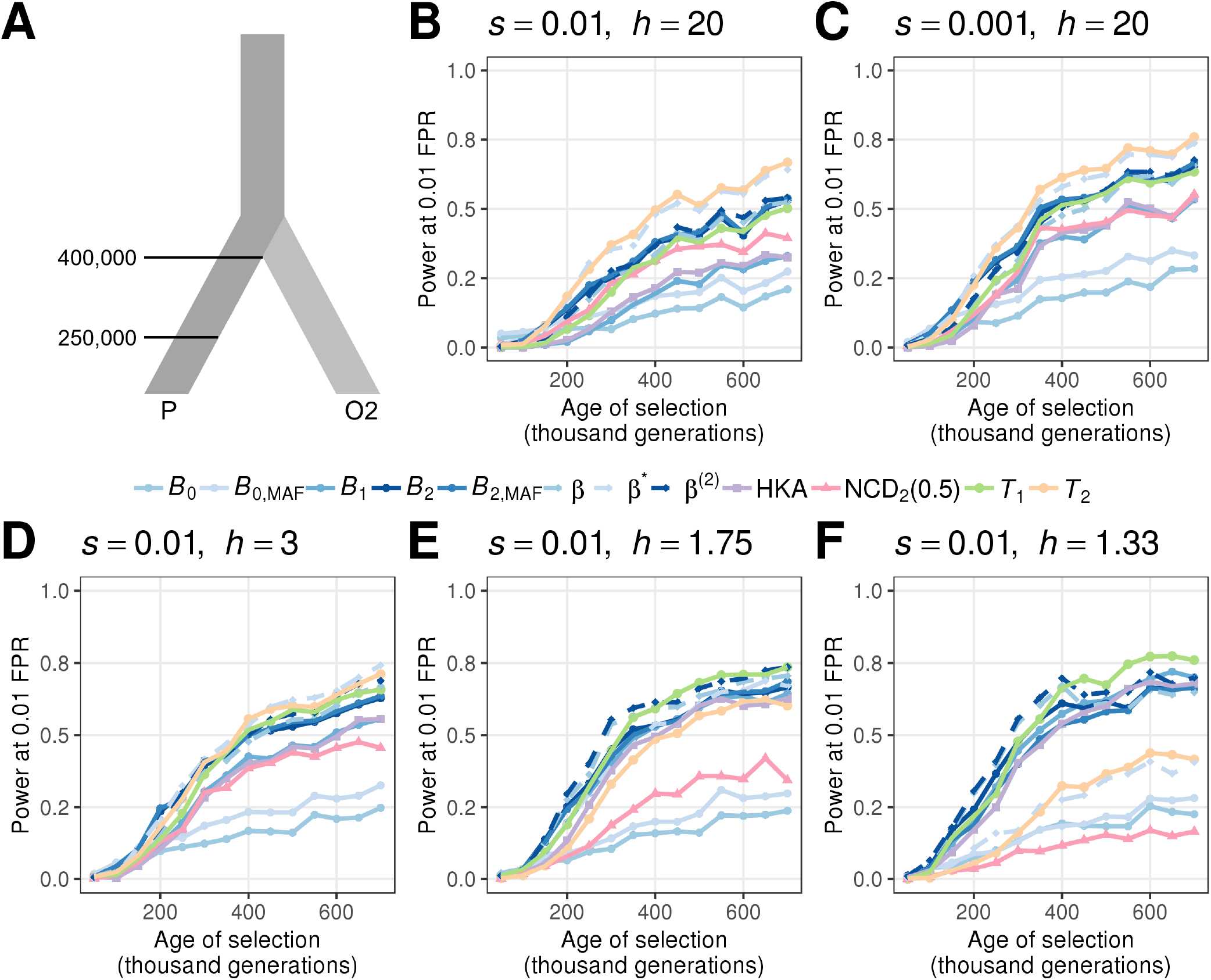
Ability to detect balancing selection for different heterozygote advantage scenarios. (A) Demographic model relating the ingroup (P) and outgroup (O2) populations, with one sample from O2 used as the outgroup sequence. (B-F) Powers at a 1% false positive rate (FPR) for each statistic as a function of age of the allele undergoing balancing selection for different selection (s) and dominance (h) coefficients. The scenarios considered are (B) *s* = 0.01 with *h* = 20, (C) *s* = 0.001 with *h* = 20, (D) *s* = 0.01 with *h* = 3, (E) *s* = 0.01 with *h* = 1.75, and (F) *s* = 0.01 with *h* = 1.33. Note that the equilibrium frequencies for panels D, E, and F are 0.4, 0.3, and 0.2, respectively.

### Robustness to recombination rate variation and elevated mutation rates

Despite their flexibility in window size and high power for detecting balancing selection, model-based methods, such as the *T* and *B* statistics, incorporate recombination distances in their inference framework, and can therefore be especially susceptible to potential inaccuracies in input recombination maps. Additionally, because many approaches for detecting balancing selection aim to identify genomic regions with increased genetic diversity, the elevation of mutation rates is also a common and potent confounding factor for detecting balancing selection (Charlesworth, 2006; Siewert and Voight, 2018; Cheng and DeGiorgio, 2019).

To test their robustness to inaccurate recombination rates, we applied *B* and *T* statistics on simulated sequences with uneven recombination maps (10^2^-fold fluctuations in recombination rates; see *Methods*). When the sequences evolve neutrally, neither approach is misled (Figure S4). When the fluctuation in recombination rate is even more drastic (*e.g*., 10^4^-fold instead of 10^2^), all methods tend to report fewer false signals than they would under a uniform map (Figure S5). This result suggests that the misleading effects of inaccurate recombination maps are limited.

To examine their performances under mutation rate variation, we next simulated a 10 kb mutational hotspot at the center of the 50 kb sequence with a mutation rate five times higher than original and surrounding rate *μ*, and applied each statistic with parameters derived from the original neutral replicates with constant mutation rate *μ* across the entire sequence. All methods exhibit considerable robustness against this regional increase of mutation rate (Figure S6).

We further considered an elevated mutation rate of 5*μ* across the entire 50 kb sequence, and re-examined the performance of each method. As expected, most statistics display substantially inflated proportions of false signals (Figures 4A and D). Among them, the *B*_2_ statistic reports the least proportion of false signals, followed by the *B*_1_ statistic. Meanwhile, at low false positive rates, *B*_2_ and *B*_2,MAF_ statistics report higher proportions of false signals than *T*_2_, their coalescence model-based analogue, whereas *B*_1_ outperformed *T*_1_. Additionally, all statistics that consider only polymorphism data, namely the *B*_0_, *B*_0,MAF_, *β*, and *β*\* statistics, are substantially misled. The *β*^(2)^ statistic, albeit taking substitutions into account, also displays surprisingly high proportions of false signals. Taken together, these results suggest that *B* statistics are reasonably robust against high mutation rates, relative to their analogous approaches, and that incorporating substitutions and polarized allele frequencies may further buttress the robustness.

**Figure 4:**
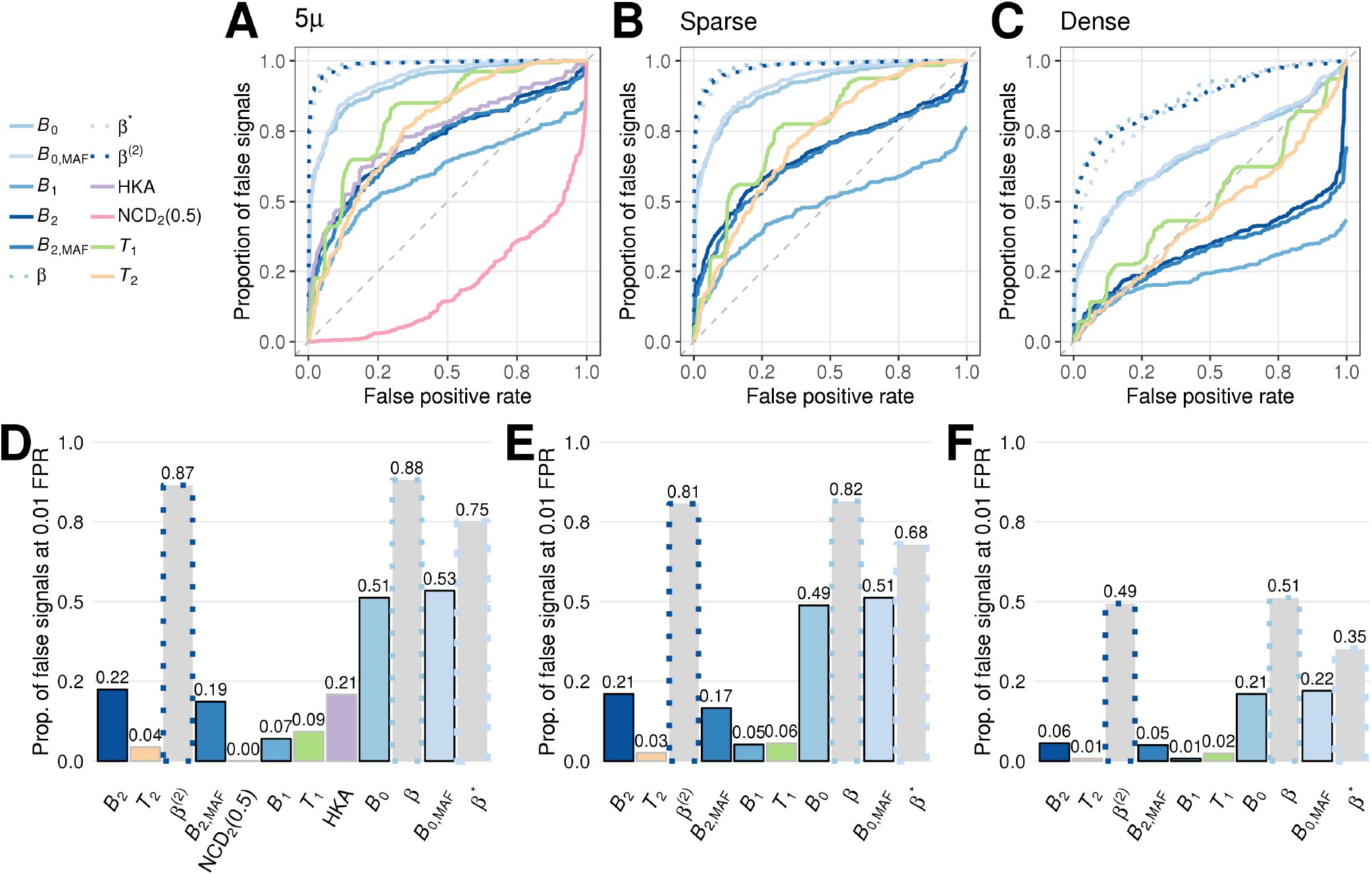
Robustness of each statistic evaluated when the mutation rate is elevated five-fold across an extended genomic region. (A-C) Proportions of false signals as a function of false positive rate (FPR) when (A) all test sites were considered to compute proportions of false signals and when (B, C) the number of test sites were down-sampled to match the number of sites expected on sequences with the original mutation rate *μ*. In panel B, five informative sites flanking either side of a test site considered were skipped (“sparse” down-sampling), whereas in panel C only the first 1200 informative sites were considered for calculating *T*, *B*_2_, and *B*_1_ statistics, and the first 240 polymorphic sites were considered for Bq and *β* statistics (“dense” down-sampling). (D-F) Proportions of false signals at a 1% FPR for each statistic for (D) all test sites, (E) sparse down-sampling, and (F) dense down-sampling. Bars for all *B* statistics are bordered by black lines, and all *β* statistics are bordered by gray lines in panels D-F. HKA and NCD_2_ (0.5) are applied with a fixed step size, and are therefore not considered when down-sampling in panels B, C, E, and F.

### Robust power under realistic demographic models

The influence of demographic history was the major motivation for *T* statistics to adopt the genome-wide SFS instead of the coalescence-based constant-size neutral model as the null hypothesis, despite that the latter being nested under the alternative model for balancing selection used by the *T* statistics. This trade-off has endowed *T* statistics with considerable robustness to population size changes (DeGiorgio et al., 2014; Cheng and DeGiorgio, 2019), but has also potentially compromised their robustness to large windows, as shown in *Robust high power under varying window sizes* subsection of the *Results*. For *B* statistics, however, because their null models both reflect the genome-wide SFS and are nested under the alternative models, they should exhibit considerable robustness to both oversized windows and demographic changes.

To evaluate their performances under recent population expansions and bottlenecks, we considered the demographic histories of contemporary European humans (Terhorst et al., 2017, CEU; Figure S7A) and bonobos (Prado-Martinez et al., 2013, Figure S8A; see details in *Methods*), respectively. The former have been extensively characterized (*e.g*., Lohmueller et al., 2009; Gravel et al., 2011; Terhorst et al., 2017), and therefore can reliably reflect the performance of each method under realistic scenarios. On the other hand, because we intend to apply the *B* statistics on bonobo genomic data, we are also interested in evaluating their performance under an inferred bonobo demographic model.

As previously described, we applied the *B* statistics with unlimited window sizes, whereas the other statistics were provided with smaller window sizes matching the theoretical size for a footprint of longterm balancing selection (see *Supplementary Note* 2). Despite being provided disadvantageous window sizes, *B* statistics still demonstrate comparable to, and often higher power than, current summary statistic approaches, both under the human (Figure S7) and the bonobo (Figure S8) demographic models. Although *T*_2_ has higher power than the *B* statistics, we note that the *T* statistics were operating with optimal window sizes, whereas the window sizes for *B* statistics are identified across a parameter range. When *B*_1_ and *B*_2_ are applied with identical window sizes as *T*_1_ and *T*_2_ (Figures S9 and S10), the margins between their performances are no longer substantial. Additionally, we noticed that most statistics tend to have higher power for sequences evolving under the bonobo demographic history than under that of the Europeans (notice that the *y*-axes in Figures S7 and S8 have different scales).

### Robust power under varying mutation rates across target and outgroup species

In addition to temporally-varying population sizes, differing mutation rates between closely-related species may also affect the performance of the coalescence-based *T* statistics, as they assume a uniform neutral mutation rate along the genealogy relating the lineages from the ingroup and outgroup species. Among great apes, for example, accumulating evidence suggests that humans have substantially lower mutation rates than other great apes (as reviewed by Scally and Durbin, 2012).

To examine the behavior of each method when mutation rates of the target and outgroup species differ, we simulated a two-species demographic history, with the target and outgroup species respectively evolving at neutral rates *μ* = 1.2 × 10^−8^ and *μ* = 2.5 × 10^−8^ mutations per site per generation (see *Methods* for details). We introduced an adaptive mutation evolving under balancing selection at varying time points prior to sampling along this demographic history, and examined the power of each statistic to detect balancing selection across a diverse array of selection parameters (Figure S11).

Across all six combinations of selection parameters considered, we observe similar trends for each statistic when compared with simulations under the constant population size (Figure 3) and CEU (Figure S7) demographic histories evolving with a constant neutral mutation rate. The *T*_2_ statistic performs the best when *s* = 0.01 with *h* = 20 (Figure S11A), under which the equilibrium frequency is closest to 0.5 and when heterozygotes are most advantageous. As the selective advantage *hs* and equilibrium frequency decrease, the margin between the powers of *T*_2_ and *B*_2_ shrinks, and even reverses for all scenarios with small dominance *h* (Figures S11C-F). Furthermore, methods based solely on polymorphism and substitution calls (*i.e*., *T*_1_, *B*_1_, and HKA) show improvements in power as the equilibrium frequency decreases, and some even outperform most of the other statistics (Figures S11D and E). Statistics that ignore substitutions (*i.e*., *B*_0_, *B*_0,MAF_, *β*, and *β*\*), on the other hand, perform especially well for recent balancing selection with high heterozygote advantage (large *hs*; Figures S11A and B). As the balanced alleles reach their equilibrium frequencies sooner when the selective advantage of heterozygotes (*i.e*., *hs*) is high, sequences with mutations of higher *hs* would have older footprints than those with mutations introduced at the same time but with lower *hs*. In this respect, it is understandable that *B*_0_ and *β* variants outperform others only for selection with large *hs* that are introduced within 150,000 generations prior to sampling.

Based on this two-species model with diverging mutation rates, we further integrated changes in population size of the target species in accordance with the demographic history of the CEU (Terhorst et al., 2017, Figure S12). From the four sets of selection parameters tested, we found that most methods exhibit lower power compared with those under constant population sizes (Figure S11). This result is consistent with the lower powers under simulations with a constant mutation rate when the target population size evolves under the CEU demographic history (Figure S7) compared with the setting in which the target evolves with constant size (Figure 3). Despite their lower powers in general, we still observe similar relative performances across statistics, with *T*_1_ and *B*_1_ exhibiting higher powers when the heterozygote advantage *hs* is small. Moreover, we found that *B*_2,MAF_ shows superior power to *B*_2_.

### Re-examining long-term balancing selection in human populations

We applied *B*_2_ on contemporary European (Europeans in Utah; CEU, Figure S14) and west African (Yoruban; YRI, Figure S13) human populations from the 1000 Genomes Project dataset (The 1000 Genomes Project Consortium, 2015) (see *Methods*) to re-examine the footprints of long-term balancing selection, which previous studies (DeGiorgio et al., 2014; Siewert and Voight, 2017) have provided cases for reference. The most outstanding candidates in both scans localize in the HLA-D region (human leukocyte antigen, also known as major histo-compatibility [MHC] Class II region) (Figures S15 and S16), agreeing with previous findings (Sanchez-Mazas, 2007; Leffler et al., 2013; DeGiorgio et al., 2014; Teixeira et al., 2015; Siewert and Voight, 2017; Meyer et al., 2017; Bitarello et al., 2018). Within the HLA-D region, the *B*_2_ scores computed for both populations show extraordinary peaks around *HLA-DQ* and *HLA-DP* gene clusters, although CEU (Figure S15) scores remarkably higher on *HLA-DP* genes than YRI (Figure S16). Echoing the critical roles of HLA-D genes in adaptive immunity, the gene *ERAP2* exhibits extraordinary scores in both populations (Figures S17 and S18). This gene has been reported in past studies of balancing selection in humans (Andrés et al., 2009, 2010; Bitarello et al., 2018), and Andrés et al. (2010) demonstrated that its splicing variants can alter the level of MHC-I presentation on *B* cells. Additionally, we also observed high *B*_2_ scores on *CADM2* (Figures S19 and S20) and *WFS1* (Figures S21 and S22), on which Siewert and Voight (2017) characterized potential non-synonymous mutations segregating in both populations.

In addition to these previously characterized candidates, both scans display extreme *B*_2_ scores on another two top-ranking regions in the *T*_2_ scans by DeGiorgio et al. (2014): the *STPG2* gene (formerly named *C4orf37*; Figures S23 and S24) and the *CCDC169-SOHLH2* (formerly named *C13orf38-SOHLH2*; Figures S25 and S26) region, with *STPG2* particularly more outstanding in the scan of YRI than in CEU. Intriguingly, both these genes are associated with gametes. The *STPG2* gene encodes sperm-tail PG-rich repeat-containing protein 2, which, despite the paucity of literature that describes its function, is found in sperm (Uhlen et al., 2015). The high-scoring region on this gene harbors a number of tissue-specific eQTLs for its expression, especially in brain and reproductive tissues (GTEx Consortium, 2017). The *SOHLH2* gene, on the other hand, encodes the transcription factor Spermatogenesis and Oogenesis Specific Basic Helix-Loop-Helix-containing protein 2, which plays important roles in both spermatogenesis and oogenesis (Toyoda et al., 2009; Suzuki et al., 2012). We observed drastically elevated *B*_2_ scores (Figure S25) across an extended region upstream of *SOHLH2* that covers the naturally occurring *CCDC169-SOHLH2*readthrough transcript (as introduced in RefSeq database; O’Leary et al., 2015). Similar to *STPG2*, this region also features numerous eQTLs for the expression of *SOHLH2*, especially in endocrine glands, brain, and reproductive tissues (GTEx Consortium, 2017).

Other regions with outstanding peaks shared by both scans include the genes *CPE* (Figures S27 and S28) and *MYOM2* (Figures S29 and S30). *CPE* encodes carboxypeptidase E, a key enzyme for synthesizing peptide hormones such as insulin and oxytocin, and its mutant mice strain (*Cpe*^fat^) exhibits endocrinic disorders such as obesity and infertility (Naggert et al., 1995). *MYOM2* encodes endosacromeric cytoskeleton M-protein 2, which serves as a structural component of muscle tissues (van der Ven et al., 1999). Both genes harbor eQTLs reported by GTEx Consortium (2017) around the high-scoring regions.

### Probing for footprints of balancing selection in bonobo genomes

We further applied the *B*_2_ statistic on the variant calls of 13 bonobos (Prado-Martinez et al., 2013) lifted-over to human genome assembly GRCh38/hg38. Only bi-allelic single nucleotide polymorphisms (SNPs) were considered, and substitutions were called using bonobo panPan2 reference sequence (Prüfer et al., 2012), with the human sequence as the ancestral state. Stringent filters were applied to remove repetitive regions and regions with poor mappability (see *Methods*), with the resulting mask unintendedly removing the majority of the MHC locus as well. We observed many genomic regions with outstanding *B*_2_ scores (Figure 5), which include both the *MHC-DQ* genes and a few novel candidates.

**Figure 5:**
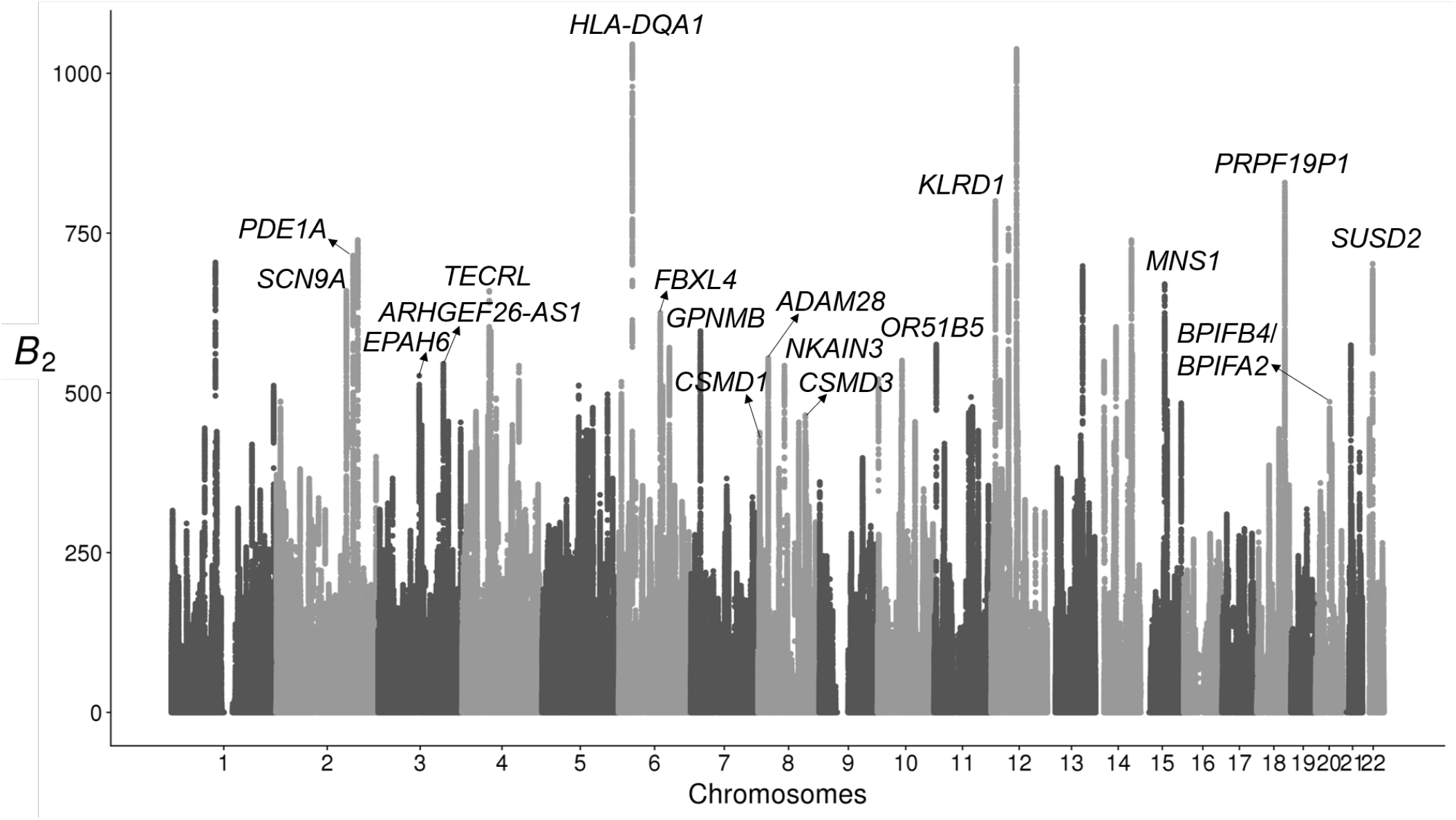
Manhattan plot displaying *B*_2_ scores across the 22 human autosomes for which the bonobo genomic data were mapped, with the top candidates annotated. Outstanding peaks (*B*_2_ > 600) without annotations do not have neighboring protein-coding regions within a 500 kb radius.

Among the outstanding peaks, the top one covers the *HLA-DQA1* and *HLA-DQB1* genes, which are among the remainder of the MHC region that survived filtering (including many filtered gaps). Nonetheless, both the *MHC-DQ* and *MHC-DR* clusters exhibit high likelihood ratio scores (Figure S31A). Such high scores can be explained both by the elevated proportion of polymorphic sites, 0.754 as compared with the genome-wide proportion of 0.237 (Figure S31B), as well as the enrichment of polymorphic sites with moderate minor allele frequencies (Figure S31C). Furthermore, the region exhibits a multimodal SFS, which may correlate to the multiple *B*_2_ peaks observed in the region.

In addition to the *HLA-DQ* genes, *KLRD1* also presents prominent *B*_2_ scores (Figure 6) on its first intron. This gene expresses a natural killer (NK) cell surface antigen, also known as CD94, and plays a pivotal role in viral defense. Unlike the region covering *MHC-DQ* genes, the minor allele frequencies at polymorphic sites around the *KLRD1* region are clearly enriched near a frequency of 0.45, instead of the multimodal distribution observed around the *MHC-DQ* genes. Another interesting candidate is the pain receptor gene *SCN9A* (Figure S32), on which the highest scores overlap with the transcript of its anti-sense RNA gene that regulates its expression. Instead of enriching toward a single value, the minor allele frequencies at the polymorphic sites across the region are dispersed, with at least two modes (approximate minor allele frequencies of 0.25 and 0.40). This finding may correlate with the multiple peaks identified around this region, which may be sensible given the large number of exons covered. Other notable candidates include two developmental genes *CSMD1*and *CSMD3* (Figures S33 and S34, respectively), as well as *PDE1A* (Figure S35), which encodes a pivotal enzyme in cellular Ca^2+^- and cyclic nucleotide signalling, and plays important roles in both neurodevelopment and sperm functionality. We also found other high-scoring regions associated with innate immunity, such as the gene *GPNMB* (Figure S36) and the intergenic region between *BPIFB4* and *BPIFA2* (Figure S37).

**Figure 6:**
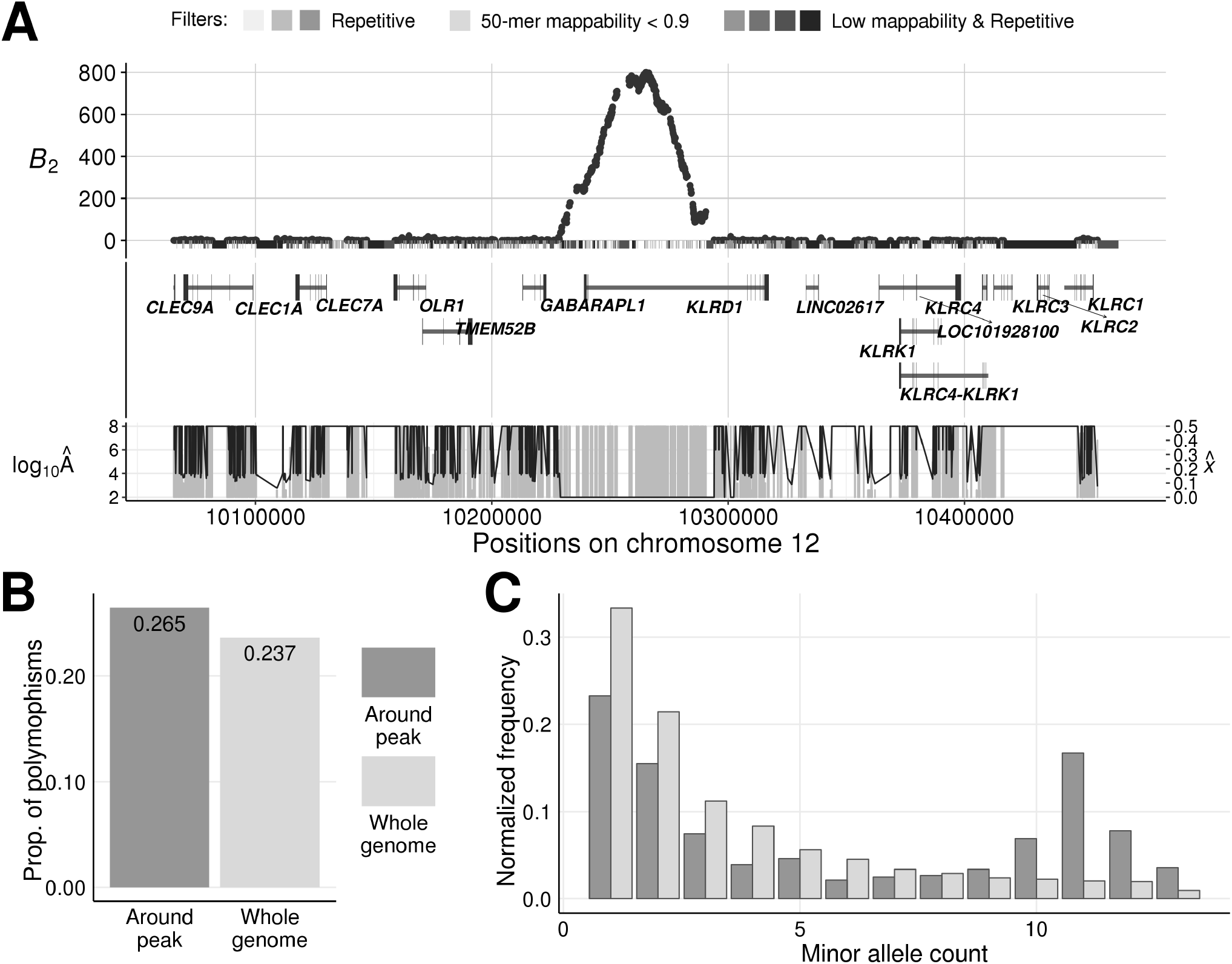
Evidence of balancing selection on *KLRD1*. (A) *B*_2_ scores across the 400 kb genomic region on chromosome 12 centered on the peak. The gray bars directly under the *B*_2_ scores represent the masked regions, as well as the features in these regions. The darker the shade, the greater number of types of repetitive sequences (*e.g*., RepeatMasker mask, segmental duplication, simple repeats, or interrupted repeats) overlapping the region. Vertical gray bars below display the estimated equilibrium minor allele frequency 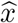 for each maximum likelihood ratio *B*_2_, and the black line traces the value for the respective inferred footprint size log_10_(*Â*). (B) Proportion of informative sites that are polymorphic in the 400 kb region centered on the peak compared with the whole-genome average. (C) Minor allele frequency distribution in the 400 kb region centered on the peak compared with the whole-genome average.

## Discussion

In this study, we introduced a novel set of composite likelihood ratio statistics—*B*_2_, *B*_2,MAF_, *B*_1_, *B*_0_, and *B*_0,MAF_—to robustly detect footprints of balancing selection with high power and flexibility. The *B* statistics are based on a mixture model creating a proper nested likelihood ratio test, which helps them overcome the common susceptibility to oversized windows held by current methods. We have extensively evaluated their performances on simulated data compared with current state-of-the-art methods, and have demonstrated the superior properties of the *B* statistics under various scenarios. We re-examined balancing selection in human populations (The 1000 Genomes Project Consortium, 2015), and recovered well-established candidates including the HLA-D genes and *ERAP2*. We further applied *B*_2_ onto the genomic data of bonobos (Prado-Martinez et al., 2013), and uncovered not only the *HLA-DQA1* /*HLA-DQB1* gene cluster, but also intriguing candidates that are involved in innate immunity.

### Evaluating the performance of *B* statistics through simulations

In our simulation study, the *B* statistics showed remarkable robustness to large window sizes, with only minor decays in power under oversized windows, whereas other methods exhibited large declines in power. Moreover, even when considering all data available as input (*i.e*., the most disadvantageous window size) all variants of *B* statistics still exhibit comparable power to extant methods and displayed satisfactory performance across varying types and strengths of balancing selection. Under scenarios with confounding factors, such as high mutation rate and non-equilibrium demographic history, the *B* statistics demonstrated satisfactory robustness as well.

The robustness against varying window sizes is of particular interest in this study, not only because it ensures high power under large windows, but it also allows the statistics to augment the size of genomic regions from which they make meaningful inferences. This flexibility grants a key advantage over previous methods that require the window size to be fixed throughout the scan in order to yield comparable results across the genome. In particular, because many factors (such as recombination rates) can influence the footprint size of balancing selection, it is not ideal to adopt a fixed window size for a whole-genome scan based on a uniform population-scaled recombination rate, and *B* statistics naturally accommodate such variability across the genome.

Admittedly, in practice, as the genomic region considered in the tests expands, non-neutral sites will inevitably be included. This indeed violates our assumption that the test locus is surrounded by neutral sites only. Nonetheless, because both positive and purifying selection reduce the presentation of sites with intermediate frequencies (Tajima, 1989; Braverman et al., 1995; Fay and Wu, 2000; Bamshad and Wooding, 2003), their effect on the SFS is in general opposite to the features expected from balancing selection. This suggests that including such sites in the window is unlikely to hamper the power to detect balancing selection. Meanwhile, when multiple sites in the considered region undergo balancing selection, the pattern of polymorphisms across the region will indeed differ from that in regions with a single selected locus. We will discuss the effects of such multi-locus balancing selection in the subsequent subsection *Performance of single-locus methods on multi-locus balancing selection*.

One important consideration is that, so far our simulation study (as well as previous ones by DeGiorgio et al., 2014; Bitarello et al., 2018; Siewert and Voight, 2018) only evaluates the method performance in the context of single-locus heterozygote advantage. For many other balancing selection mechanisms, such as negative frequency-dependent selection (Asmussen and Basnayake, 1990) and periodic environmental fluctuations (Bergland et al., 2014), a stable equilibrium cannot be guaranteed (Cockerham et al., 1972; Asmussen and Feldman, 1977; Ginzburg, 1977). In non-overdominance settings for which particular equilibrium frequencies indeed exist, the balanced alleles are still maintained near these fixed frequencies, thereby satisfying the general assumptions of the statistical models underlying our *B* statistics. Moreover, when such intrinsic equilibrium frequencies do not exist, allele frequencies may still fluctuate around some mean values. Even if such mean values are unattainable, there will still persist an enrichment of sites with intermediate frequencies, thereby presenting characteristic footprints of balancing selection. We therefore believe that our mixture model framework should still have high power to detect footprints of non-overdominance balancing selection, and that overall, our results have comprehensively characterized the promising performance of the *B* statistics.

### Confounding effects of mutation rate or recombination rate variation

In our simulation study, sequences with a central 10 kb mutational hotspot did not mislead methods as much as those with the mutation rate elevated across the entire sequence (Figure S6). This result may seem counter-intuitive at first, as a smaller region of increased mutation rate may better resemble the footprints of long-term balancing selection. However, upon a closer examination of the site frequency spectra and proportions of polymorphic sites (Figure S38), sequences with an extended region of high mutation rate exhibit a greater departure in these features under scenarios with no elevated mutation rate than for scenarios with a central mutational hotspot. Specifically, these sequences have more sites with high derived allele frequencies and a higher proportion of polymorphic sites overall (Figure S38B), likely resulting from the recurrent mutation on sites that were originally substitutions. The increase is also more profound on sites with high derived allele frequency. For example, the proportions of sites with derived allele frequency of 0.96 increased by almost two-fold from approximately 0.00104 to 0.00190, and the proportions of sites with derived allele frequency of 0.98 increased by almost three-fold from 0.00105 to 0.00273. By contrast, the difference in scale between the proportions of polymorphisms (0.182 versus 0.189) is minor. The larger fold-change in the proportions of high-frequency polymorphisms *(i.e*., sites with *k* = *n* − 1, *n* − 2, and *n* − 3 derived alleles) relative to that of substitutions (*k* = *n* derived alleles) could explain the more profound inflation in power for the statistics relying only on information at polymorphic sites. Similarly, after folding the SFS, the large changes in the proportions of low-frequency alleles were substantially mitigated, echoing the superior performance of *B*_2,MAF_ and *β* relative to their unfolded counterparts.

Another unexpected result from the simulations of elevated mutation rate is the drastic inflation of false signals reported by *β* statistics (Figure 4), which can also be observed in the non-standardized *β* statistics (Figure S39). Although Siewert and Voight (2018) tested their power to detect balancing selection under high mutation rate, it was unexplored whether their *β* statistics would mis-classify highly mutable neutral sequences as those undergoing balancing selection, and our results show that they could be easily misled. However, we further found that the performances of the standardized *β* statistics largely improve when provided with the correct mutation rate and divergence time (Figure S39B). This result partly confirms the superiority of standardized *β* statistics over the unstandardized ones. It also suggests that *β* statistics are considerably susceptible to the confounding effect of mutation rate elevation, and that their performance relies highly on the accuracy of the provided mutation rate. Instead of using a constant mutation rate for the entire scan, we propose that providing locally-inferred population-scaled mutation rates *θ* may help improve the robustness of *β* statistics. Indeed, when we instead estimate *θ* using the mean pairwise sequence difference 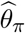 (Tajima, 1983) for each replicate and provided `BetaScan` the respective inferred value as the *θ* parameter, the standardized statistics no longer report as many false signals (Figure S39C).

In contrast to mutation rate variation, all statistics are robust to recombination rate variation, with *B*_0_ and *B*_0,MAF_ reporting substantially fewer false signals than the others (Figure S4). This robustness to recombination rate variation may be explained by the high similarity in the SFS and proportion of polymorphic sites to sequences evolving under a uniform recombination rate (Figure S40).

### Effect of multiple testing on sequences with high mutation rates

Because *B*, *T*, and *β* statistics are computed on every informative site, as suggested by Cheng and DeGiorgio (2019), multiple-testing can account for some inflation in their powers because sequences with a higher mutation rate will have a greater number of informative sites. To evaluate the effect of multiple testing for sequences with high mutation rates, we down-sampled the test sites (see *Methods*) such that the number of test scores being computed approximately matches that under the original mutation rate *μ*. Although all statistics show varying levels of improvements in performance (Figures 4B, C, E, and F), some still report high proportions of false signals, especially all *β* statistics and *B*_0,MAF_. That is, multiple-testing cannot account for all the factors that drive these statistics to mis-identify features of elevated mutation rates as footprints of balancing selection. This result corresponds to the fact that both the SFS and the density of polymorphic sites are altered under scenarios with extended regions of elevated mutation rate (Figure S38), likely due to recurrent mutation.

Furthermore, we observed that both before and after down-sampling, the *T* statistics report fewer false signals than their respective *B* statistic analogues. One potential factor behind their marginally superior performance may be that *T* statistics perform tests on fixed numbers of informative sites, instead of genomic regions measured by physical lengths (as did *B* statistics and the summary statistics). For *T* statistics, the size of the genomic region covered by the same number of informative sites would be much narrower under rapidly mutating sequences than in sequences with the original mutation rate. This means that the resulting *T* scores in either scenario are reflective of the levels of variation for sequences with drastically different lengths. To account for this factor, we provide *B*_1_ and *B*_2_ with informative site-based windows identical to that of *T* statistics and re-examined their performances (Figure S41). After matching the windows, *B*_1_ and *B*_2_ variants in turn display higher robustness than *T*_1_ and *T*_2_ to elevated mutation rates, suggesting that *B* statistics are at least comparably robust to *T* statistics. Meanwhile, we also matched the window size for B0 variants and *β* to gauge the effect of adopting large windows on the proportions of false signals from *B*_0_ variants. When *B*_0_ scans the sequences with one kb windows, though there is an increase in the resulting number of false signals (Figure S41A), at a 1% false positive rate the proportions of false signals for the two *B*_0_ variants only increase by less than 0.1, and are still substantially lower than that of *β* and *β*\* (Figure S41B).

### Comparing the *B* statistics with the *T* statistics

Because the *T* statistics of DeGiorgio et al. (2014) have previously been the only model-based approach for the detection of long-term balancing selection from polymorphism data in a single species, the comparisons between the model-based *B* and *T* approaches is particularly intriguing for researchers with empirical data suitable for the application of either. The *T* statistics are based on an explicit coalescent model (Hudson and Kaplan, 1988; Kaplan et al., 1988), and have been shown to have superior power to a number of other methods in previous studies (DeGiorgio et al., 2014; Siewert and Voight, 2017, 2018; Bitarello et al., 2018; Cheng and DeGiorgio, 2019), consistent with our simulation results. The *B* statistics, on the other hand, employ a mixture model, where the component modeling balancing selection is not based on an explicit evolutionary model, but nevertheless shows impressive performance on simulated data, as the shape of the distribution of allele frequencies is similar to what might be expected under balancing selection. The often superior performances of both approaches over summary statistics is understandable, as both utilize the genomic spatial distribution of genetic diversity in their inferences.

However, within the *T* statistic framework, the model for the null hypothesis (neutrality) is not nested in the alternative hypothesis (balancing selection). Although the *T*_1_ and *T*_2_ statistics could have adopted nested models by employing the standard neutral coalescent as the model for the null hypothesis, doing so would increase susceptibility to demographic factors, which can also alter the genome-wide SFS. To better account for these factors, DeGiorgio et al. (2014) instead employed the genome-wide distribution of genetic variation to compute probabilities under the null hypothesis of neutrality. This explains the substantial decay in power for both *T* statistics as the window size increases (Figure 2), as well as its robust performance under varying sized demographic models (DeGiorgio et al., 2014; Cheng and DeGiorgio, 2019, Figures S7 and S8). In contrast to the *T* statistics, the null model for *B* statistics (which also employs the genome-wide SFS) is nested within the alternative, due to their mixture model framework. This feature mitigates the biases introduced by sites far from the test site, while simulataneously accounting for demographic factors. Consequently, the *B* statistics display robust performance under oversized windows and realistic demographic models in our simulations (Figures 2, S7, and S8).

Another advantage of the *B* statistics over the *T* statistic approach, especially for *B*_2_ compared with *T*_2_, is the computational load. Because the probability distribution of allele frequencies under the Kaplan-Darden-Hudson (Kaplan et al., 1988) model is difficult to compute, the *T*_2_ statistic relies on previously-generated sets of simulated site frequency spectra over a grid of equilibrium frequencies *x* ∈ {0.05,0.10,…, 0.95} for each distinct sample size *n* and recombination distance *d*. Generation of such frequency spectra is computationally intensive, and the load increases substantially with the increase in sample size, thereby limiting the application of *T*_2_ to datasets with larger sample sizes. However, this is not a limitation of *B*_2_, as the SFS under balancing selection is determined simply as a mixture of the given genome-wide distribution of allele frequencies and a statistical distribution with closed-form solutions whose computational cost is minor, and only increases linearly with the sample size. Moreover, the rapid computation of this spectrum permits a finer grid of equilibrium frequencies *x* to be interrogated.

### Considering multi-allelic or multi-locus balancing selection

Both model-based approaches employed by the *T* and *B* statistics assume that balancing selection acts on a single bi-allelic locus. Whereas this case may be the most intuitive and simplistic scenario to model and simulate, many well-established empirical examples of balancing selection—such as the MHC locus in animals (Wills, 1991; Hedrick, 2002), the ABO blood group in primates (Saitou and Yamamoto, 1997; Fumagalli et al., 2009; Ségurel et al., 2012; Leffler et al., 2013), and the plant self-incompatibility locus (Charlesworth et al., 2000)—feature multiple alleles balanced across an extended genomic region. It therefore brings into question how these methods perform on genomic regions evolving under multi-allelic or multi-locus balancing selection, and whether current frameworks can be extended to consider these more complicated cases of balancing selection.

### Extending mixture models to account for multi-allelic balancing selection

There exist theoretical models of multi-allelic balancing selection based on the coalescent (Hey, 1991; Muirhead and Wakeley, 2009). However, possibly due to computational constraints, such models have not been implemented within a likelihood-ratio framework for detecting the footprints they characterize. Here, instead of following DeGiorgio et al. (2014) to compute the densities of polymorphisms and substitutions or to approximate the SFS using simulations under an explicit coalescent model, our mixture models can be readily extended to account for multi-allelic balancing selection at a single locus without the extensive computational burden of coalescent-based approaches that integrate selection. Specifically, we consider samples with multiple balanced alleles as following multinomial distributions (see *Supplementary Note 1*), and henceforth use the mixture models to approximate the SFS at bi-allelic neutral sites that are linked to a selected locus with *m* ∈ {2,3,4,…} balanced allelic classes. This extension is also implemented in our `BalLeRMix` software, with the special case of *m* = 2 reducing to the model introduced in the *Theory* section.

To simulate single-locus multi-allelic balancing selection, we employed SLiM version 3.3, which can simultaneously incorporate the four standard nucleotides of DNA, and thus allows these distinct nucleotides to coexist at the same site. We introduced two, three, or four distinct mutations with fitness parameters *s* = 0.001 and *h* = 20 in each simulated replicate 500,000 generations in the past to examine the relative performances of *T*, bi-allelic *B*, and multi-allelic *B* statistics. Under this fitness scheme, the equilibrium frequencies when two, three, or four alleles are balanced in the population are approximately (1/2,1/2), (1/3,1/3,1/3), or (1/4,1/4,1/4,1/4), respectively (see *Methods* for details). As the number of balanced alleles assumed by *B* statistics (*i.e*., parameter *m*) increases, the powers of *B* statistics barely change when two (Figures S42A-C) overdominant mutations are introduced. When more than two overdomiant alleles are balanced in the population, it is remarkable that *B* statistics with *m* set to three or four (Figures S42E and F, respectively) outperform those with *m* = 2 (Figure S42D). Furthermore, we also observe that the optimal equilibrium minor allele frequencies reported by the *B* statistics match well with the true equilibrium frequencies in the simulated replicates (Figure S43).

To further dissect the relative performances of *B* statistics (with *m* = 4), we also applied other statistics with their optimal window sizes on these simulated sequences (Figure S44). As the number of balanced alleles increases, each statistic demonstrated improvements in their power. Furthermore, the *B*_1_, *B*_2_ and *B*_2,MAF_ statistics outperform their respective *T* -or summary-statistic analogs under all three scenarios considered.

Taken together, these results suggest that the multi-allelic *B* statistics can substantially improve the detection power for balancing selection with more than two balanced alleles. Moreover, *B* statistics with larger *m* parameters, the presumed number of balanced alleles, are downward compatible with population samples carrying fewer than *m* balanced alleles, as the presumed equilibrium minor allele frequencies of the extra allelic classes would be optimized close to zero (see Figure S43).

### Performance of single-locus methods on multi-locus balancing selection

Similar to multi-allelic balancing selection, despite previous theoretical work to model or simulate multilocus balancing selection (Navarro and Barton, 2002; Barton and Navarro, 2002; Tennessen, 2018), no detection approach has yet been developed accordingly. Meanwhile, neither model-based detection framework underlying the *T* statistics nor the *B* statistics can address these cases without jointly accounting for allelic combinations at multiple informative sites as the target of selection. Therefore, without shifting the paradigm to consider such site-to-site combinations so as to accurately locate the set of neighboring selected loci, one can still examine the performance of extant balancing selection approaches for locating genomic regions containing more than one locus under balancing selection.

To this end, we tested the simplest case with two nearby loci carrying independent overdominant alleles (see *Methods*). To ensure individuals heterozygous at both loci are as advantageous as in the single-locus balancing selection simulations with *s* = 0.001 and *h* = 20 (Figures S45A and B), we set the selective coefficients of both overdominant mutations to *s* = 0.0005. That is, a two-locus genotype that is heterozygous at each of the loci would have fitness approximately equal to 1 + 2*hs* = 1.2. Despite this adjustment, we observed that all statistics show drastic improvements in their powers (Figure S45C and D), with the lowest power among them of 0.8 (Figure S45D). This result suggests that multi-locus balancing selection can potentially create more-prominent footprints compared with single-locus balancing selection. To further gauge the extent to which the additional selected locus can boost detection power, we simulated sequences with two nearby loci each evolving under *s* = 10^−5^ and *h* = 20, such that the selective coefficient *s* is two orders of magnitude smaller than that of the mutations introduced in the sequences evolving under single-locus balancing selection (Figures S45A and B). Remarkably, all methods still exhibit substantially higher powers for sequences with two nearby loci with weakly-advantageous (*s* = 10^−5^) alleles undergoing balancing selection (Figures S45E and F).

The higher powers observed for simulated multi-locus balancing selection scenarios is understandable, as Tennessen (2018) demonstrated that two non-interacting neighboring loci tend to reinforce the maintenance of polymorphisms when both are independently subjected to balancing selection. However, multi-locus balancing selection can also be achieved by epistasis (Barton and Navarro, 2002; Navarro and Barton, 2002), whereby the fitness effect of one locus is contingent on the allelic state of another locus, and has been shown by a growing body of empirical studies to be pervasive in the genome (as reviewed by Shao et al., 2008; Lehner, 2011; Mackay, 2014). Though we did not simulate such scenarios in this study, because two interacting loci would better maintain polymorphisms at the selected loci than would two non-interacting ones (Barton and Navarro, 2002; Navarro and Barton, 2002; Tennessen, 2018), it would not be surprising that they would produce even stronger footprints than what we observe here.

Furthermore, genomic sequences with multiple nearby balanced loci will have more extended footprints of balancing selection. With the capability to optimize over window sizes, *B* statistics should be more sensitive to such regions than other approaches applied with small fixed windows. Indeed, *B*_2_ substantially outperforms *T*_2_ (applied with 12 informative sites on either side of a test site) when the two neighboring loci under selection are weakly advantageous themselves (Figures S45E and F). The margins between their powers still persist even when *T* statistics adopt windows with 122 informative sites on either side of the test site (Figures S46E and F), despite the marginal increases in their powers for two-locus balancing selection.

Our exploratory results not only imply that extant approaches for detecting balancing selection have high power when applied to genomic regions carrying multiple balanced loci, but that such power are also likely much higher than they would have for single-locus regions. For *B* statistics in particular, because they optimize over window sizes, the gap between their sensitivity for multi-locus balancing selection and that for single-locus settings may be more profound than other methods when applied with small windows. Our results also support the speculation that top candidates identified in previous scans for balancing selection may be more likely to carry more than one functional polymorphic site, as is the case for the MHC locus, considering all methods we evaluated show higher powers for multi-locus balancing selection than for the single-locus process.

### Application of *B*_2_ to empirical data

In this study, we applied the *B*_2_ statistic on both human and bonobo genomic data, and identified sensible candidate targets in each species. We first re-examined the CEU and YRI human populations in the 1000 Genomes Project dataset (The 1000 Genomes Project Consortium, 2015) with *B*_2_, which have been previously probed for long-term balancing selection in multiple studies (DeGiorgio et al., 2014; Siewert and Voight, 2017; Bitarello et al., 2018). We found that top candidates reported by *B*_2_ overlap largely with previous scans, lending confidence in the power of *B* statistics to make replicable discoveries. Next, we performed the first model-based scan for footprints of balancing selection on bonobo polymorphism data. In addition to the genomic regions previously reported to be under ancient balancing selection in humans and chimpanzees (*e.g*., the *HLA-DQ* genes at the MHC locus; Leffler et al., 2013; Teixeira et al., 2015; Cheng and DeGiorgio, 2019), we have also uncovered novel candidates such as *KLRD1* and *SCN9A*,which play roles in pathogen defense and pain perception, respectively. Our results may correspond to the unique features and evolutionary history of bonobos, as suggested by accumulating evidence (de Waal, 1990; Hare et al., 2012; de Groot et al., 2017; Wroblewski et al., 2017; Maibach and Vigilant, 2019) on bonobo behavior and physiology.

### Potential balancing selection on gamete-associated genes in humans

In the scans of human populations, we recovered previously reported candidates *STPG2* (formerly *C4orf13*) and *CCDC169* (formerly *C13orf38*), in addition to the HLA-D locus and *ERAP2*. Neither of the two former genes was discussed in previous studies after reporting them as top candidates, probably due to their late characterization. Intriguingly, both genes are related to gametogenesis, with recent association and clinical studies underscoring their functional importance. In particular, the expression of *STPG2* has been detected in male tissues, endocrine tissues, as well as the brain (Uhlén et al., 2015). Structural mutations deleting this gene have been linked to azoospermia (Yakut et al., 2013) and velocardiofacial syndrome (Wu et al., 2019), and association studies of SNPs in this have correlated it with autism (Connolly et al., 2017) and preclampsia (Johnson et al., 2012). A recent study even reported footprints of ongoing positive selection on a segregating preclampsia-associated SNP in this gene (Arthur, 2018). Note that these authors only analyzed the disease-associating variants and applied haplotype-based selection tests, which tend to reveal regions with at least one dominant haplotype. The footprints reported by Arthur (2018) can result from either recent partial sweeps or balancing selection, with only the latter matching the kilobase-scale size of the increased diversity surrounding the region (Figures S23 and S24).

The conjoined gene *CCDC169-SOHLH2* encodes a read-through transcript of the gene *CCDC169* and its immediate downstream *SOHLH2*, a crucial gene for gametogenesis. In addition to its potential to initiate the transcription of *SOHLH2* on occasions of read-through, *CCDC169* has also been found to have specific expression in pre-natal brain tissues (Pletikos et al., 2014). More interestingly, the *B*_2_ scores across this gene do not form a typical peak as seen in many other candidate regions (Figures S25 and S26). Instead, we observed a plateau of elevated *B*_2_ scores above the region joining the two genes. Furthermore, both the mean pairwise sequence difference (*π*) and *T*_2_ with a 22-informative-site-radius window show two minor peaks across this region. Considering our results for multi-locus balancing selection (Figure S45), such footprints may be reflective of multiple loci undergoing balancing selection, probably interactively via epistasis, which can create footprints of extended tracks of elevated genetic diversity (Barton and Navarro, 2002; Navarro and Barton, 2002).

Lastly, despite the intriguing functional implications behind our candidates, we are aware that some of our candidate regions show worrying signs for artifacts. For example, *STPG2* (also a top candidate in the scan by DeGiorgio et al., 2014) has low 35-mer sequence uniqueness scores across the whole 40 kb region examined, despite surviving the 50-mer mappability filter. The peak linking *CCDC169* and *SOHLH2* shows overall higher sequence uniqueness than *STPG2*, but the few regions with relatively lower uniqueness co-localize with the peaks reported by *π* and *T*_2_. This co-localization is also observed in the gene *CPE*, where peak regions with a drop in sequence uniqueness also display lower sequencing depths than other regions. Though not all regions with low mappability necessarily yield outstanding scores for balancing selection, these signs could still be indicative of erroneous mapping and warrant further investigation and caution in interpretation.

### Footprints of balancing selection in bonobos and their implications

As one of the two sister species to humans, bonobos (initially known as the pygmy chimpanzees; Prüfer et al., 2012) have been drawing increasing attention from the genomics community (*e.g*., Prüfer et al., 2012; Prado-Martinez et al., 2013; de Manuel et al., 2016). However, compared with chimpanzees (the other sister species), bonobos are relatively understudied, despite their close relationship to humans and unique social behaviors. For bonobos, one of their most idiosyncratic traits is their high prevalence of sociosexual activities (de Waal, 1990; Kano, 1992; Wrangham, 1993), which serve important non-reproductive functions and include frequent same-sex encounters. As a close relative to humans, their female-dominance, low-aggression, and hypersexual social behaviors contrast fiercely with those of humans and chimpanzees (Kano, 1992; Wrangham, 1993), and has challenged the traditional understanding of intra- and inter-sexual relationships in hominid societies (Parish et al., 2000). In fact, a standing hypothesis postulates self-domestication as a main player in the evolution of bonobos, in that their unique social behavior has driven the selection against aggression and shaped other related traits (Hare et al., 2012). A growing number of recent studies have also characterized the differences in physiological responses between bonobos and chimpanzees behind their social behaviors (Heilbronner et al., 2008; Hohmann et al., 2009; Wobber et al., 2010; Deschner et al., 2012; Surbeck et al., 2012), yet the genetic component underlying their unique behaviors, however, remains largely elusive.

From the *B*_2_ scan of bonobo genomes, we identified a number of interesting top candidates involved in pathogen defense. Despite that most of the MHC region was removed by a mappability filter (see *Methods*), we still observed extraordinary signals from the remainder of this region. More specifically, the *HLA-DQA1* and *HLA-DQB1* genes harbor the highest peak across the genome (Figures 5 and S31). These two genes encode the α and *β* chains of HLA-DQ molecule, which is a surface receptor on antigen-presenting cells (Ball and Stastny, 1984), and has long been known to be highly polymorphic in great apes (Takahata et al., 1992; Prüfer et al., 2012; Teixeira et al., 2015).

Another immune-related gene, *KLRD1*, which encodes the cell surfacr antigen CD94, also exhibited outstanding *B*_2_ scores. The interaction between KLRD1 (CD94) and NKG2 family proteins can either inhibit or activate the cytotoxic activity of NK cells (Pende et al., 1997; Cantoni et al., 1998; Masilamani et al., 2006), as well as pivot the generation of cell memory in NK cells (Cerwenka and Lanier, 2016). Furthermore, KLRD1 (CD94) has been shown to play an important role in combating viral infections such as cytomegalovirus (CMV; Cerwenka and Lanier, 2016) and influenza (Bongen et al., 2018) in humans, as well as the mousepox virus in mice (Fang et al., 2011). In humans and chimpanzees, *KLRD1* is highly conserved (Khakoo et al., 2000; Shum et al., 2002). Here, the involvement in viral defense of *KLRD1* presents an especially intriguing case for bonobos. Bonobos have been recently shown to harbor reduced levels of polymorphism in MHC class I genes (Maibach et al., 2017; Wroblewski et al., 2017), which were further predicted to have lower ability to bind with viral peptides when compared with chimpanzees (Maibach and Vigilant, 2019). The genes encoding another regulator of MHC class I molecules, the Killer cell Immunoglobin-like Receptors (KIR), were also found to have contracted haplotypes in bonobos (Rajalingam et al., 2001; Walter, 2014; Wroblewski et al., 2019), with the lineage III *KIR* genes serving reduced functions (Wroblewski et al., 2019). In fact, many studies have pointed out that these reduced features are unlikely the natural consequences of demographic factors—even after considering the harsher bottlenecks bonobos have undergone compared with chimpanzees—and speculate that selective sweeps in bonobos on these regions (Prüfer et al., 2012; Walter, 2014; Maibach et al., 2017; Wroblewski et al., 2017, 2019) may have eliminated the diversity in these critical immunity genes. In this light, the polymorphisms on *KLRD1* may be compensating the reduced diversity in their binding partners in bonobos.

Several other genes in high-scoring regions are also found to be involved in immunity. For one, the highest peak on chromosome 7 encompasses the entire gene *GPNMB* (Figure S36), with elevated *B*_2_ scores particularly on exons. This gene encodes osteoactivin, a transmembrane glycoprotein found on osteoclast cells, macrophages, and melanoblast (Loftus et al., 2009; Yu et al., 2016), and is shown to regulate proinflammatory responses (Ripoll et al., 2007). Aside from its heavy involvement in cancer (Zhou et al., 2012), the protein GPNMB has also been shown to facilitate tissue repair (Li et al., 2010; Rose et al., 2010; Hu et al., 2013) as well as influence iris pigmentation (Bächner et al., 2002; Maric et al., 2013). Other potential evidence for balancing selection operating on innate immunity-related genes includes the high *B*_2_ scores observed around the intergenic region between *BPIFB4* and *BPIFA2* (Figure S37), which encode two Bacterialcidal Permeability-Increasing Fold-containing (BPIF) family proteins. The *BPIFA2*genic region is recently shown to harbor many SNPs significantly associated with enteropathy (Fujimori et al., 2019), whereas the *BPIFB4* gene is better-known by its association with longevity (Villa et al., 2015b; Spinetti et al., 2017; Villa et al., 2018), speculated to partly result from its protection of vascular functions (Villa et al., 2015a; Puca et al., 2016; Spinelli et al., 2017)

In addition to pathogen defense, we also found other interesting candidates relating to neurosensory and neurodevelopment. One such gene is *SCN9A* (Figure S32), which encodes Na_V_ 1.7, a voltage-gated sodium channel, with mutations on the gene associated with various pain disorders (Yang et al., 2004; Cox et al., 2006; Reimann et al., 2010). The peak we observe covers the overlapping RNA gene encoding its anti-sense transcript, *SCN1A-AS1*, which regulates the expression of *SCN9A* (Koenig et al., 2015), suggestive of diversified regulation of pain perception in bonobos. Also relating to neural functions, the other two major signals on chromosome 8 sit in *CSMD1* and *CSMD3* (Figures S33 and S34, respectively), both of which have been associated with neurological conditions such as bipolar disorder (Xu et al., 2014), autism (Floris et al., 2008), schizophrenia (Håvik et al., 2011; Kwon et al., 2013), and epilepsy (Shimizu et al., 2003). Despite that both genes are large in size (approximately two and 1.5 Mb, respectively) and may be more likely to harbor increased densities of polymorphisms by chance, the high peaks we observe mostly locate on exons, lending increased support for selection to diversify their functions.

Lastly, we noticed that some candidate genes carry multiple distinct functions, and may have been undergoing balancing selection due to potential evolutionary conflicts between some of their functions. For example, the gene *CSMD1* is not only associated with antibody response to smallpox vaccines (Ovsyannikova et al., 2012), it also plays a role in neuro-development (Kwon et al., 2013; Xu et al., 2014; Athanasiu et al., 2017). Likewise, the gene *GPNMB* plays roles not only in tissue repair (Li et al., 2010), but also in iris pigmentation (Bächner et al., 2002). Another candidate, *PDE1A* gene (Figure S35), encodes a phosphodiesterase that is pivotal to Ca^2+^- and cyclic nucleotide-signaling (Lefièvre et al., 2002). It is expressed in brain, endocrine tissues, kidneys, and gonads (Uhlén et al., 2015), and has multiple splicing variants. In fact, the high-scoring peak we observed on this gene happens to locate around the exons that are spliced out in some variants (Figure S35). Studies have demonstrated the relation of this gene to brain development (Yan et al., 1994), mood and cognitive disorders (Xu et al., 2011; Martinez and Gil, 2013; Pekcec et al., 2018; Betolngar et al., 2019), and hypertension (Kimura et al., 2017). Meanwhile, the PDE1A protein is also a conserved component of mammalian spermatozoa (Lefièvre et al., 2002; Vasta et al., 2005), and is involved in the movement of flagella—organelles featured in both animal sperms and some protozoa such as the *Trypanosoma* parasites (Oberholzer et al., 2007), with which the endemic geographical regions are avoided by bonobos (Inogwabini and Leader-Williams, 2012). Though it is difficult to judge for these genes which functions may be subject to selective pressures, they nonetheless indicate that pleiotropy can be an important driver of balancing selection.

### Concluding remarks

Extant methods for detecting long-term balancing selections are constrained by the pliability of their inferences as a function of genomic window size. In this study, we presented *B* statistics, a set of composite likelihood ratio statistics based on nested mixture models. We have comprehensively evaluated their performances through simulations and demonstrated their robust high performances over varying window sizes in uncovering genomic loci undergoing balancing selection. Moreover, we showed that even when applied with the least optimal window sizes, the *B* statistics still exhibit high power comparable to current methods, which operated under optimal window sizes, in uncovering balancing selection of varying age and selection parameters, as well as robust performance under confounding scenarios such as elevated mutation rates, variable recombination rates, and population size changes. We re-examined the 1000 Genomes Project YRI and CEU populations with *B*_2_ statistics, and have recovered well-characterized genes previously-hypothesized to be undergoing long-term balancing selection in humans, such as the HLA-D genes, *ERAP2*, and *CSMD2*. We also characterized previously-reported top candidates *STPG2*and *CCDC169-SOHLH2*, both of which are related to gametogenesis. We further applied the *B*_2_ statistic on the whole-genome polymorphism data of bonobos, and discovered not only the well-established *MHC-DQ* genes, but also novel candidates such as *KLRD1, PDE1A, SCN9A*, and *CSMD1*, with functional implications ranging from pathogen defense to neuro-behavior. Moreover, we have extended the *B* statistics to consider multi-allelic balancing selection, with these extensions demonstrating superior properties to all previous methods for detecting selected loci with more than two balanced alleles. We also show that methods tend to have higher powers for two-locus balancing selection than for single-locus processes. Lastly, we have implemented these statistics in the open source software BalLeRMix, which, along with other key scripts used in this study, can be accessed at https://github.com/bioXiaoheng/BalLeRMix/. We have also released the empirical scan results for balancing selection in both humans and bonobos, which can be downloaded at http://degiorgiogroup.fau.edu/ballermix.html.

## Methods

In this section, we discuss sets of simulations used to evaluate the performances of the *B* statistics relative to previously-published state-of-the-art approaches (Hudson et al., 1987; DeGiorgio et al., 2014; Siewert and Voight, 2017, 2018; Bitarello et al., 2018). Finally, we describe the application of our *B* statistics to an empirical bonobo dataset (Prado-Martinez et al., 2013).

### Evaluating methods through simulations

We employed the forward-time genetic simulator SLiM (version 3.2; Haller and Messer, 2019) to generate sequences of 50 kb in length evolving with or without balancing selection. Based on the respective levels in humans and other great apes, we assumed a mutation rate of *μ* = 2.5 × 10^−8^ per base per generation (Nachman and Crowell, 2000), and a recombination rate of *r* = 10^−8^ per base per generation (Payseur and Nachman, 2000). In scenarios with constant population sizes, we set the diploid effective population size as *n* = 10^4^. To create baseline genetic variation, each replicate simulation was initiated with a burn-in period of 10*N* = 10^5^ generations. To speed up simulations, we applied the scaling parameter λ to the number of simulated generations, population size, mutation rate, recombination rate, and selection coefficient, which allows for the generation of the same levels of variation with a speed up in computational time by a factor λ^2^. For scenarios based on a model of constant population size, we used λ = 100. For the demographic models of European humans and bonobos, we used λ = 20. We simulated 500 replicates for each scenario considered, and sampled 50 haploid lineages from the target population and one lineage from the outgroup in each simulation for downstream analyses.

We simulated data from two other diverged species, under the demographic history inspired by that of humans, chimpanzees (Kumar et al., 2005), and gorillas (Scally et al., 2012). Specifically, the closer and farther outgroups diverged 2.5 × 10^5^ and 4 × 10^5^ generations ago, respectively, which correspond to five million and eight million years ago, assuming a generation time of 20 years.

To evaluate the power of each method to detect balancing selection with varying selective coefficient *s*, dominance coefficient *h*, and age, for each combination of *s* and *h*, we considered 15 time points at which the selected allele was introduced, ranging from 5 × 10^4^ to 6.5 × 10^5^ generations prior to sampling with time points separated by intervals of 5 × 10^4^ generations. Assuming a generation time of 20 years, these time points are equivalent to 1,2,3,…, 15 million years before sampling. In each scenario, a single selected mutation was introduced at the center of each sequence at the assigned time point, and we only considered simulations where the introduced allele was not lost.

### Accelerated mutation rate

To evaluate whether the *B* statistics are robust to high mutation rates, we applied the methods on simulated sequences evolving neutrally along the same demographic history (Figure S1), but instead with a fivefold higher mutation rate of 5*μ* = 1.25 × 10^−7^ per site per generation. To generate sequences with regional increases in mutation rate, we simulated 50 kb sequences with a five-fold higher mutation rate of 5*μ* = 1.25 × 10^−7^ per site per generation at the central 10 kb of the sequence, and the surrounding region with the original rate *μ*.

### Recombination rate estimation error

For evaluating the robustness to erroneous estimation of recombination rates, we simulated sequences with uneven recombination maps, and applied the model-based methods with the assumption that the recombination rate is uniform. In particular, we divided the 50 kb sequence into 50 regions of one kb each, and in turns inflate or deflate the recombination rate of each region by *m* fold, such that the recombination rates of every pair of neighboring regions have a *m*^2^-fold difference. We tested *m* = 10 and *m* = 100 in this study.

### Demographic history

To examine the performance of methods under realistic demographic parameters, we considered the demographic histories of a European human population (CEU; Terhorst et al., 2017) and of bonobos (Prado-Martinez et al., 2013). For the human population, we adopted the history of population size changes inferred by SMC++ (Terhorst et al., 2017) that spans 10^5^ generations, assuming a mutation rate of *μ*= 1.25 × 10^−8^ per site per generation (assumed when estimating the CEU demographic history in Terhorst et al., 2017), a generation time of 20 years, and a scaling effective size of 10^4^ diploids. To account for recombination rate variation, we allowed each simulated replicate to have a uniform recombination rate drawn uniformly at random between *r* = 5 × 10^−9^ and *r* = 1.5 × 10^−8^ per site per generation. We also simulated an additional population that split from the human population 2.5 × 10^5^ generations ago, which is identical to the outgroup (named O1) in the demographic model depicted in Figure 3A, with an effective size of *N* = 10^4^ diploid individuals.

For the bonobo population history, we scaled the PSMC history inferred from the genome of individual A917 (Dzeeta; sample SRS396202) by Prado-Martinez et al. (2013) with a mutation rate of *μ* = 2.5 × 10^−8^ per site per generation, identical to the simulations on the three-population demographic history (Figure 3A). Because the inferred PSMC model provides a specific ratio of the mutation and recombination rates, we set the recombination rate to *r* = 2.84 × 10^−9^ per site per generation. To be consistent with the three-population demographic history, we set the population size prior to 71,640 generations ago, which is the maximum time covered by the PSMC inference, to *N* = 10^4^ diploid individuals, and had the outgroup split 2.5 × 10^5^ generations ago with the same diploid population size, identical to the outgroup O1 in the three-population demographic history (Figure 2A).

To simulate species with distinct mutation rates, we split the simulation into two stages, with the first stage concerning the sequences in the ancestral species up until the two populations diverge five million years ago. Upon divergence, two separate SLiM simulations are used to distinguish the mutation rates in the target and outgroup populations, and samples are output separately before being integrated in subsequent analyses. We set the target species to mutate at a rate of *μ* = 1.2 × 10^−8^ per site per generation (Scally and Durbin, 2012) after divergence, and the other species (including the ancestral species) evolving with the mutation rate of *μ* = 2.5 × 10^−8^ per site per generation (Nachman and Crowell, 2000). The recombination rate across all simulations is *r* = 10^−8^ per site per generation (Payseur and Nachman, 2000). For the simulations with constant population sizes, we set the effective size of all populations as *N* = 10^4^ diploid individuals, and adopted the scaling parameter λ = 100. For simulations employing realistic demographic histories, we used λ = 20, set the effective population size of the ancestral and the outgroup species as *N* = 10^4^ diploids (Takahata et al., 1995), and the target species following the demographic history inferred from the CEU human population (Terhorst et al., 2017) for 10^5^ generations prior to sampling. Additionally, we set the generation time of the target species to be 25 years (akin to humans; Scally and Durbin, 2012), while for the outgroup and ancestral species we used 20 years (akin to non-human great apes; Prado-Martinez et al., 2013). Consequently, the species divergence occurred 200,000 generations ago for the target species, and 250,000 generations ago for the outgroup.

### Three- and four-allelic balancing selection at a single site

To simulate balancing selection on a single site with more than two balanced alleles, we used SLiM3.3 (Haller and Messer, 2019) so that all four nucleotides, instead of binary representations of 0s and 1s, can be incorporated into the simulations. We adopted the same three-species demographic history as illustrated in Figure S1, and simulated sequences of length 50 kb consisting of randomly-generated strings of four nucleotides at the beginning of each replicate, with equal chance of occurrence for each nucleotide. We considered the Jukes-Cantor substitution model and set the between-nucleotide mutation rate as *μ* = 8.3 ×10^−9^ per site per generation, such that the total mutation rate (three times the between-nucleotide mutation rate) is *μ*= 2.49 × 10^−8^ per site per generation—roughly the same as adopted in the bi-allelic balancing selection simulations. We also assumed a uniform recombination rate of *r* = 10^−8^ per site per generation (Payseur and Nachman, 2000). At 500,000 generations before sampling, we introduced two, three, or four mutations of distinct nucleotides that have selective coefficient *s* = 0.001 and dominance coefficient *h* = 20. Note that SLiM considers co-localized mutations of distinct types as if they were at different positions, and computes fitness for the individual by multiplying fitness values of each mutation. That is, a diploid individual who is heterozygous at a site harboring two distinct selectively advantageous mutant alleles with parameters *s* = 0.001 and *h* = 20 would have fitness (1 + *hs*)(1 + *hs*) = 1.44, whereas a homozygote for either selectively advantageous mutation would have fitness 1 + *s* = 1.001. At the completion of the simulation, we sampled 25 diploid individuals uniformly at random from each of the sister species (P and O1), and one diploid individual was sampled uniformly at random from species O2, with only one haplotype of this individual being considered as the reference sequence. Only bi-allelic sites were considered in the downstream analysis.

### Application to empirical data

#### Human genomic data from the 1000 Genomes Project

We obtained variant calls from the 1000 Genomes Project dataset (The 1000 Genomes Project Consortium, 2015), which were mapped to human reference genome hg19, and extracted the haplotypes for the CEU and YRI populations. We used the chimpanzee reference genome panTro5 downloaded from the UCSC Genome Browser (Kent et al., 2002; Haeussler et al., 2018) to call ancestral alleles, and only retained mappable monomorphic or bi-allelic polymorphic sites based on the variation in the CEU (or YRI) population together with the chimpanzee reference genome. For mappable sites not included in the variant call dataset, we assumed the site is monomorphic for the hg19 reference genome, and called substitutions accordingly.

To avoid making inference on potentially problematic regions, we applied the RepeatMasker filter and removed segmental duplications, both of which were downloaded from the UCSC Genome Browser (Kent et al., 2002; Haeussler et al., 2018). Genomic regions with mappability 50-mer score (Derrien et al., 2012) lower than 0.9 were discarded as well. Moreover, we performed one-tailed Fisher’s exact tests for Hardy-Weinberg equilibrium (Wigginton et al., 2005) on each polymorphic site and removed those with a significant (*p* < 10^−4^) excess of heterozygous genotypes.

We applied *B*_2_ to each CEU and YRI dataset separately, assuming the human recombination map of the hg19 reference genome (International HapMap Consortium, 2007). We did not fix the window size of these scans, and instead permitted *B*_2_ to optimize over both free parameters *A* and *x*. To better compare our results with previous studies, we also applied the *T*_2_ statistic (DeGiorgio et al., 2014) to the same input datasets, adopting window sizes of 22 or 100 informative sites on either side of a test informative site. We also computed sequence diversity *π* averaged across each five kb window as a reference.

For downstream examination of the mappability of candidate regions, we consulted the 35-mer uniqueness score (UCSC hg19 database; Kent et al., 2002; Haeussler et al., 2018) averaged across each one kb region. Furthermore, we also downloaded the BAM files for each individual in the CEU or YRI population and generated per-base read depths with BEDTools 2.26 (Quinlan, 2014). We then computed sample-wide mean read depths, their standard deviations, and the number of individuals without coverage for each population after merging read depths of all samples with BEDTools. These references further aided in flagging potentially problematic regions that survived initial filters, as they typically feature lower mappability (mean 35-mer uniqueness) or abnormally low or high read depths.

#### Bonobo genomic data from the Great Ape Project

We obtained the genotype calls of 13 bonobos from the Great Ape Project (Prado-Martinez et al., 2013), which were mapped to human genome assembly NCBI36/hg18. We lifted over the variant calls to human genome assembly GRCh38/hg38, and polarized the allele frequencies with the bonobo genome assembly panPan2, with the sequence in hg38 considered as the ancestral allele. Only genomic regions mappable across hg38 and panPan2 were considered for further analyses. For mappable polymorphic sites, we only considered bi-allelic SNPs. For mappable sites without variant calls in bonobo, we assumed these sites were monomorphic for the panPan2 reference genome sequence, and called substitutions based on whether the panPan2 reference allele was different from the hg38 reference allele.

To circumvent potential artifacts, we performed one-tailed Hardy-Weinberg equilibrium tests on each site and removed sites with an excess of heterozygotes (*p* < 0.01). This *p*-value was determined by the distribution of the *p*-values of all polymorphic sites across the genome, such that 0.035% of such sites are outliers. We also applied the RepeatMasker, segmental duplication, simple repeat, and interrupted repeat filters (all downloaded from UCSC Genome Browser) to remove repetitive regions. To assess the mappability of each genomic region, we employed the mappability module GEMtools (Derrien et al., 2012), and computed the mappability tracks of 50-mers. Regions with 50-mer mappability scores lower than 0.9 were removed. Because `BalLeRMix` employs a pre-specified grid of *A* values to accompany the distances *d* in centi-Morgans (cM), when applying the method, we assumed a uniform recombination rate of 10^−6^ cM per site, which is the approximate recombination rate in humans (Payseur and Nachman, 2000).

## Acknowledgments

We thank J. Terhorst and J. Prado-Martinez for sharing the inferred parameters for the demographic history of CEU and great apes, repectively. We also appreciate the assistance from B. Haller on the fitness scheme of multi-allelic balancing selection with SLiM. We are also grateful to three anonymous reviewers for their comments that helped improve this manuscript. This work was funded by National Institutes of Health grant R35GM128590, by National Science Foundation grant DEB-1753489, and by the Alfred P. Sloan Foundation. Portions of this research were conducted with Advanced CyberInfrastructure computational resources provided by the Institute for CyberScience at Pennsylvania State University.

