## Supplementary Note for "Flexible mixture model approaches that accommodate footprint size variability for robust detection of balancing selection"

#### 1 Mixture models for multi-allelic balancing selection

##### 1.1 Balancing selection with $m$ alleles

For balancing selection with $m$ alleles, denote each allele as $A_1, A_2, \dots, A_m$, and their corresponding equilibrium frequencies as $x_1, x_2, \dots, x_m$, with $\sum_{i=1}^m x_i = 1$. To avoid redundancy, we index the alleles by decreasing frequency, such that $x_1 \geq x_2 \geq \cdots \geq x_m$.

###### 1.1.1 Probability distributions for allele frequencies of bi-allelic neutral sites linked to an $m$-allelic balanced locus

Consider $n$ sampled lineages, and denote the counts and frequencies of each of the $m$ balanced alleles, $A_1, A_2, \dots, A_m$, at the selected locus as $\mathbf{n} = [n_1, n_2, \dots, n_m]$ and $\mathbf{x} = [x_1, x_2, \dots, x_m]$, respectively. Assuming that the numbers of each of the $m$ allelic categories at the selected locus follows a multinomial distribution, we have

$$\mathbb{P}[\mathbf{n}; n, \mathbf{x}] = \frac{n!}{\prod_{i=1}^m n_i!} \prod_{i=1}^m x_i^{n_i}.$$

For any neutral bi-allelic site linked to the selected locus, suppose $k$ derived alleles are observed out of the $n$ sampled lineages, and denote the number of derived alleles linked to each balanced allele on the selected locus by the vector $\mathbf{k} = [k_1, k_2, \dots, k_m]$. We suppose that the $k_i$ derived alleles are completely linked with balanced allele $A_i$. That is, a non-zero $k_i$ value would be considered equal to $n_i$, the number of copies of balanced allele $A_i$ at the selected locus that this neutral site is linked to. Correspondingly, denote the vector of numbers of ancestral alleles at this neutral site linked to each balanced allelic class as $\mathbf{j} = \mathbf{n} - \mathbf{k} = [j_1, j_2, \dots, j_m]$. It follows that any non-zero $j_i$ reflects the number of $A_i$. In other words, $k_i$ is either 0 or $n_i$, and likewise for $j_i$. We can denote $M \subseteq \{1, 2, \dots, m\}$ as the subset of allele class indices at the selected locus linked to a derived allele at the bi-allelic neutral site, and hence

$$k_i = n_i \mathbf{1}_{\{i \in M\}},$$
$$j_i = n_i \mathbf{1}_{\{i \notin M\}},$$

such that for derived neutral alleles,

$$\sum_{i=1}^m k_i = \sum_{i \in M} n_i = k,$$

and for ancestral neutral alleles,

$$\sum_{i=1}^m j_i = \sum_{i \notin M} n_i = n - k.$$

Note that, for each possible vector $\mathbf{k}$, there could be more than one vector $\mathbf{j} = \mathbf{n} - \mathbf{k}$ partition that satisfies the relationships described above, as there can exist more than one way to partition the $j_i$ ancestral alleles into being linked to the remaining $\overline{M} = \{1, 2, \dots, m\} \setminus M$ balanced allele classes at the selected locus (see section 1.2 of the *Supplementary Note* for finding all possible partitions). For example, when $n = 10$, $m = 4$, and $k = 5$, for one possible partition $\mathbf{k} = [0, 3, 0, 2]$, $\mathbf{n}$ can be $[5, 3, 0, 2]$, $[0, 3, 5, 2]$, $[4, 3, 1, 2]$, $[1, 3, 4, 2]$, $[3, 3, 2, 2]$, or $[2, 3, 3, 2]$. It follows then that possible partition vectors $\mathbf{j} = \mathbf{n} - \mathbf{k}$ of the ancestral alleles is $[5, 0, 0, 0]$, $[0, 0, 5, 0]$, $[4, 0, 1, 0]$, $[1, 0, 4, 0]$, $[3, 0, 2, 0]$, or $[2, 0, 3, 0]$. Likewise, there will be multiple sets of partition vectors $\mathbf{j}$ of ancestral alleles that allow the partition vector $\mathbf{k}$ of derived alleles to be, say, $[3, 2, 0, 0]$, $[2, 0, 3, 0]$, or $[0, 0, 4, 1]$, such that $\sum_{i=1}^m k_i = k = 5$ and $\sum_{i=1}^m n_i = n = 10$.

Let $S^*_m(k,n)$ denote the total number of such ways to separate $n$ lineages into $m$ allelic classes and distribute $k$ derived and $j = n-k$ ancestral alleles across the $m$ allelic classes, with the restriction that in each allelic class $A_i$, $n_i$ equates to either $k_i$ or $j_i$. We consider each of these ways to be equally likely to happen. Therefore, by summing the probability for all possible $\mathbf{k}$ and $\mathbf{j}$ linked neutral derived and ancestral allele partitions respectively, the general form for the probability to observe $k$ derived alleles out of $n$ sampled lineages on a linked neutral site is

$$h_{n,m,\mathbf{x}}(k) = \frac{1}{S^*_m(k,n)} \sum_{\substack{\mathbf{k}=[k_1,k_2,\ldots,k_m] \\ k_i\in\{0,1,\ldots,k\} \\ \sum_{i=1}^m k_i=k}} \sum_{\substack{\mathbf{j}=[j_1,j_2,\ldots,j_m] \\ j_i\in\{0,1,\ldots,n-k\} \text{ if } k_i=0 \\ j_i=0 \text{ if } k_i>0 \\ \sum_{i=1}^m (k_i+j_i)=n}} \mathbb{P}[\mathbf{k}+\mathbf{j};n,\mathbf{x}]. \tag{1}$$

Note that because the set of feasible $\mathbf{k}$ with $\mathbf{n}$ combinations is only a subset of all possible $\mathbf{n}$ combinations, it is sensible that $\sum_{k=0}^n h_{n,m,\mathbf{x}}(k) < 1$, and thus it is necessary to normalize all $h_{n,m,\mathbf{x}}(k)$ across the range values considered for $k$. Furthermore, because we expect the ancestral alleles to be linked to at least one allelic class at the selected locus, for neutral sites with $k=n$, this one class of balanced alleles is hence considered lost or not observed, and the total number of allelic class that the $k=n$ derived alleles are linked to would therefore be $m-1$ instead of $m$.

To tailor the distribution to different input types, normalize $h_{n,m,\mathbf{x}}(k)$ based on the possible values of $k$. Following the nomenclature adopted for the two-allele balancing selection (see *Theory*), we use "0", "0,MAF", "1", "2", and "2,MAF" to annotate the distributions normalized based on derived allele frequencies only (*i.e.*, $k = 1,2,\ldots,n-1$), minor allele frequencies only (*i.e.*, $k = 1,2,\ldots,\lfloor \frac{n}{2} \rfloor$), polymorphic or substitution calls only (*i.e.*, $k \neq n$ or $k = n$), substitution calls with derived allele frequencies (*i.e.*, $k = 1,2,\ldots,n$), and substitution calls with minor allele frequencies (*i.e.*, $k = 1,2,\ldots,\lfloor \frac{n}{2} \rfloor$ for polymorphisms and $k=0$ for substitutions), respectively. The probability mass functions for these normalized distributions can be computed as

$$\begin{aligned}
h^{(0)}_{n,m,\mathbf{x}}(k) &= \frac{h_{n,m,\mathbf{x}}(k)}{\sum_{k=1}^{n-1} h_{n,m,\mathbf{x}}(k)} && k \in \{1,2,\ldots,n-1\} \\
h^{(0,\text{MAF})}_{n,m,\mathbf{x}}(k) &= \frac{h_{n,m,\mathbf{x}}(k) + \mathbf{1}_{\{k\neq n/2\}} h_{n,m,\mathbf{x}}(n-k)}{\sum_{k=1}^{n-1} h_{n,m,\mathbf{x}}(k)} && k \in \{1,2,\ldots,\lfloor n/2 \rfloor\} \\
h^{(1)}_{n,m,\mathbf{x}}(k) &= \frac{\mathbf{1}_{\{k=n\}} h_{n,m,\mathbf{x}}(n) + \mathbf{1}_{\{k\neq n\}} \sum_{i=1}^{n-1} h_{n,m,\mathbf{x}}(i)}{\sum_{k=1}^{n} h_{n,m,\mathbf{x}}(k)} && k \in \{1,2,\ldots,n\} \\
h^{(2)}_{n,m,\mathbf{x}}(k) &= \frac{h_{n,m,\mathbf{x}}(k)}{\sum_{k=1}^{n} h_{n,m,\mathbf{x}}(k)} && k \in \{1,2,\ldots,n\} \\
h^{(2,\text{MAF})}_{n,m,\mathbf{x}}(k) &= \frac{\mathbf{1}_{k\neq 0} h_{n,m,\mathbf{x}}(k) + \mathbf{1}_{\{k\neq n/2\}} h_{n,m,\mathbf{x}}(n-k)}{\sum_{k=1}^{n} h_{n,m,\mathbf{x}}(k)} && k \in \{0,2,\ldots,\lfloor n/2 \rfloor\}.
\end{aligned}$$

On the other hand, let $\xi_n(k)$ denote the number of sites with $k$ derived alleles observed out of $n$ sampled lineages in the whole genome (or chromosome), and let $\eta_n(k) = \xi_n(k)\mathbf{1}_{\{k\neq 0\}} + \xi_n(n-k)\mathbf{1}_{\{k\neq n/2\}}$. Under neutrality, the probability to observe $k$, $k = 1,2,\ldots,n$ derived alleles on sites unlinked with the balanced locus takes the general form

$$g_n(k) = \frac{\xi_n(k)}{\sum_j \xi_n(j)}. \tag{2}$$

Likewise, with respect to the type of input data, the normalized probability distributions for neutrality can be obtained as

$$g_n^{(0)}(k) = \frac{\xi_n(k)}{\sum_{j=1}^{n-1}\xi_n(j)} \qquad k \in \{1, 2, \ldots, n-1\}$$

$$g_n^{(0,\mathrm{MAF})}(k) = \frac{\eta_n(k)}{\sum_{j=1}^{n-1}\eta_n(j)} \qquad k \in \{1, 2, \ldots, \lfloor n/2 \rfloor\}$$

$$g_n^{(1)}(k) = \frac{\mathbf{1}_{\{k=n\}}\xi_n(k) + \mathbf{1}_{\{k\neq n\}}\sum_{i=1}^{n-1}\xi_n(i)}{\sum_{j=1}^{n}\xi_n(j)} \qquad k \in \{1, 2, \ldots, n\}$$

$$g_n^{(2)}(k) = \frac{\xi_n(k)}{\sum_{j=1}^{n-1}\xi_n(j)} \qquad k \in \{1, 2, \ldots, n\}$$

$$g_n^{(2,\mathrm{MAF})}(k) = \frac{\eta_n(k)}{\sum_{j=0}^{n-1}\eta_n(j)} \qquad k \in \{0, 2, \ldots, \lfloor n/2 \rfloor\}.$$

##### 1.1.2 Composite likelihood ratios based on the mixture models

For neutral site $d$ recombination units away from the selected locus, use $\alpha_{d,A} = e^{-Ad}$ to describe the chance of being linked to the balanced locus, the full model for observing $k$ derived alleles on this site can be considered as the mixture of the influence of balancing selection (the $h_{n,m,\mathbf{x}}(k)$ in Equation 1 multiplied by $\alpha_A(d) = e^{-Ad}$) and neutrality (the $g_n(k)$ in Equation 2 multiplied by $1 - \alpha_A(d)$).

$$f_{n,m,\mathbf{x},A}(k, d) = \alpha_A(d) \cdot h_{n,m,\mathbf{x}}(k) + [1 - \alpha_A(d)]g_n(k). \tag{3}$$

Let $\mathbf{d} = [d_1, d_2, ..., d_L]$ denote the vector of distances to the test site for $L$ informative sites across the genomic region of interest. The composite likelihood for the null hypothesis of neutrality would be $\mathcal{L}_0(\mathbf{n}, \mathbf{k}) = \prod_{i=1}^{L} g_{n_i}(k_i)$, and the composite likelihood for the alternative hypothesis, $\mathcal{L}_a(\mathbf{x}, A; \mathbf{n}, \mathbf{k}, \mathbf{d})$, of being linked to balancing selection can be maximized at

$$(\widehat{\mathbf{x}}, \widehat{A}) = \underset{\mathbf{x},\, A}{\arg\max} \prod_{i=1}^{L} f_{n_i,m,\mathbf{x},A}(k_i, d_i).$$

The general $B$ statistic can thus be formulated as

$$B = 2\Big[\ln \mathcal{L}_a(\widehat{\mathbf{x}}, \widehat{A}; \mathbf{n}, \mathbf{k}, \mathbf{d}) - \ln \mathcal{L}_0(\mathbf{n}, \mathbf{k})\Big]. \tag{4}$$

#### 1.2 Possible partitions of $k$ alleles into $m$ allelic classes

For $k$ identical elements and $m$ distinct non-empty classes, where $1 \le m \le k$, let $S_m(k)$ denote the number of ways to distribute these $k$ elements across the $m$ classes. Because the $m$ classes are distinct, they can therefore be ordered, and this allocation process becomes equivalent to dividing $k$ identical elements into $m$ ordered compartments, *i.e.*, inserting $m-1$ dividers into $k-1$ slots with each slot hosting no more than one divider. Therefore, the number of such ways to distribute $k$ elements across $m$ classes is

$$S_m(k) = \binom{k-1}{m-1}. \tag{5}$$

In the context of a bi-allelic neutral site with $k$ derived alleles in a sample of $n$ alleles that is linked to a locus balancing $m$ allelic classes, let $m_d$ and $m_a$ denote the numbers of balanced allelic classes that the $k$ derived alleles and $n-k$ ancestral alleles are completely linked with, respectively. Let $m_0 \in \{0, 1, \ldots, m\}$ denote the number of balanced allelic classes that are linked to at least one derived or ancestral allele at the neutral site. It follows that $m_0 = m_d + m_a$, with $1 \le m_d \le k$ and $1 \le m_a \le n-k$, and therefore $m_0 \ge 2$. Moreover, let $S_m^\star(k, n)$ denote the number of ways in which the $k$ derived and $n-k$ ancestral alleles at the linked neutral locus can be distributed across the $m$ balanced allelic classes, such that derived and ancestral alleles are not linked to the same allelic class. This total number of partitions can be counted by summing the product of $S_{m_d}(k)$ and $S_{m_a}(n-k)$ for each pair of $m_d$ and $m_a$, and for each possible $m_0$ value, and its value is given by

$$\begin{aligned} S^{\star}_m(k,n) &= \sum_{m_0=2}^{m} \binom{m}{m_0} \left[ \sum_{m_d=1}^{\min\{m_0-1,k\}} \binom{m_0}{m_d} S_{m_d}(k) S_{m_0-m_d}(n-k) \cdot \mathbf{1}_{\{n-k\geq m_0-m_d\}} \right], \\ &= S^{\star}_m(n-k,n); \quad k=1,2,\ldots,n-1. \end{aligned}$$

Note that in this formula, $k$ and $n-k$ are interchangeable. Because we assume the derived neutral allele be linked to at least one balanced allele, when $k=0$, at least one balanced allele would have zero representation in the data, and so $m_0$ cannot be greater than $m-1$ leading to

$$S^{\star}_m(n,n) = S^{*}_m(0,n) = \sum_{m_0=1}^{m-1} \binom{m}{m_0} S_{m_0}(n).$$

##### 1.3 Probability distributions for balancing selection with $m = 2$ alleles

When $m=2$, the number of ways to partition $k$ ($k=1,2,\ldots,n-1$) derived alleles is

$$\begin{aligned} S^{\star}_2(k,n) &= \sum_{m_0=2}^{2} \left[ \sum_{m_d=1}^{\min\{m_0-1,k\}} \binom{m_0}{m_d} S_{m_d}(k) S_{m_0-m_d}(n-k) \right], \\ &= \binom{2}{1} S_1(k) S_1(n-k) \ = 2. \end{aligned}$$

For sites with $k=n$ or $k=0$, the number of ways is $\binom{2}{1}S_1(k)=2$.

Intuitively, for all possible $k$ values, the two ways are when the $k$ alleles linked to either allele $A_1$ or allele $A_2$; *i.e.*, $\mathbf{k}$ is either $[0,k]$ or $[k,0]$. The corresponding $\mathbf{j}$ therefore takes the respective value of $[n-k,0]$ or $[0,n-k]$. Let $\mathbf{x}=[x,1-x]$. With these values plugged into Equation 1, the probability of observing $k$ derived alleles on a neutral site in complete linkage with a two-allele balanced locus would hence take the general form

$$\begin{aligned} h_{n,x}(k) &= \frac{1}{2}\Big[\frac{n!}{k!(n-k)!}x^k(1-x)^{n-k} + \frac{n!}{(n-k)!k!}(1-x)^k x^{n-k}\Big], \\ &= \frac{1}{2}\binom{n}{k}\Big[x^k(1-x)^{n-k} + (1-x)^k x^{n-k}\Big]; \quad k=0,1,2,\ldots,n \end{aligned}$$

This formulation is consistent with the ones described in *Theory*. Because the probability mass function for neutrality stays the same regardless of selection, the formulation of $g_n(k)$ would not change. Therefore, the resulting mixture models are consistent with those adopted in the main manuscript.

### 2 Matching the number of sites in a genomic region with fixed length

In general, we considered two types of sliding windows in the simulation study when evaluating each statistic's robustness to large window sizes. The first type of sliding window was a region with a fixed number of informative sites, whereas the second was a region with a fixed number of nucleotides (nt). Because `BALLET` (*i.e.*, $T_1$ and $T_2$) and `BetaScan` (*i.e.*, $\beta$ and $\beta^{(2)}$) adopt different types of sliding windows, we applied $B$ statistics in both ways separately so as to better compare their performances with current methods. To approximately match the lengths of genomic regions analyzed by $T$ statistics and the summary statistics, we calculated the expected number of informative sites $I_n$ for a sample of $n$ lineages, based on the parameters of the simulated neutral demographic histories.

#### 2.1 Constant-size demographic history relating three species with a uniform mutation rate

For the majority of our simulations, we adopted a constant-size demographic history inspired by great apes that relates a population P to two outgroup populations O1 and O2 (depicted in Figure S1). Under this scenario, O1 and O2 diverged from P $t_1 = 2.5 \times 10^5$ and $t_2 = 4 \times 10^5$ generations ago, respectively. For these calculations, we assumed the population size is a constant $N = 10^4$ diploid individuals across the entire demographic model. In our analyses, we randomly sampled $n = 50$ lineages from population P and used one lineage in either population O1 or O2 to call substitutions and to polarize allele frequencies in the sample from P. Let $C_1$ and $C_2$ denote the inter-species coalescence time between P and O1 or O2, respectively, and let $H_n$ and $L_n$ be the expected tree height and tree length for a sample of $n$ lineages taken from P, respectively. Under neutrality, these quanities can be computed as (Ewens, 1974; Watterson, 1975)

$$
\begin{aligned}
C_1 &= \frac{t_1}{2N} + 1 \\
&= \frac{2.5 \times 10^5}{2 \times 10^4} + 1 = 13.5 \\
C_2 &= \frac{t_2}{2N} + 1 \\
&= \frac{4 \times 10^5}{2 \times 10^4} + 1 = 21
\end{aligned}
$$

and

$$
\begin{aligned}
H_{50} &= 2\left(1 - \frac{1}{n}\right) \\
&= 2\left(1 - \frac{1}{50}\right) = 1.96 \\
L_{50} &= 2\sum_{k=1}^{n-1} \frac{1}{k} \\
&= 2\sum_{k=1}^{50-1} \frac{1}{k} \approx 8.96.
\end{aligned}
$$

Therefore, the expected numbers of informative sites for using either O1 or O2 as the outgroup are respectively

$$
\begin{aligned}
I_{50}(\text{P-O1}) &= 2N\mu(2C_1 - H_n + L_n) \\
&= 2(10^4)(2.5 \times 10^{-8})(2C_1 - H_{50} + L_{50}) \approx 0.016999 \text{ per nucleotide} \\
I_{50}(\text{P-O2}) &= 2N\mu(2C_2 - H_n + L_n) \\
&= 2(10^4)(2.5 \times 10^{-8})(2C_2 - H_{50} + L_{50}) \approx 0.024499 \text{ per nucleotide,}
\end{aligned}
$$

assuming a per-site per-generation mutation rate of $\mu = 2.5 \times 10^{-8}$.

That is, when using O2 as the outgroup, a window with a radius of 10 informative sites (21 informative sites in total) on average approximately spans a genomic region of length 857 nt, and when using O1 as the outgroup this length is approximately 1235 nt. With O2 as the outgroup, window sizes of 1, 2, 3, 5, 10, 15, 20, or 25 kb should on average approximately correspond to windows of a fixed radius of 12, 24, 36, 60, 122, 183, 245, and 367 informative sites, respectively.

#### 2.2 Demographic history relating two species with realistic parameters

For simulations adopting the realistic demographic histories of the human CEU population or of bonobos, mutation rates and recombination rates are altered to correspond to the parameters used to infer the demographic histories. The theoretical footprint sizes as well as the density of informative sites consequently vary from sequences simulated under constant population size models.

Instead of $\mu = 2.5 \times 10^{-8}$ mutation and $r = 10^{-8}$ recombination events per site per generation, the CEU demographic history adopts a per site per generation mutation rate of $\mu = 1.25 \times 10^{-8}$ and a recombination rate drawn uniformly at random between $r = 5 \times 10^{-9}$ and $r = 1.5 \times 10^{-8}$ per site per generation. Given the range of recombination rates, the theoretical size of footprints would vary between five kb to approximately 1.67 kb. With the halved mutation rate, on the other hand, $I_{50}$(P-O1) would fall around 0.0085 per nucleotide, which translates to genomic regions covering on average 14.2 to 42.5 informative sites. To ensure that $T$ statistics consider enough data to reduce noise, we allow it to employ 10 informative sites at either side of the test informative site (21 informative sites in total covered by each window), and the corresponding mean physical length is approximately 2.5 kb.

For sequences simulated under the bonobo demographic history, the recombination rate was set to $r = 2.84 \times 10^{-9}$ per site per generation, under which the theoretical footprint size of long-term balancing selection is expected to be approximately 8.8 kb. Considering most tested statistics showed optimal power with one kb windows when the theoretical footprints was 2.5 kb, we adopted a window size of 3.6 kb here each summary statistic. With the $I_{50}$(P-O1) obtained under the mutation rate of $\mu = 2.5 \times 10^{-8}$, a 3.6 kb region should on average cover approximately 60 informative sites. We therefore set the window size for $T$ statistics as 29 sites on either side of the test informative site, such that each test considers a genomic region covering 59 informative sites.

#### 2.3 Demographic history relating two species with species-specific mutation rates

When simulating a demographic history with species-specific mutation rates, sequences of the target population mutates at a rate of $\mu_{\mathrm{H}} = 1.2 \times 10^{-8}$ per generation per site after splitting with the outgroup population, which has a constant per generation mutation rate of $\mu = 2.5 \times 10^{-8}$ per site, which is also the same mutation rate as the ancestral population. Moreover, assume that the target population and the group population diverge at the same number of generations as populations P and O1 (depicted in Figure S1).

Recall that we found the inter-species coalescence time between P and O1 to be $C_1 = 13.5$ coalescent units, and that we estimate the tree height and tree length for 50 sampled lineages in the target population to be $H_{50} = 1.96$ and $L_{50} \approx 8.96$, respectively.

It follows that the expected number of informative sites using O1 as the outgroup is

$$\begin{aligned}
I_{50}(\text{P-O1}) &= 2N\mu_{\mathrm{H}}(C_1 - 1 - H_{50} + L_{50}) + 2N\mu(C_1 + 1) \\
&= 2(10^4)(1.2 \times 10^{-8})(13.5 - 1 - 1.96 + 8.96) + 2(10^4)(2.5 \times 10^{-8})(13.5 + 1) \\
&= 0.01193 \text{ per nucleotide.}
\end{aligned}$$

Therefore, a one kb region would cover 11.93 informative sites, and a 2.5 kb region would cover approximately 29.825 informative sites. To again ensure $T$ statistics have enough data for each test, we adopted a window size of 15 informative sites on either side of the test informative site, and applied the summary statistics with windows of length 2.5 kb.

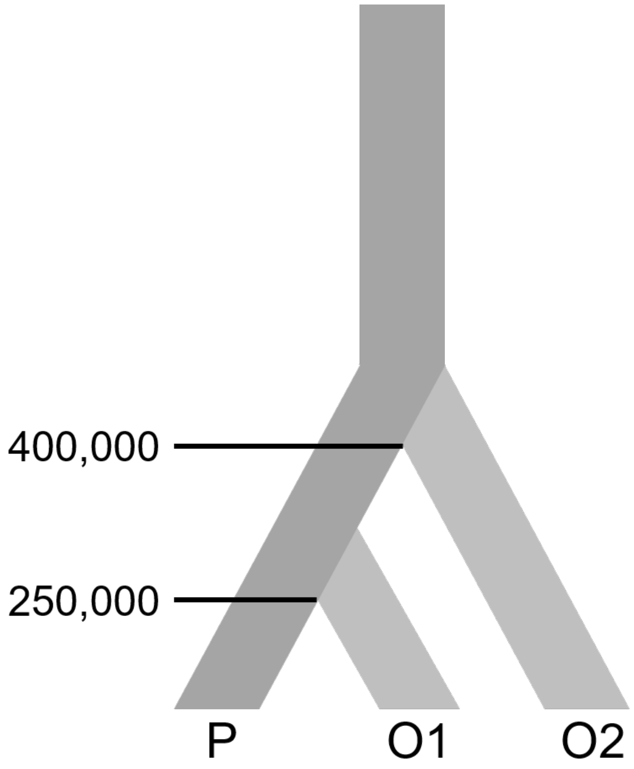


Figure S1: Schematic for the three-species demographic history considered in our simulation study. Divergence times of the closer (O1) and more distant (O2) outgroup are labeled in terms of the numbers of generations, and the population size across the entire phylogeny is $N = 10^4$ diploid individuals. In scenarios with balancing selection, the mutation under selection is only introduced in the lineage ancestral to population P, indicated in darker gray shading.

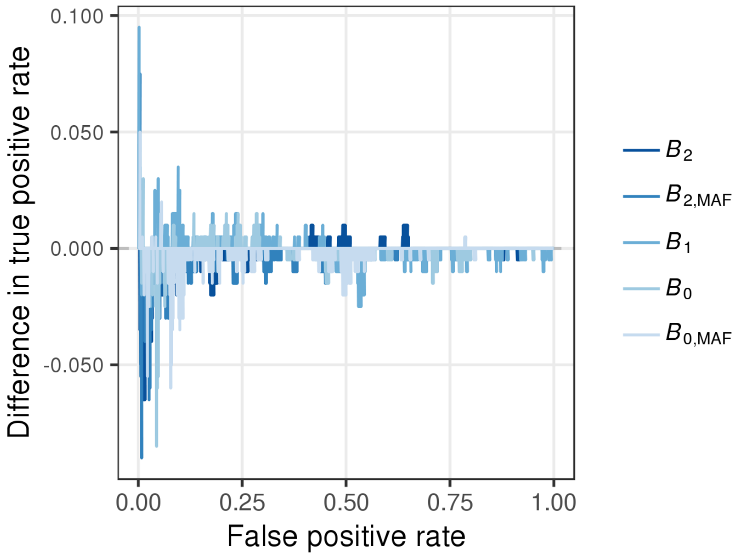


Figure S2: Effect of truncating the calculation of $B$ statistics, illustrated by the difference in true positive rate for each $B$ statistic as a function of false positive rate, where the difference in true positive rate for a $B$ statistic is computed as the true positive rate when using all available informative sites on a simulated sequence minus the true positive rate when using only sites in which $\alpha_A(d) \geq 10^{-8}$.

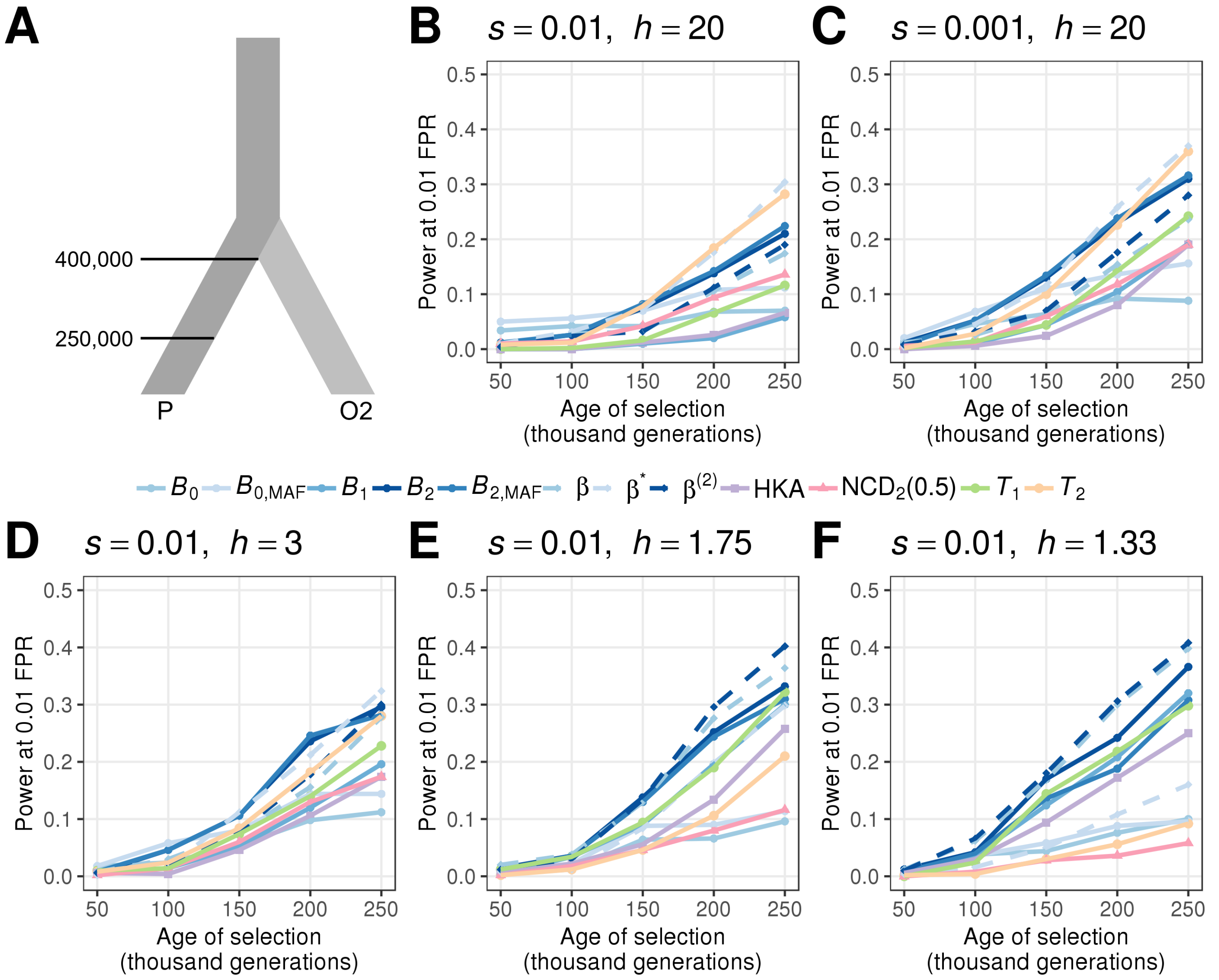


Figure S3: Ability to detect balancing selection for different heterozygote advantage scenarios occurring within 250,000 generations ago. (A) Demographic model relating the ingroup (P) and outgroup (O2) populations, with one sample from O2 used as the outgroup sequence. (B-F) Powers at a 1% false positive rate (FPR) for each statistic as a function of age of the allele undergoing balancing selection for different selection ($s$) and dominance ($h$) coefficients. The scenarios considered are (B) $s = 0.01$ with $h = 20$, (C) $s = 0.001$ with $h = 20$, (D) $s = 0.01$ with $h = 3$, (E) $s = 0.01$ with $h = 1.75$, and (F) $s = 0.01$ with $h = 1.33$. Note that the equilibrium frequencies for panels D, E, and F are 0.4, 0.3, and 0.2, respectively.

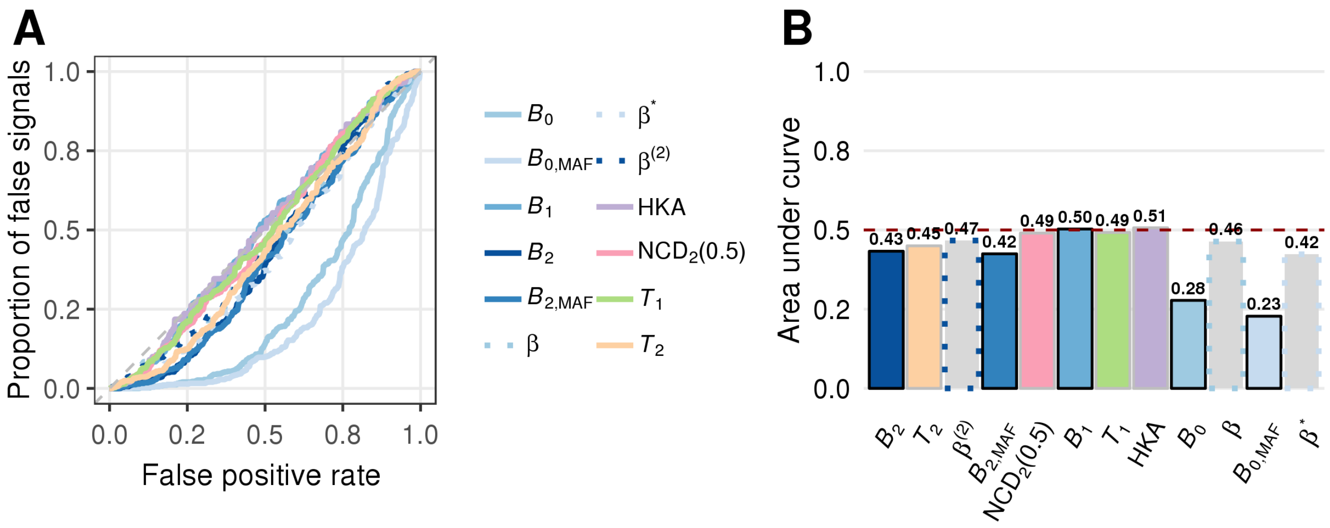


Figure S4: Performance of all statistics on neutral sequences with an uneven recombination map. (A) Proportions of false signal as a function of false positive rate. (B) Area under the curve for all statistics. Bars for all $B$ statistics are bordered by black lines, and all $\beta$ statistics are filled in gray.

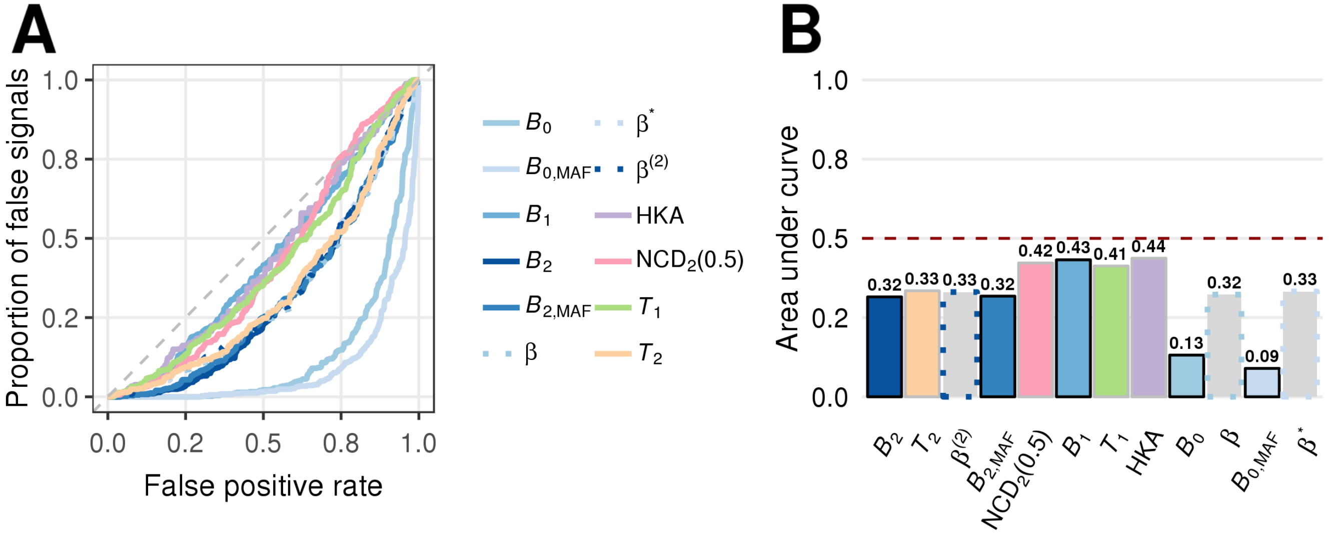


Figure S5: Robustness of each statistic to neutral sequences evolving under an uneven recombination map, with the recombination rate fluctuating from 100-fold higher to 100-fold lower than the uniform recombination rate $r$ between neighboring one kb windows. (A) Proportions of false signal as a function of false positive rate. (B) Area under the curve for all statistics. Bars for all $B$ statistics are bordered by black lines, and all $\beta$ statistics are filled in gray.

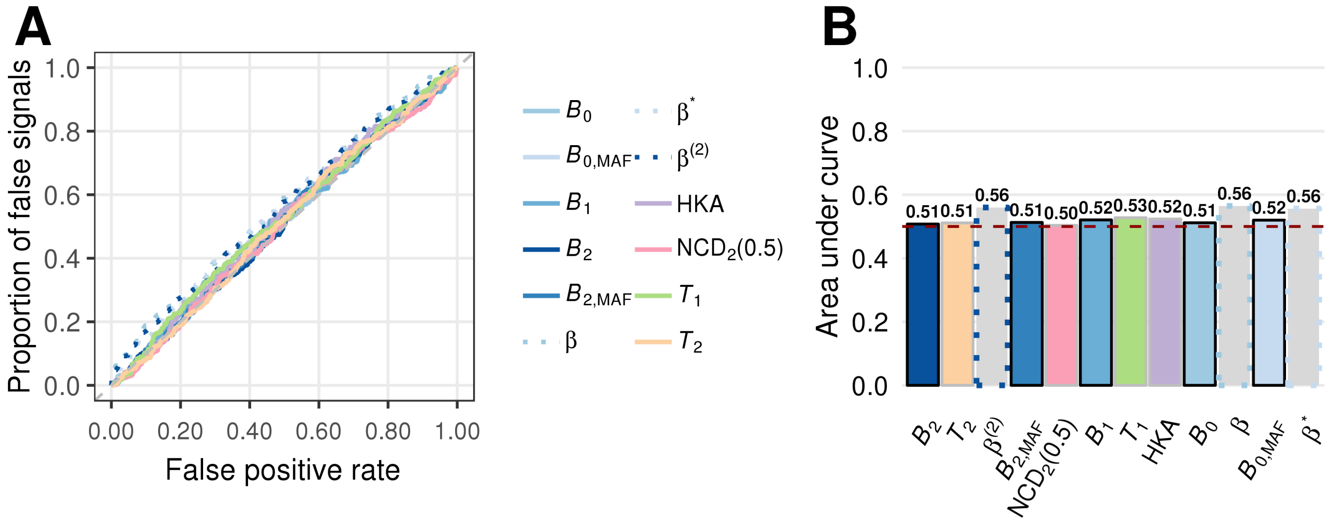


Figure S6: Robustness of each statistic evaluated when the mutation rate is elevated five-fold in a central 10 kb genomic region. (A) Proportions of false signals as a function of false positive rate (FPR). (B) Area under the curve for all statistics. Bars for all $B$ statistics are bordered by black lines, and all $\beta$ statistics are filled in gray.

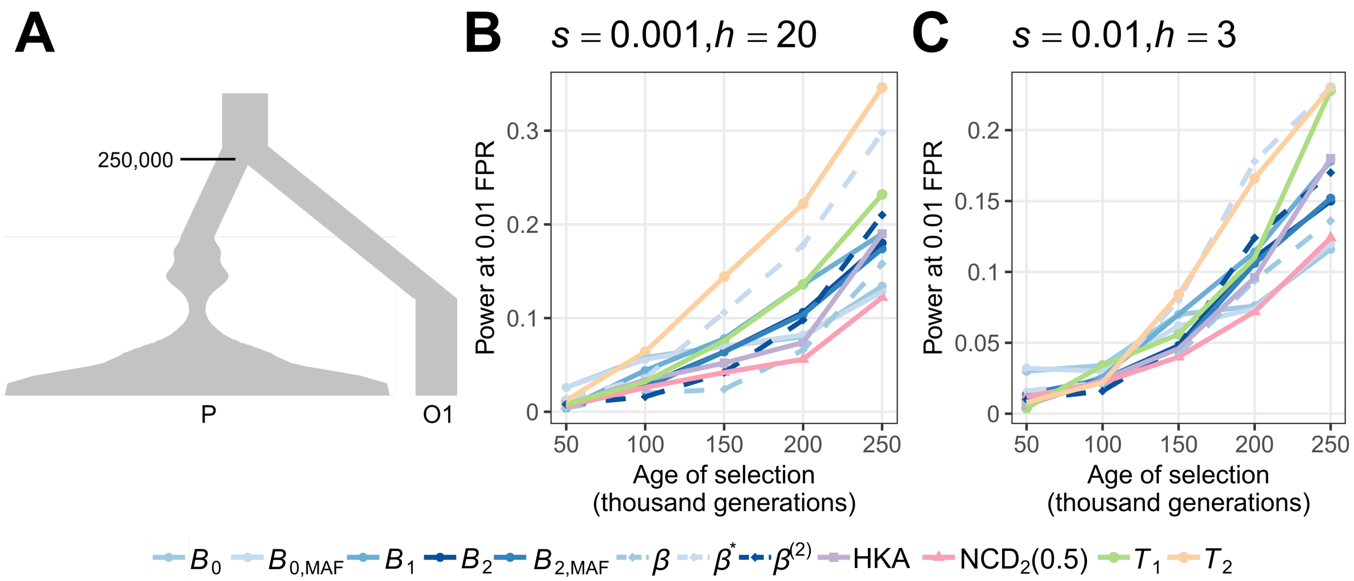


Figure S7: Power of all statistics as a function of the age of selection under the demographic model of CEU. (A) Schematic for the simulated demographic history. The outgroup O1 diverged from population P $2.5 \times 10^5$ generations ago. (B, C) Power at a 1% false positive rate (FPR) of each statistics for detecting balancing selection with selection parameters (B) $s = 0.001$ with $h = 20$ and (C) $s = 0.01$ with $h = 3$.

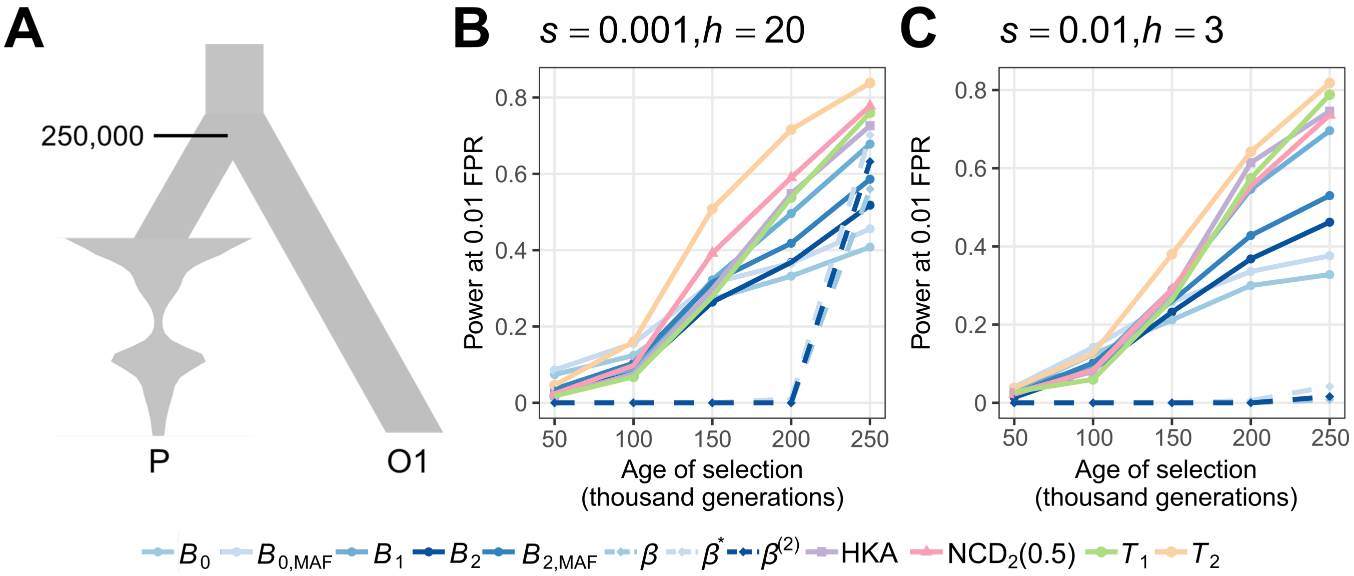


Figure S8: Power of all statistics as a function of the age of selection under the demographic model of bonobos. (A) Schematic for the simulated demographic history. The outgroup O1 diverged from population P $2.5 \times 10^5$ generations ago. (B, C) Power at a 1% false positive rate (FPR) of each statistics for detecting balancing selection with selection parameters (B) $s = 0.001$ with $h = 20$ and (C) $s = 0.01$ with $h = 3$.

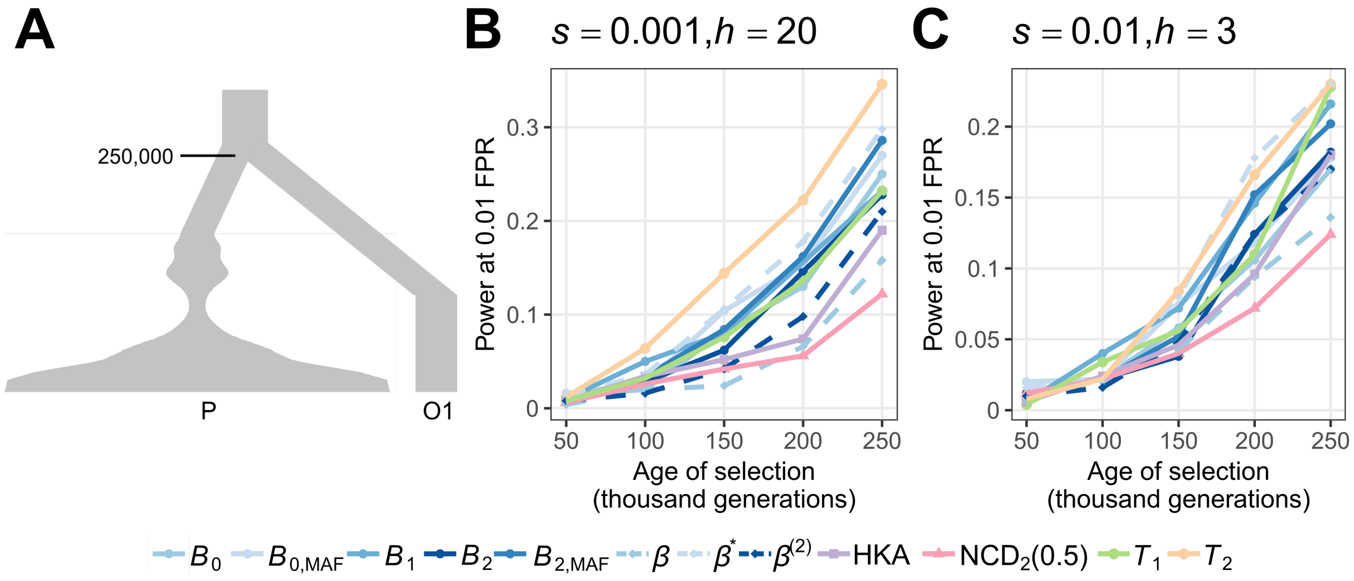


Figure S9: Power of $B$ statistics when provided with identical window sizes as $T$ statistics or $\beta$ statistics, in comparison with other statistics for detecting balancing selection with varying age and parameters under the demographic history of CEU. (A) Schematic of the demographic model of CEU. (B, C) Power at a 1% false positive rate (FPR) of each statistics for detecting balancing selection with selection parameters (B) $s = 0.001$ with $h = 20$ and (C) $s = 0.01$ with $h = 3$.

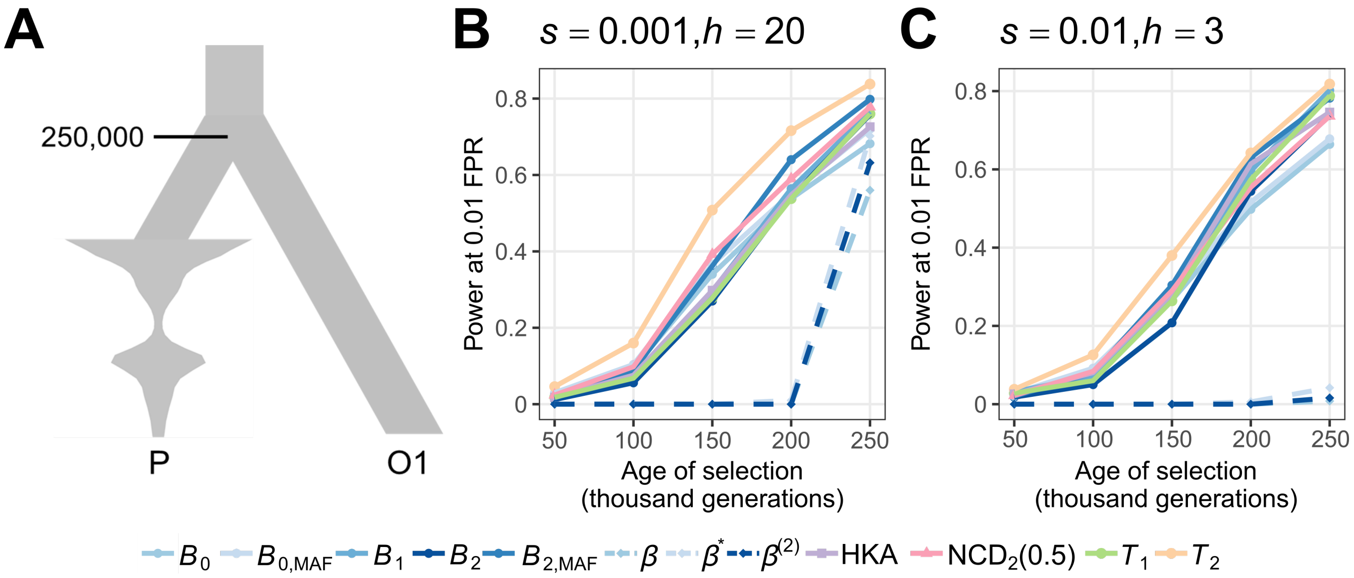


Figure S10: Power of $B$ statistics when provided with identical window sizes as $T$ statistics or $\beta$ statistics, in comparison with other statistics for detecting balancing selection with varying age and parameters under the demographic history of bonobos. (A) Schematic of the demographic model of bonobos. (B, C) Power at a 1% false positive rate (FPR) of each statistic for detecting balancing selection with selection parameters (B) $s = 0.001$ with $h = 20$ and (C) $s = 0.01$ with $h = 3$.

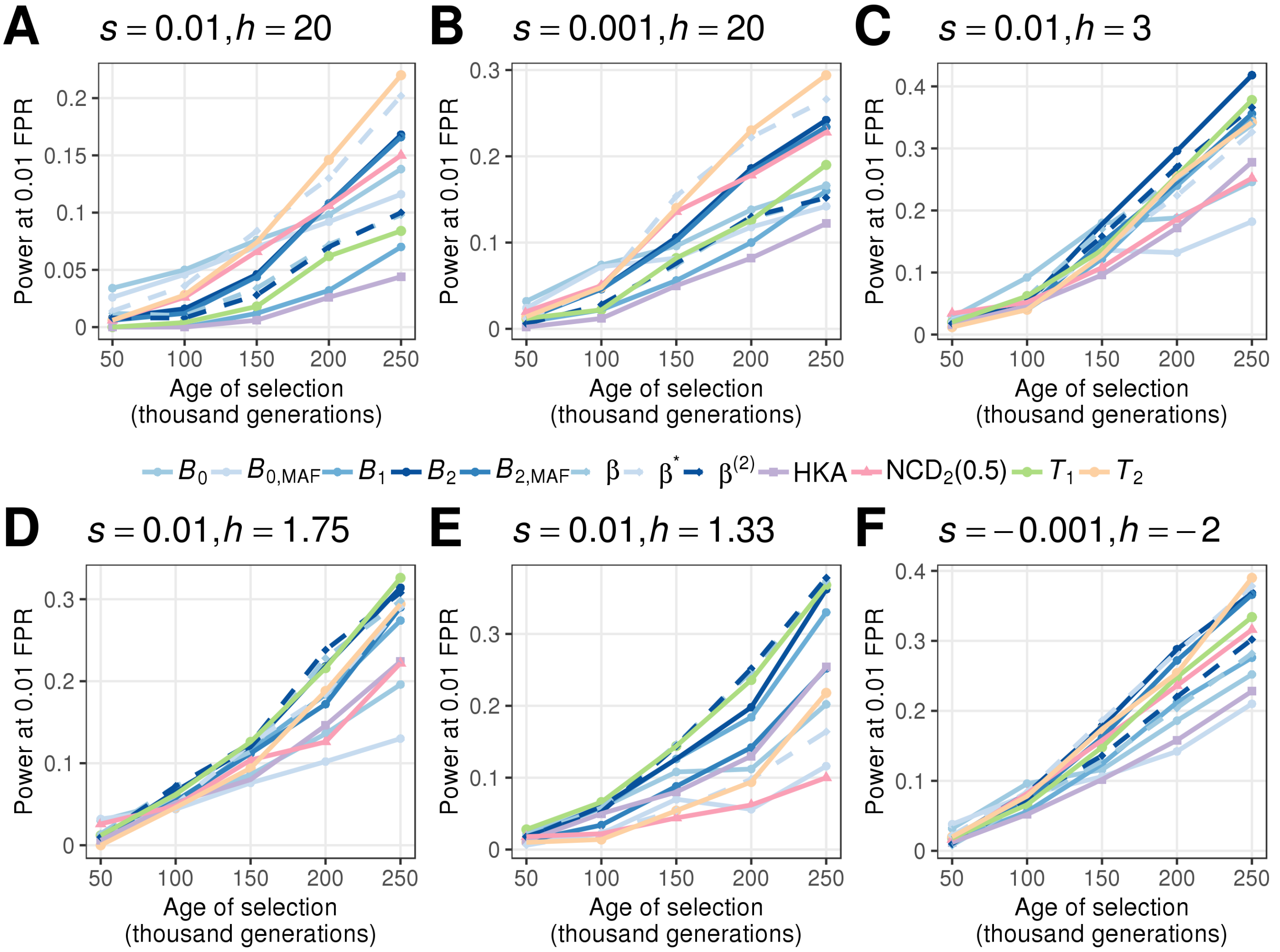


Figure S11: Powers at a 1% false positive rate (FPR) for each statistic as a function of age of the allele undergoing balancing selection for different selection ($s$) and dominance ($h$) coefficients when mutation rates vary between the target and outgroup species. The scenarios considered are (A) $s = 0.01$ with $h = 20$, (B) $s = 0.001$ with $h = 20$, (C) $s = 0.01$ with $h = 3$, (D) $s = 0.01$ with $h = 1.75$, (E) $s = 0.01$ with $h = 1.33$, and (F) $s = -0.001$ with $h = -2$.

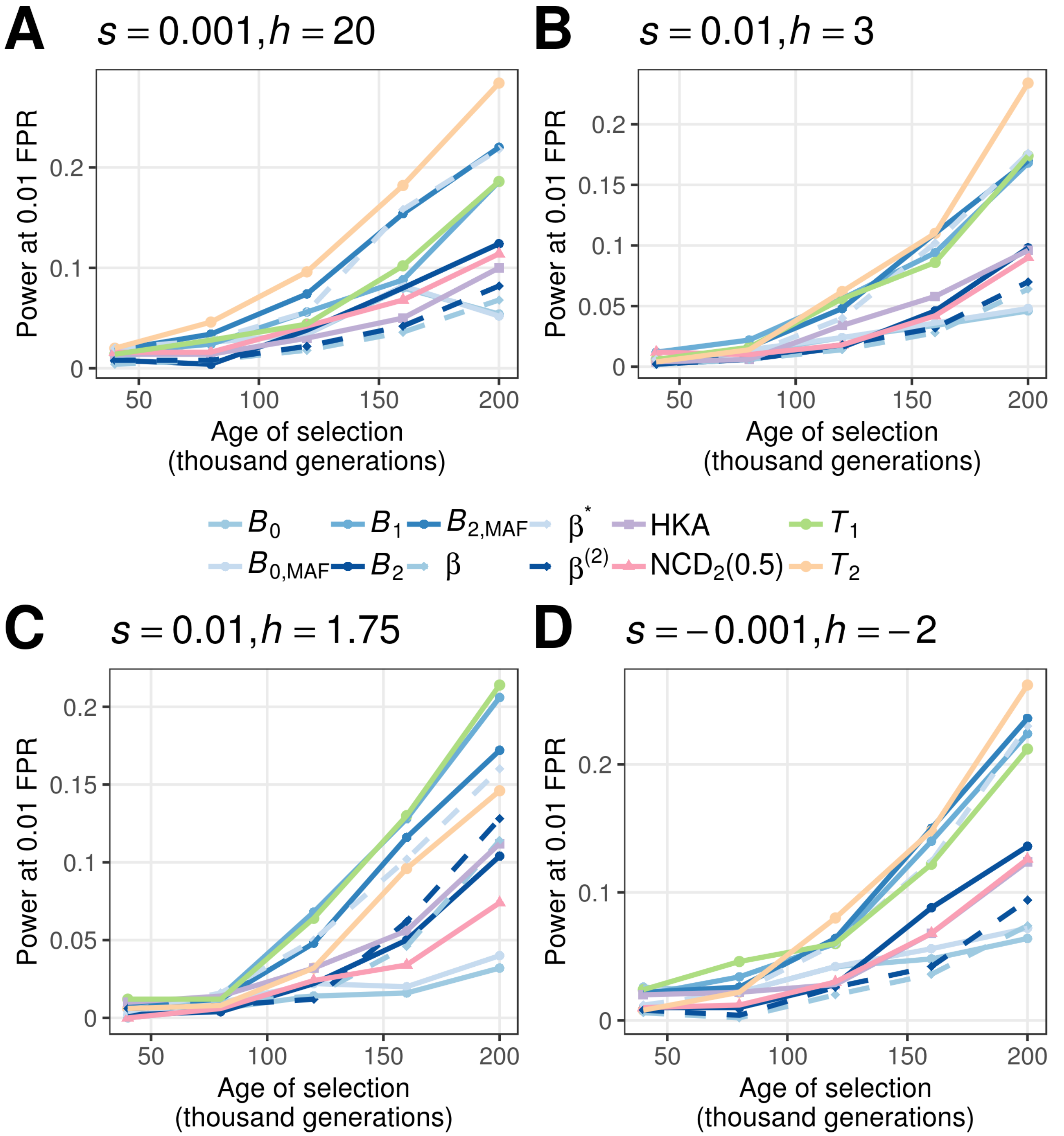


Figure S12: Powers at a 1% false positive rate (FPR) for each statistic as a function of age of the allele undergoing balancing selection for different selection ($s$) and dominance ($h$) coefficients when mutation rates vary between the target and outgroup species, and where the target species evolves under the demographic history of the CEU human population. The scenarios considered are (A) $s = 0.001$ with $h = 20$, (B) $s = 0.01$ with $h = 3$, (C) $s = 0.01$ with $h = 1.75$, and (D) $s = -0.001$ with $h = -2$.

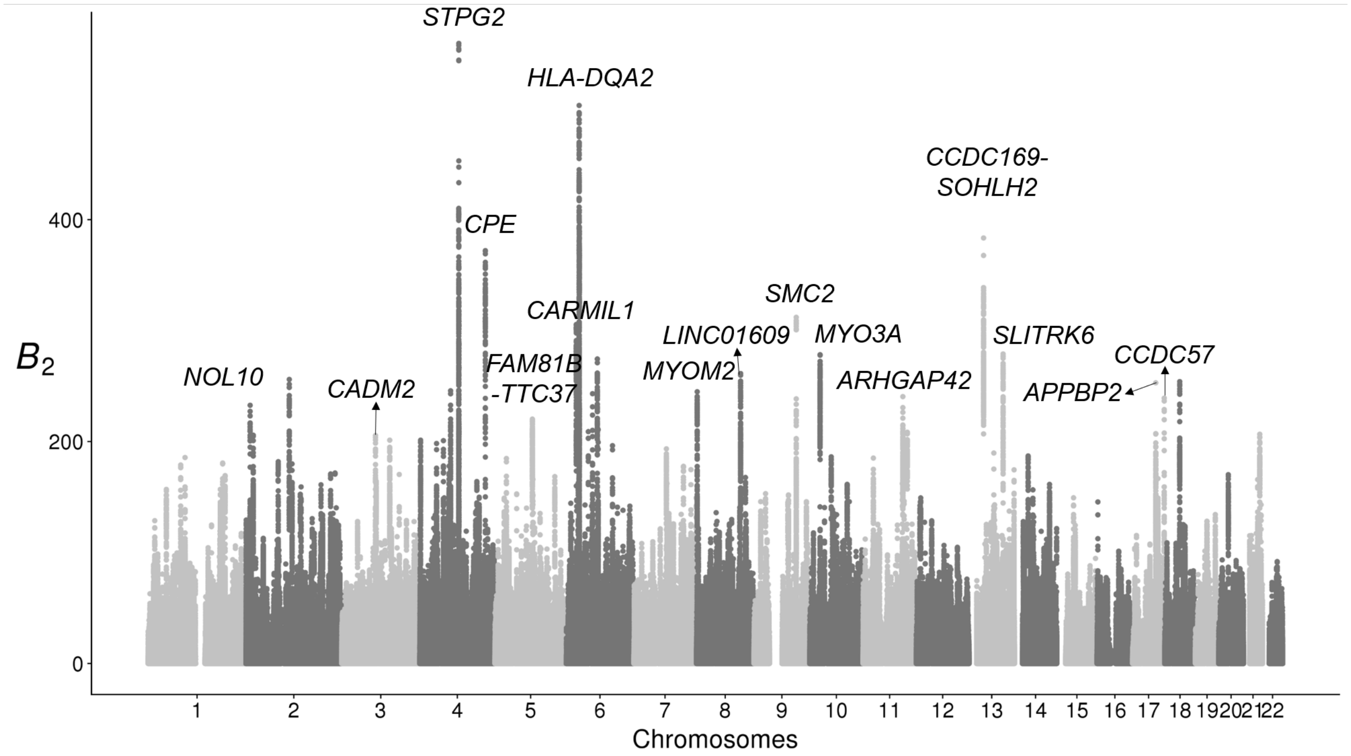


Figure S13: Manhattan plot displaying $B_2$ scores across the 22 human autosomes for the YRI population, with top candidates annotated. Outstanding peaks ($B_2 > 213$, top 0.01%) without annotations either fall in regions with abnormal sequencing depths or low sequence uniqueness, or do not neighbor any coding regions within a 100 kb radius.

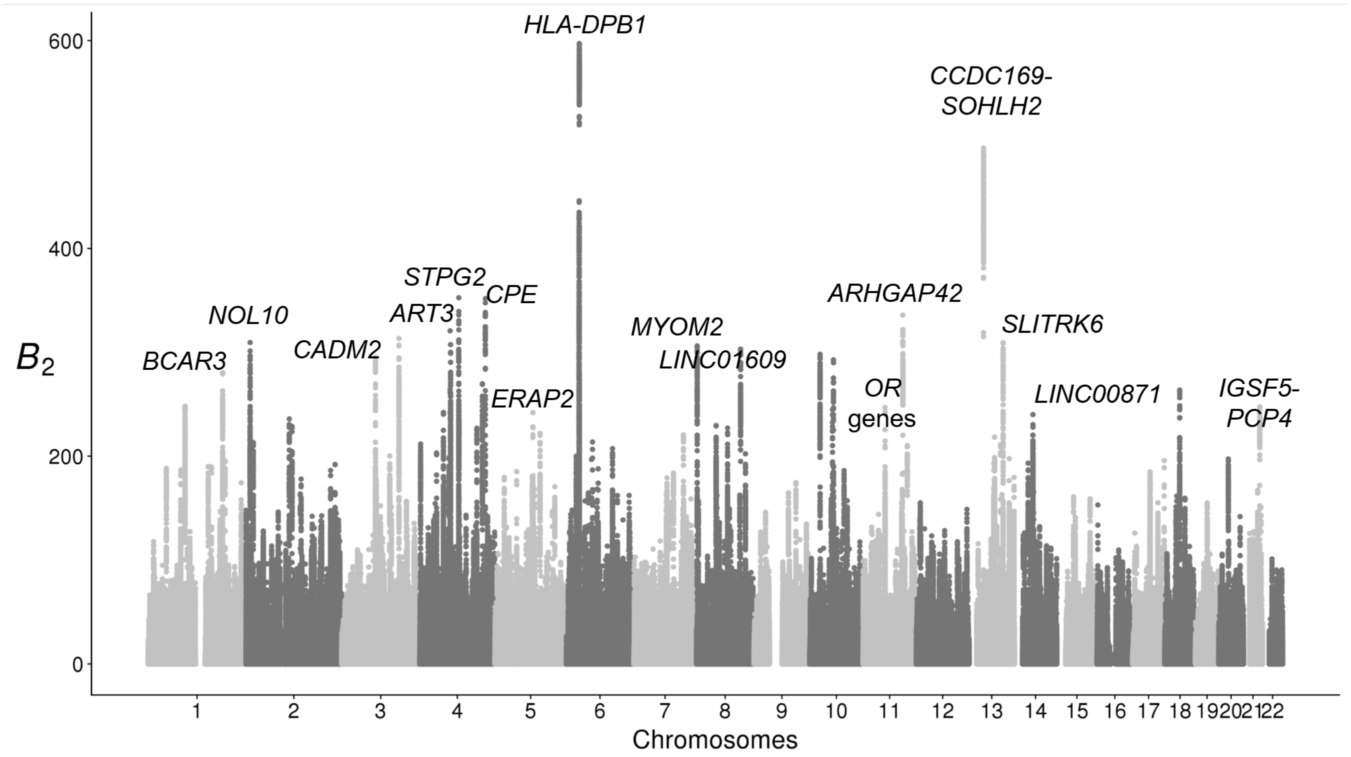


Figure S14: Manhattan plot displaying $B_2$ scores across the 22 human human autosomes for the CEU population, with top candidates annotated. Outstanding peaks ($B_2 > 241$, top 0.01%) without annotations either fall in regions with abnormal sequencing depths or low sequence uniqueness, or do not neighbor any coding regions within a 100 kb radius.

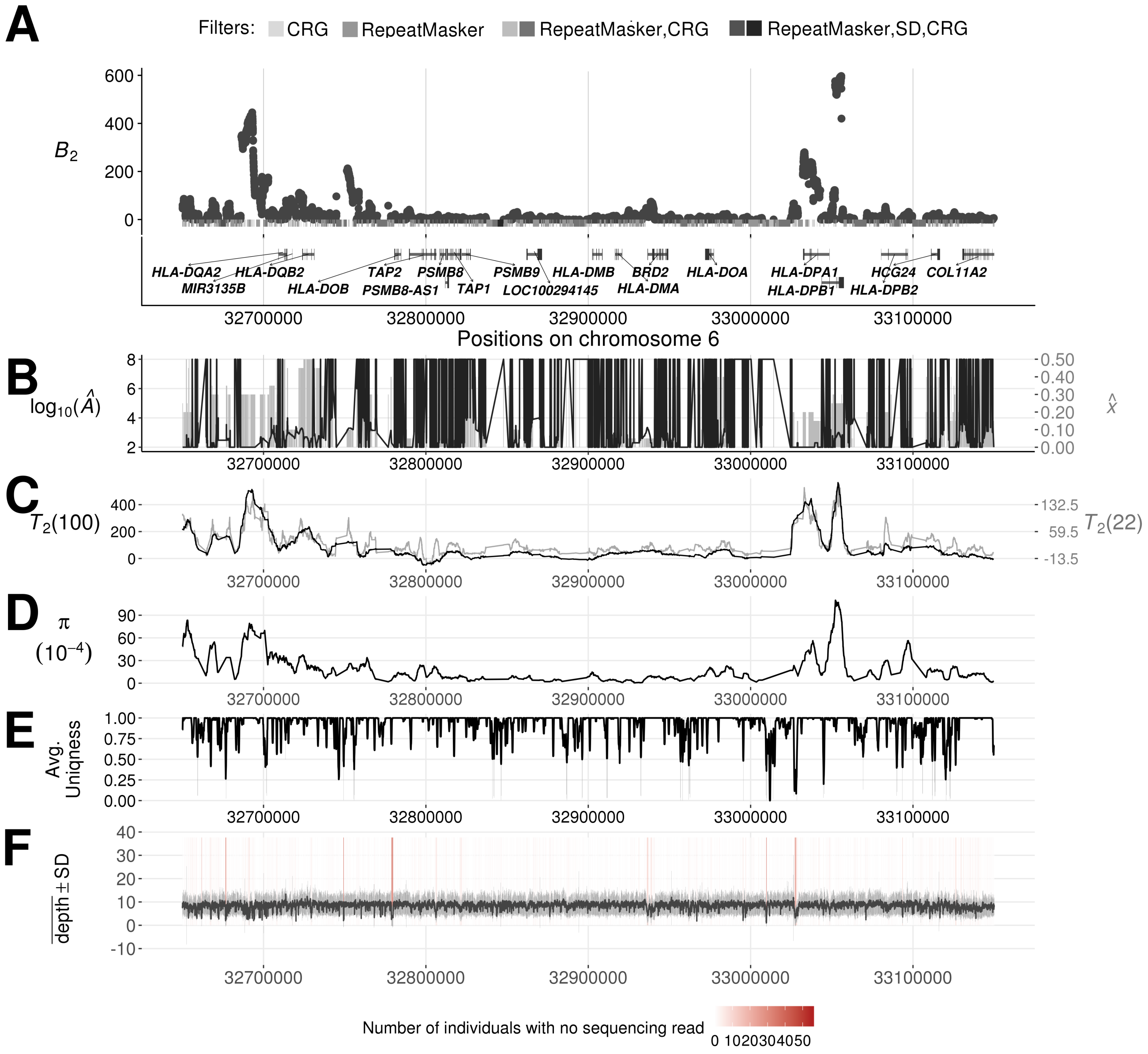


Figure S15: Evidence of balancing selection and sequencing quality around the MHC-D (HLA-D) region in the CEU population. (A) $B_2$ scores across the region, with gray bars indicating the regions removed by filters. (B) The optimal equilibrium frequency $\widehat{x}$ (gray bars) and $\log_{10}(\widehat{A})$ (black lines) for each maximized likelihood ratio. (C) $T_2$ scores across the region, with the black and gray lines respectively stand for the scans using 100 and 22 informative sites on either side of each test site. (D) Nucleotide diversity $\pi$ computed from every window of length five kb across the region. (E) The 35 nucleotide sequence uniqueness, indicating mappability, averaged over windows of length 500 nucleotides across the region.

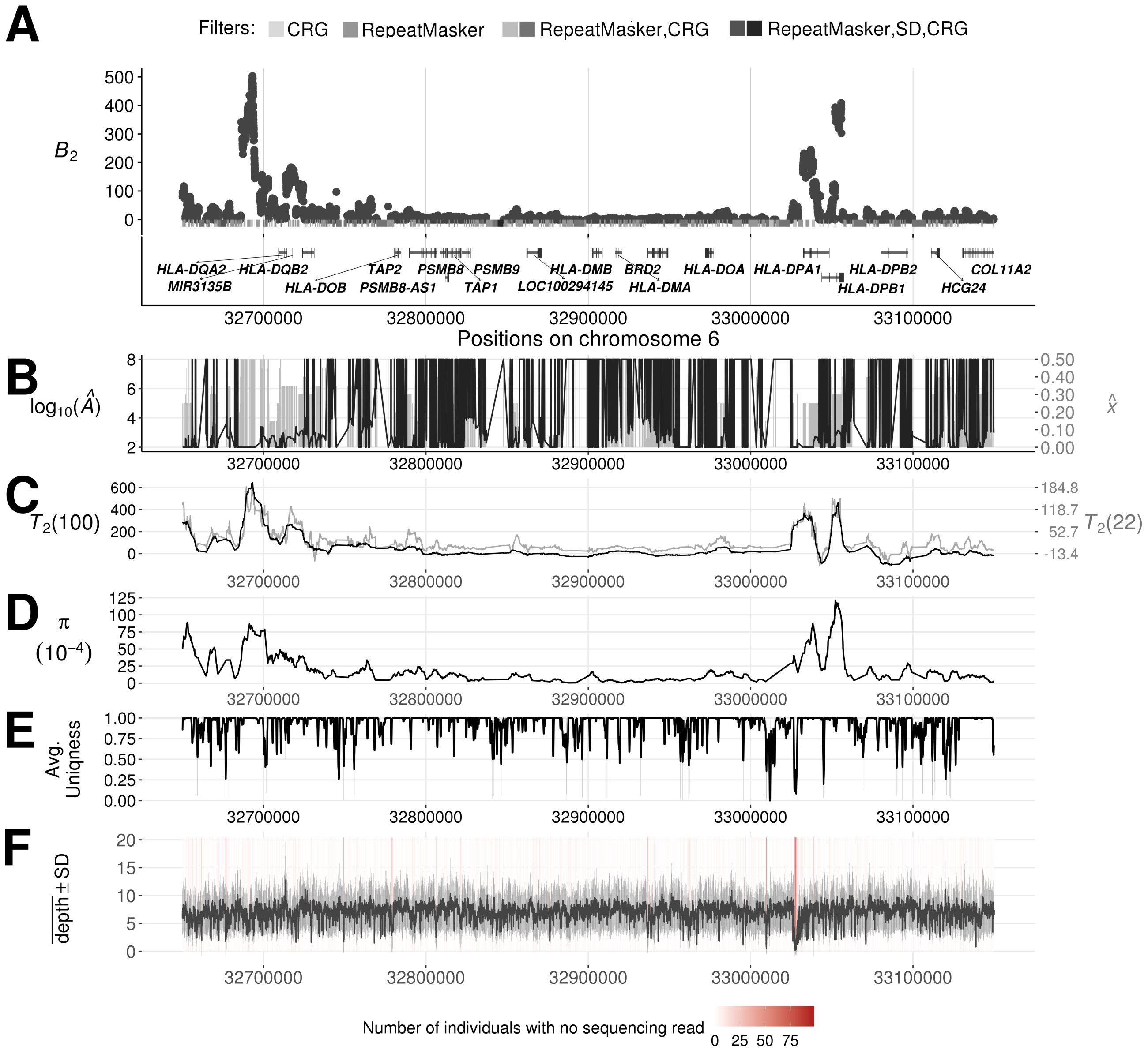


Figure S16: Evidence of balancing selection and sequencing quality around the MHC-D (HLA-D) region in the YRI population. (A) $B_2$ scores across the region, with gray bars indicating the regions removed by filters. (B) The optimal equilibrium frequency $\widehat{x}$ (gray bars) and $\log_{10}(\widehat{A})$ (black lines) for each maximized likelihood ratio. (C) $T_2$ scores across the region, with the black and gray lines respectively stand for the scans using 100 and 22 informative sites on either side of each test site. (D) Nucleotide diversity $\pi$ computed from every window of length five kb across the region. (E) The 35 nucleotide sequence uniqueness, indicating mappability, averaged over windows of length 500 nucleotides across the region.

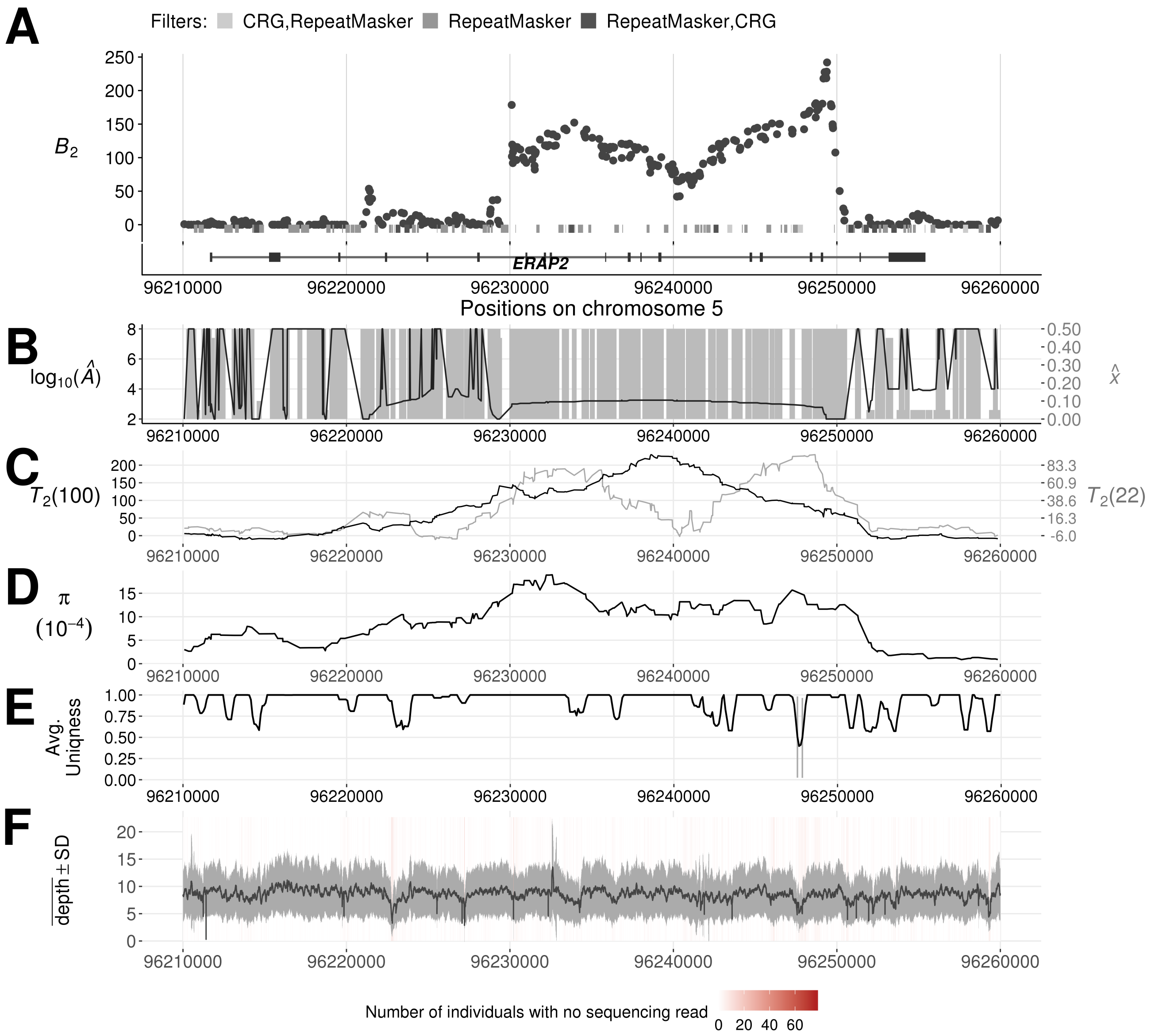


Figure S17: Evidence of balancing selection and sequencing quality across the *ERAP2* gene in the CEU population. (A) $B_2$ scores across the region, with gray bars indicating the regions removed by filters. (B) The optimal equilibrium frequency $\widehat{x}$ (gray bars) and $\log_{10}(\widehat{A})$ (black lines) for each maximized likelihood ratio. (C) $T_2$ scores across the region, with the black and gray lines respectively stand for the scans using 100 and 22 informative sites on either side of each test site. (D) Nucleotide diversity $\pi$ computed from every window of length five kb across the region. (E) The 35 nucleotide sequence uniqueness, indicating mappability, averaged over windows of length 500 nucleotides across the region.

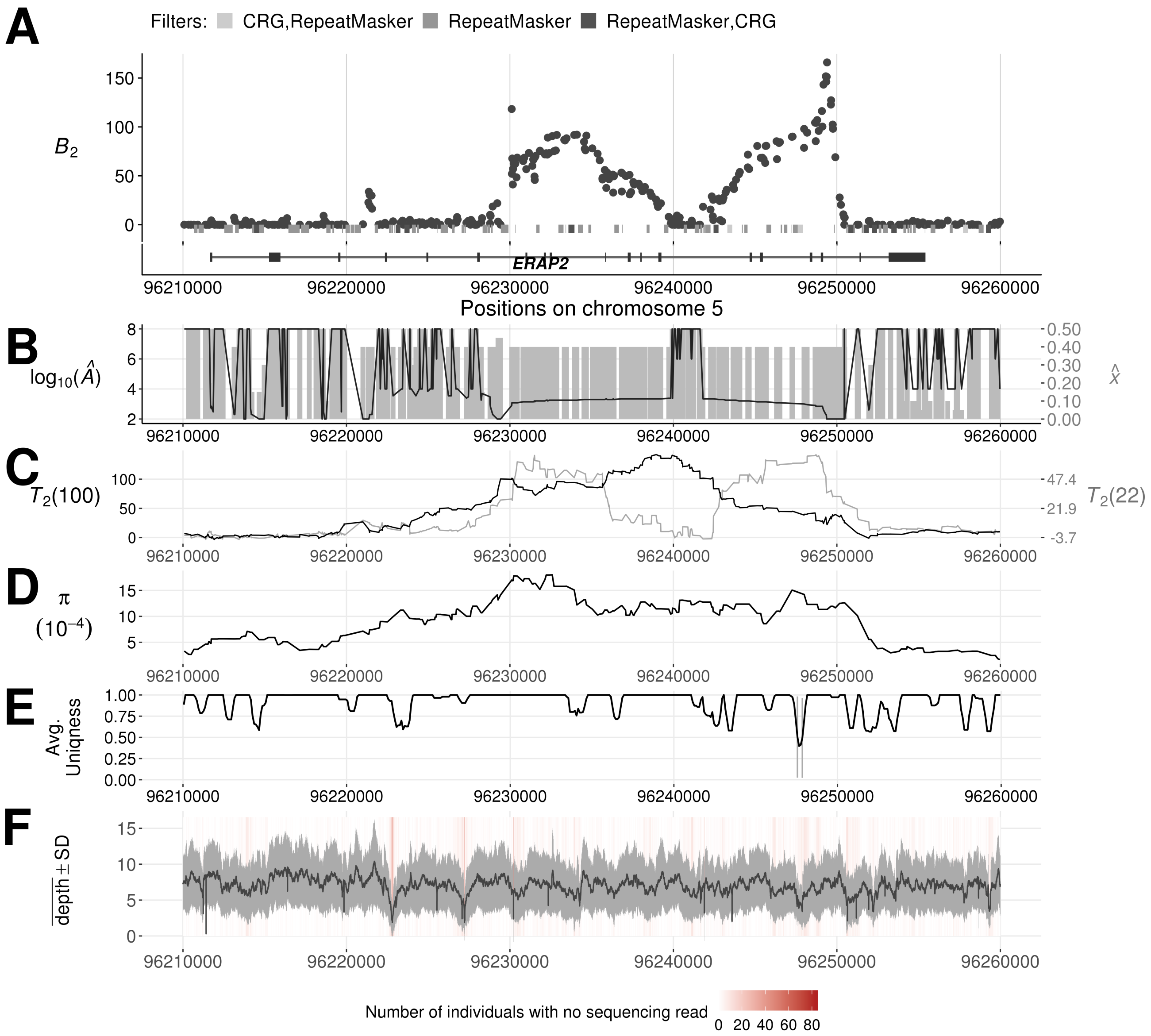


Figure S18: Evidence of balancing selection and sequencing quality across the *ERAP2* gene in the YRI population. (A) $B_2$ scores across the region, with gray bars indicating the regions removed by filters. (B) The optimal equilibrium frequency $\widehat{x}$ (gray bars) and $\log_{10}(\widehat{A})$ (black lines) for each maximized likelihood ratio. (C) $T_2$ scores across the region, with the black and gray lines respectively stand for the scans using 100 and 22 informative sites on either side of each test site. (D) Nucleotide diversity $\pi$ computed from every window of length five kb across the region. (E) The 35 nucleotide sequence uniqueness, indicating mappability, averaged over windows of length 500 nucleotides across the region.

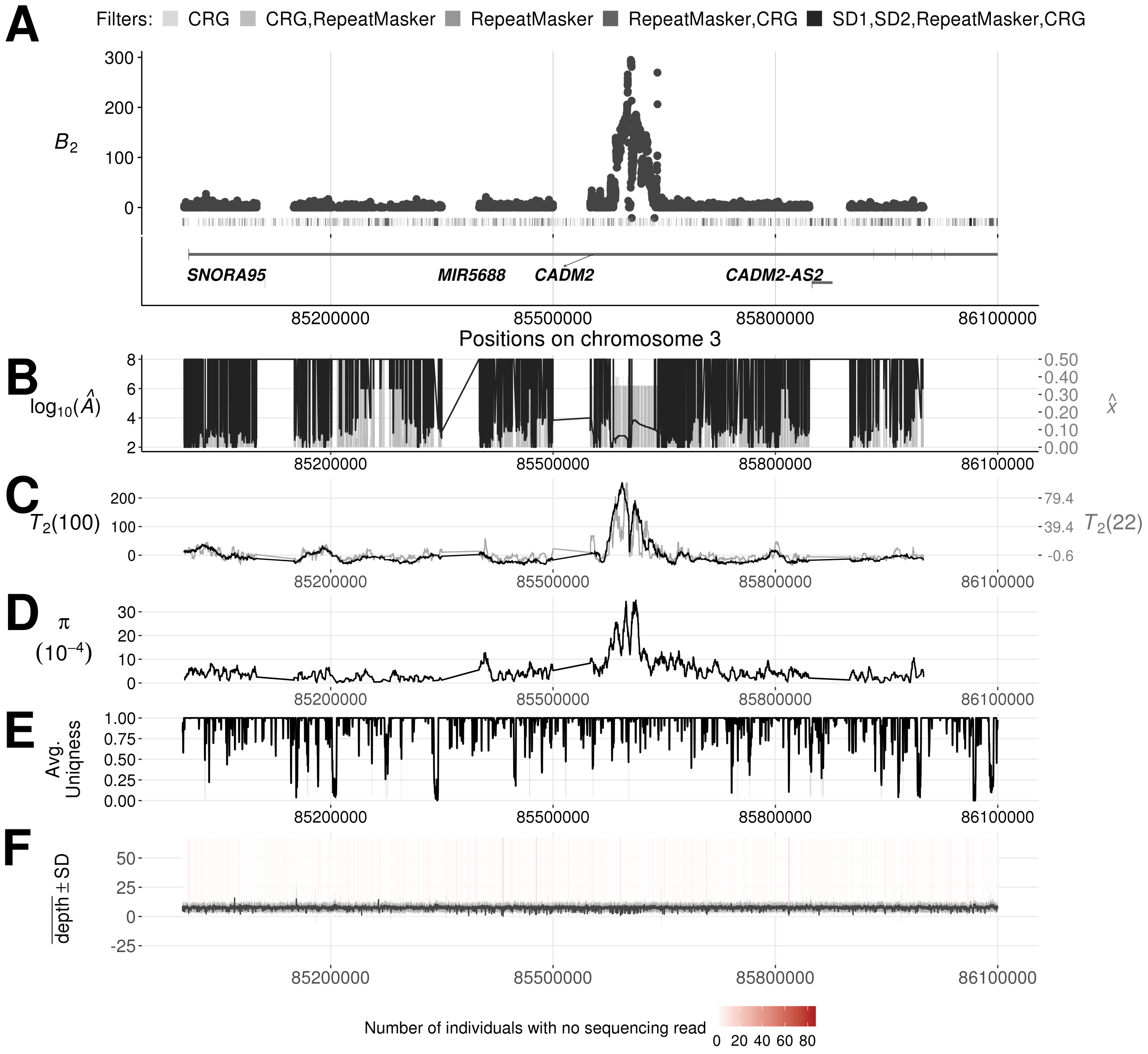


Figure S19: Evidence of balancing selection and sequencing quality across the *CADM2* gene in the CEU population. (A) $B_2$ scores across the region, with gray bars indicating the regions removed by filters. (B) The optimal equilibrium frequency $\widehat{x}$ (gray bars) and $\log_{10}(\widehat{A})$ (black lines) for each maximized likelihood ratio. (C) $T_2$ scores across the region, with the black and gray lines respectively stand for the scans using 100 and 22 informative sites on either side of each test site. (D) Nucleotide diversity $\pi$ computed from every window of length five kb across the region. (E) The 35 nucleotide sequence uniqueness, indicating mappability, averaged over windows of length 500 nucleotides across the region.

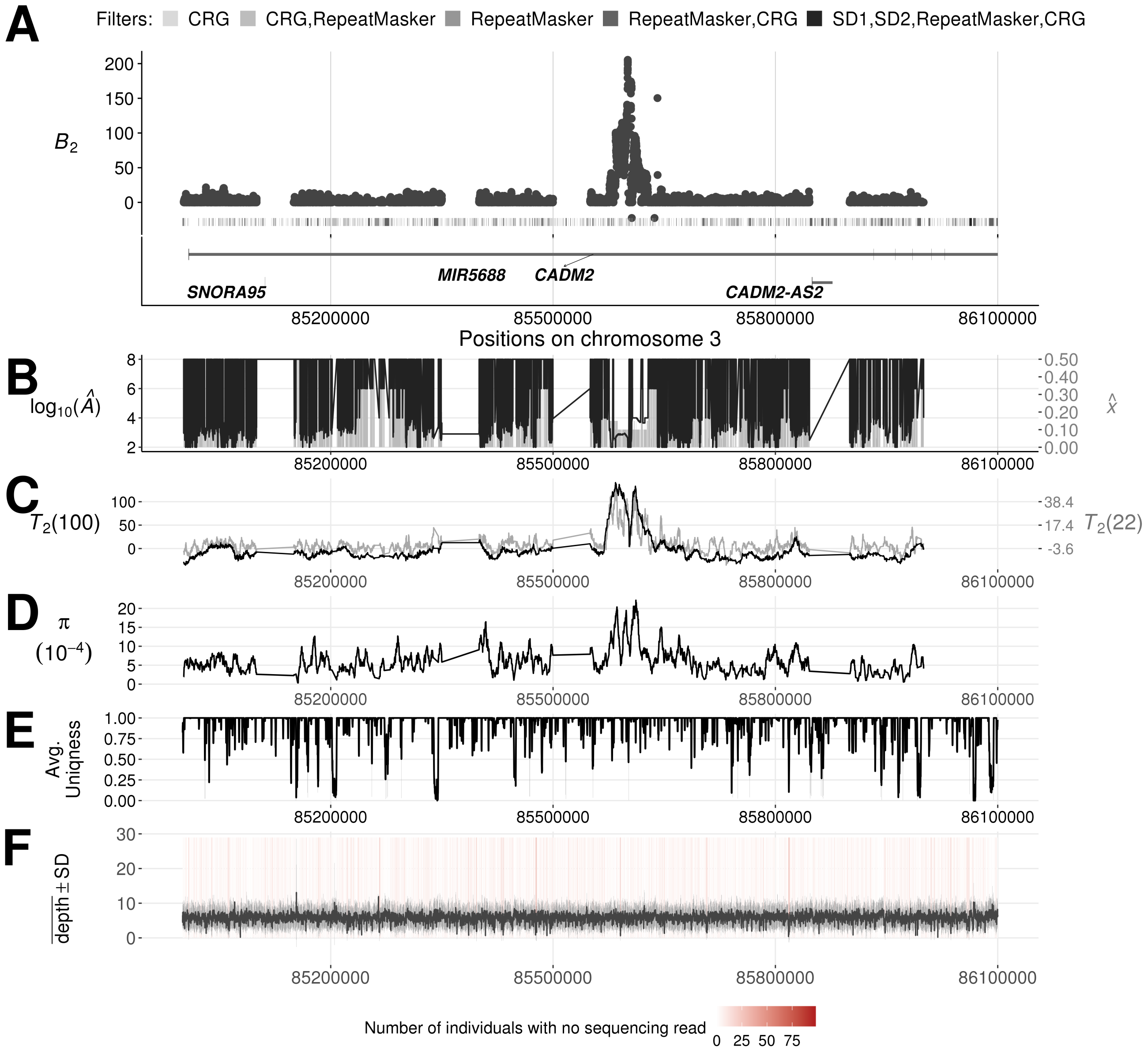


Figure S20: Evidence of balancing selection and sequencing quality across the *CADM2* gene in the YRI population. (A) $B_2$ scores across the region, with gray bars indicating the regions removed by filters. (B) The optimal equilibrium frequency $\widehat{x}$ (gray bars) and $\log_{10}(\widehat{A})$ (black lines) for each maximized likelihood ratio. (C) $T_2$ scores across the region, with the black and gray lines respectively stand for the scans using 100 and 22 informative sites on either side of each test site. (D) Nucleotide diversity $\pi$ computed from every window of length five kb across the region. (E) The 35 nucleotide sequence uniqueness, indicating mappability, averaged over windows of length 500 nucleotides across the region.

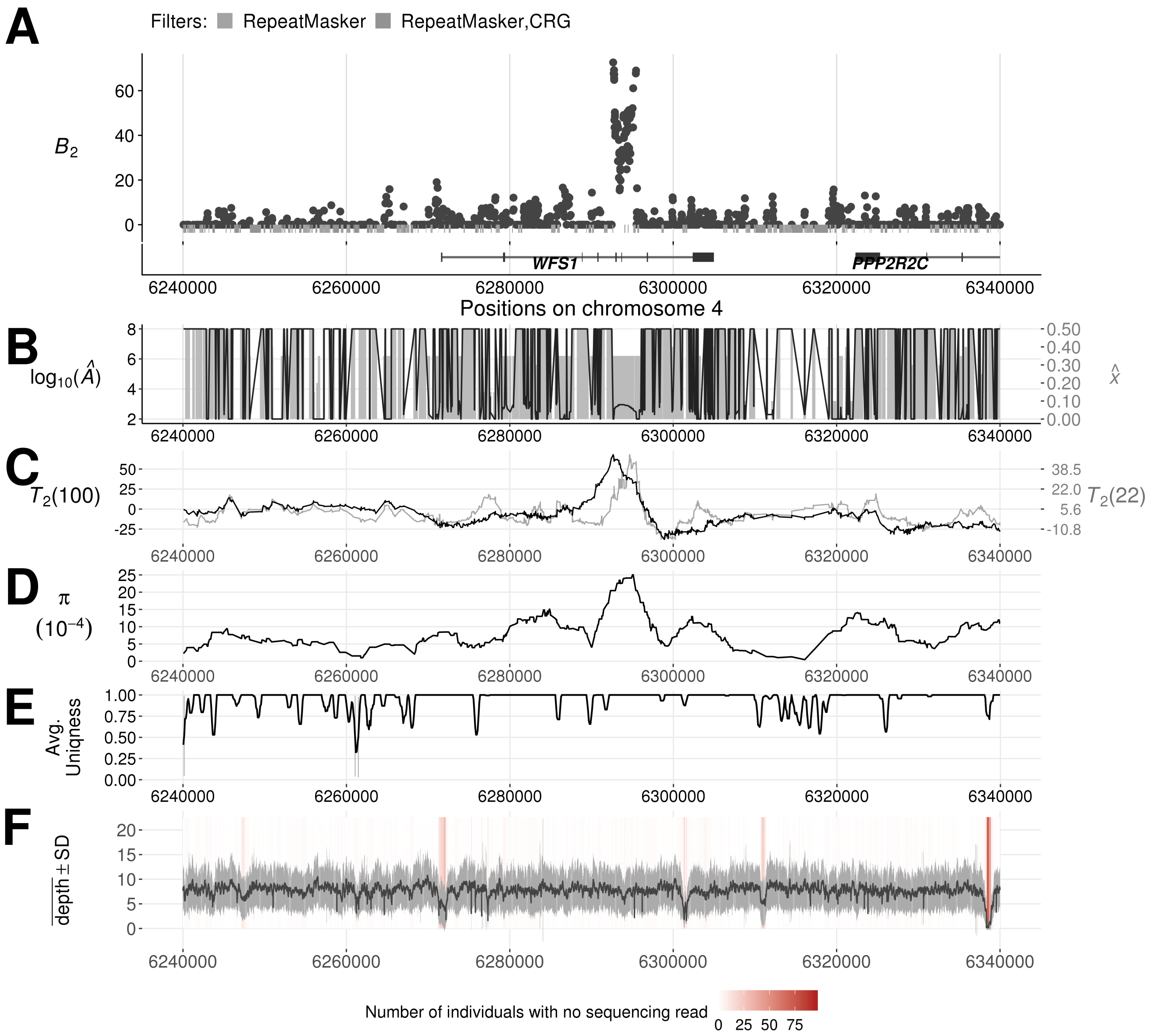


Figure S21: Evidence of balancing selection and sequencing quality around the *WFS1* gene in the CEU population. (A) $B_2$ scores across the region, with gray bars indicating the regions removed by filters. (B) The optimal equilibrium frequency $\widehat{x}$ (gray bars) and $\log_{10}(\widehat{A})$ (black lines) for each maximized likelihood ratio. (C) $T_2$ scores across the region, with the black and gray lines respectively stand for the scans using 100 and 22 informative sites on either side of each test site. (D) Nucleotide diversity $\pi$ computed from every window of length five kb across the region. (E) The 35 nucleotide sequence uniqueness, indicating mappability, averaged over windows of length 500 nucleotides across the region.

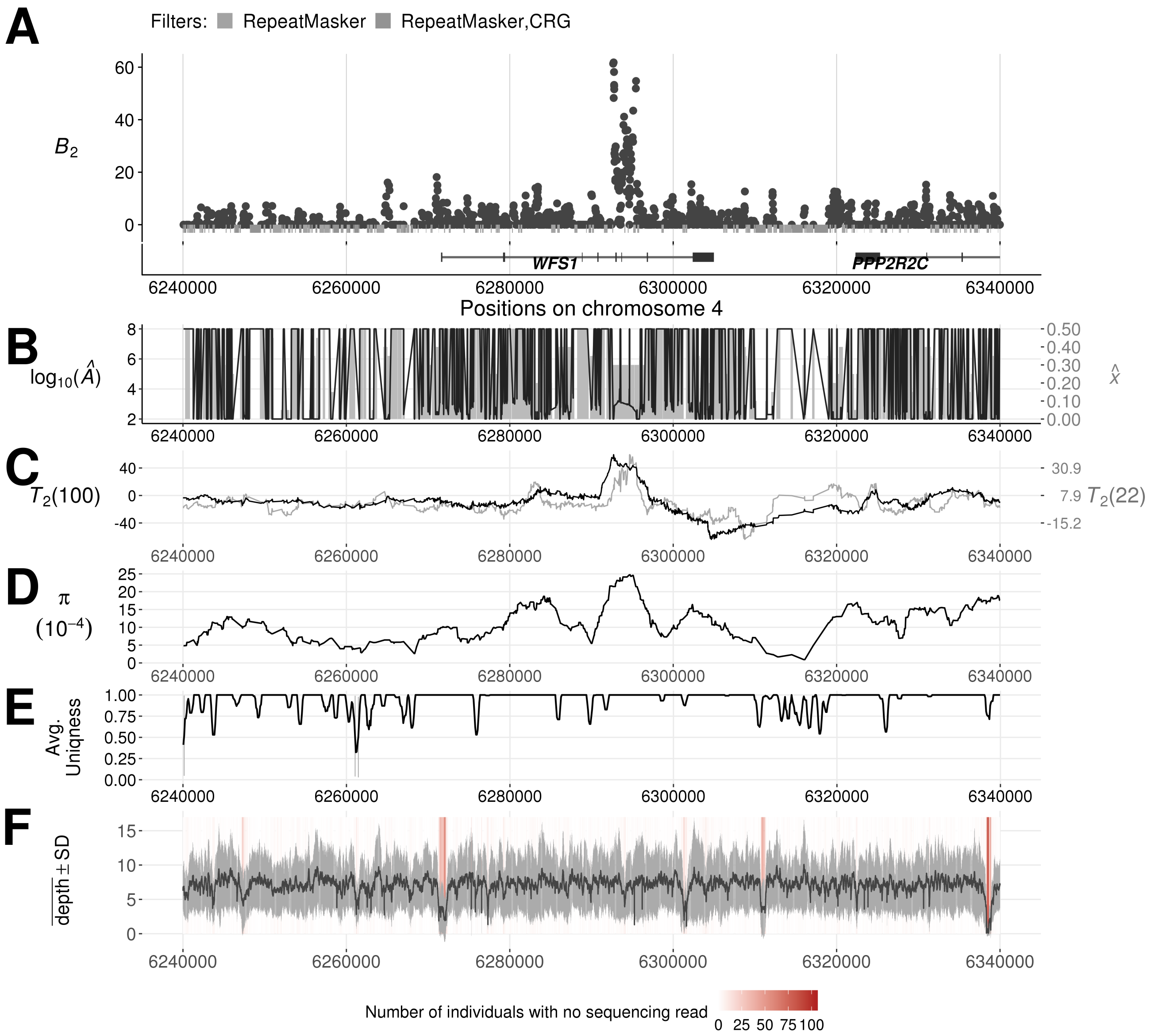


Figure S22: Evidence of balancing selection and sequencing quality around the *WFS1* gene in the YRI population. (A) $B_2$ scores across the region, with gray bars indicating the regions removed by filters. (B) The optimal equilibrium frequency $\widehat{x}$ (gray bars) and $\log_{10}(\widehat{A})$ (black lines) for each maximized likelihood ratio. (C) $T_2$ scores across the region, with the black and gray lines respectively stand for the scans using 100 and 22 informative sites on either side of each test site. (D) Nucleotide diversity $\pi$ computed from every window of length five kb across the region. (E) The 35 nucleotide sequence uniqueness, indicating mappability, averaged over windows of length 500 nucleotides across the region.

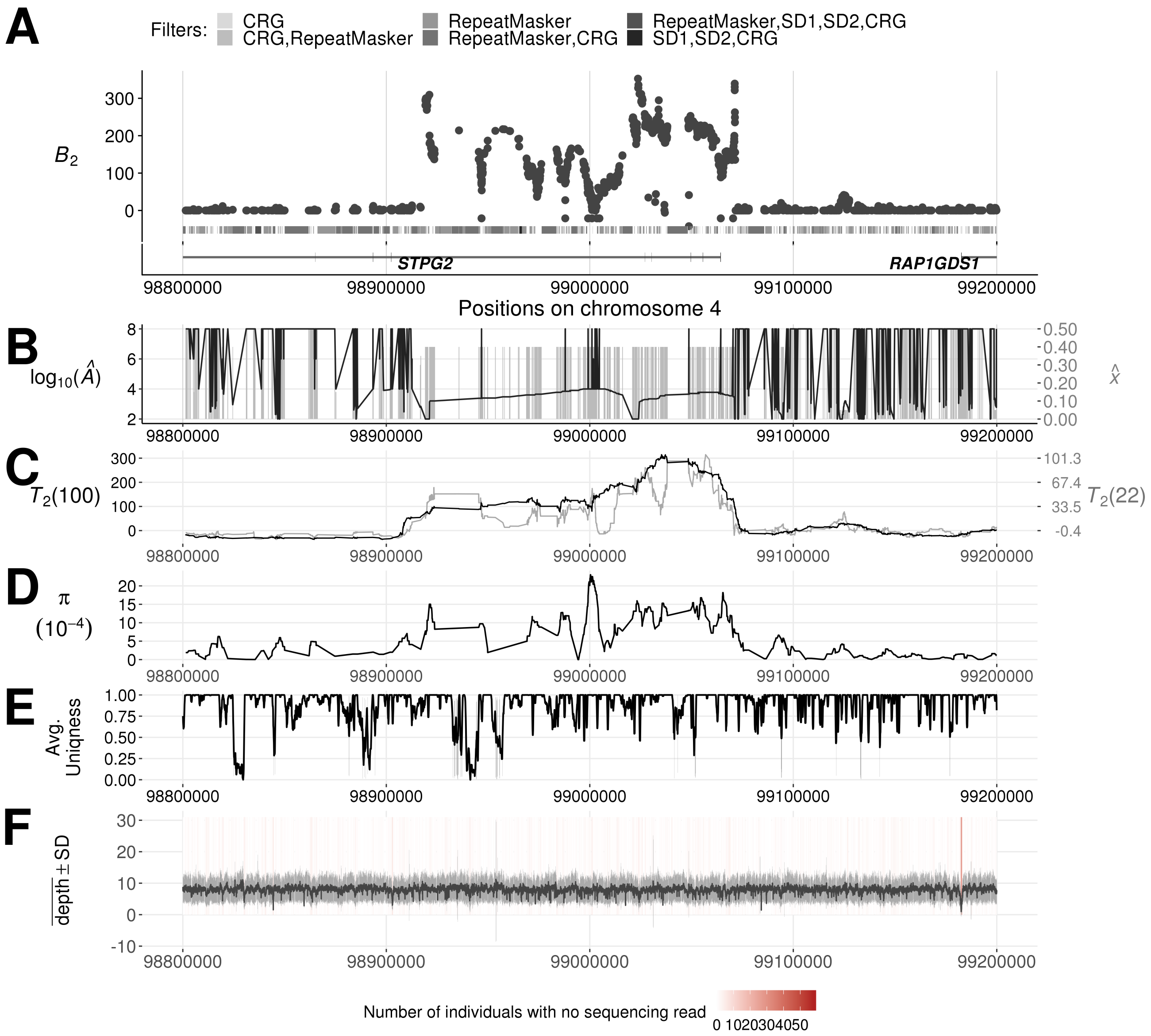


Figure S23: Evidence of balancing selection and sequencing quality around the *STPG2* gene in the CEU population. (A) $B_2$ scores across the region, with gray bars indicating the regions removed by filters. (B) The optimal equilibrium frequency $\widehat{x}$ (gray bars) and $\log_{10}(\widehat{A})$ (black lines) for each maximized likelihood ratio. (C) $T_2$ scores across the region, with the black and gray lines respectively stand for the scans using 100 and 22 informative sites on either side of each test site. (D) Nucleotide diversity $\pi$ computed from every window of length five kb across the region. (E) The 35 nucleotide sequence uniqueness, indicating mappability, averaged over windows of length 500 nucleotides across the region.

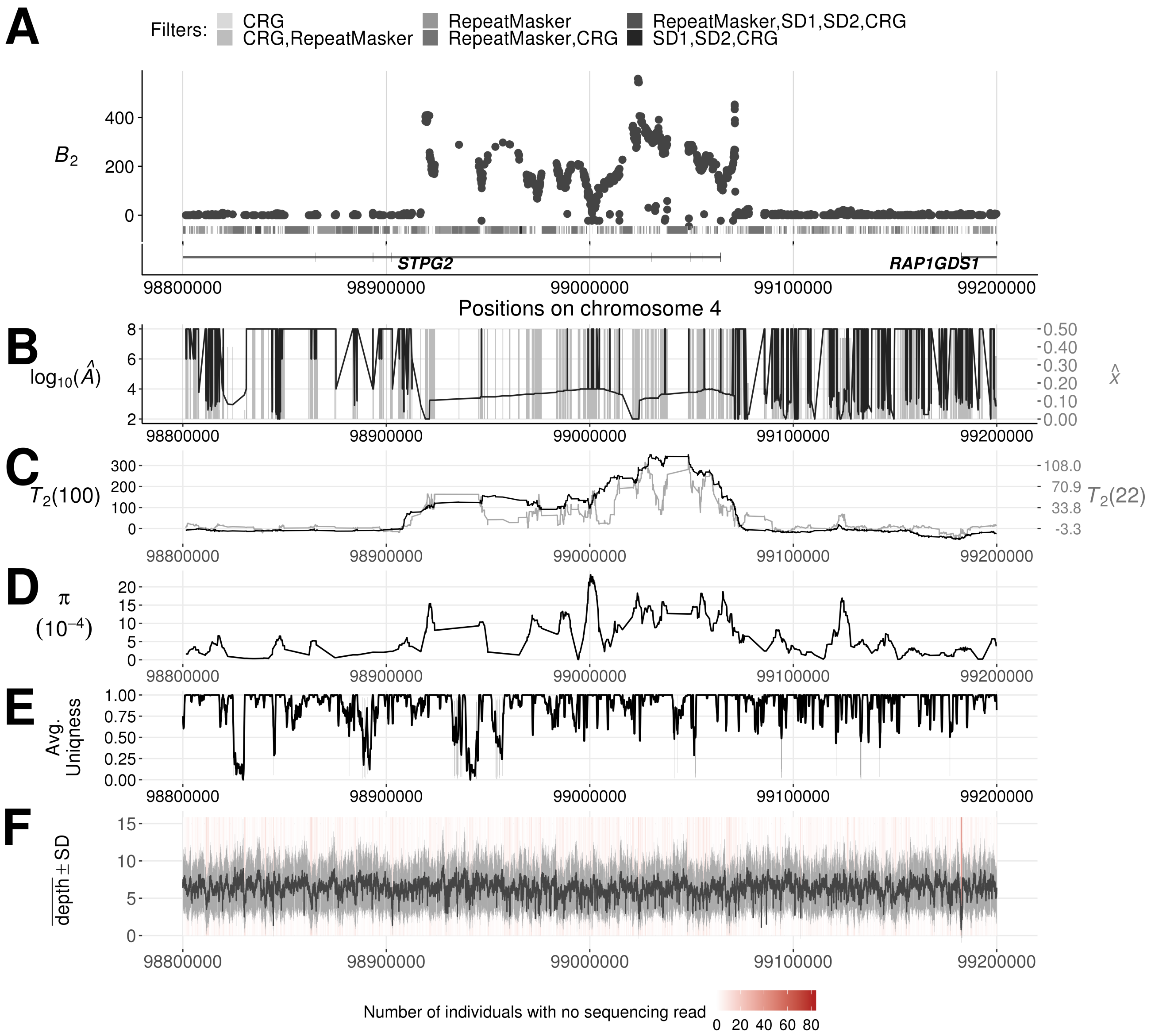


Figure S24: Evidence of balancing selection and sequencing quality around the *STPG2* gene in the YRI population. (A) $B_2$ scores across the region, with gray bars indicating the regions removed by filters. (B) The optimal equilibrium frequency $\widehat{x}$ (gray bars) and $\log_{10}(\widehat{A})$ (black lines) for each maximized likelihood ratio. (C) $T_2$ scores across the region, with the black and gray lines respectively stand for the scans using 100 and 22 informative sites on either side of each test site. (D) Nucleotide diversity $\pi$ computed from every window of length five kb across the region. (E) The 35 nucleotide sequence uniqueness, indicating mappability, averaged over windows of length 500 nucleotides across the region.

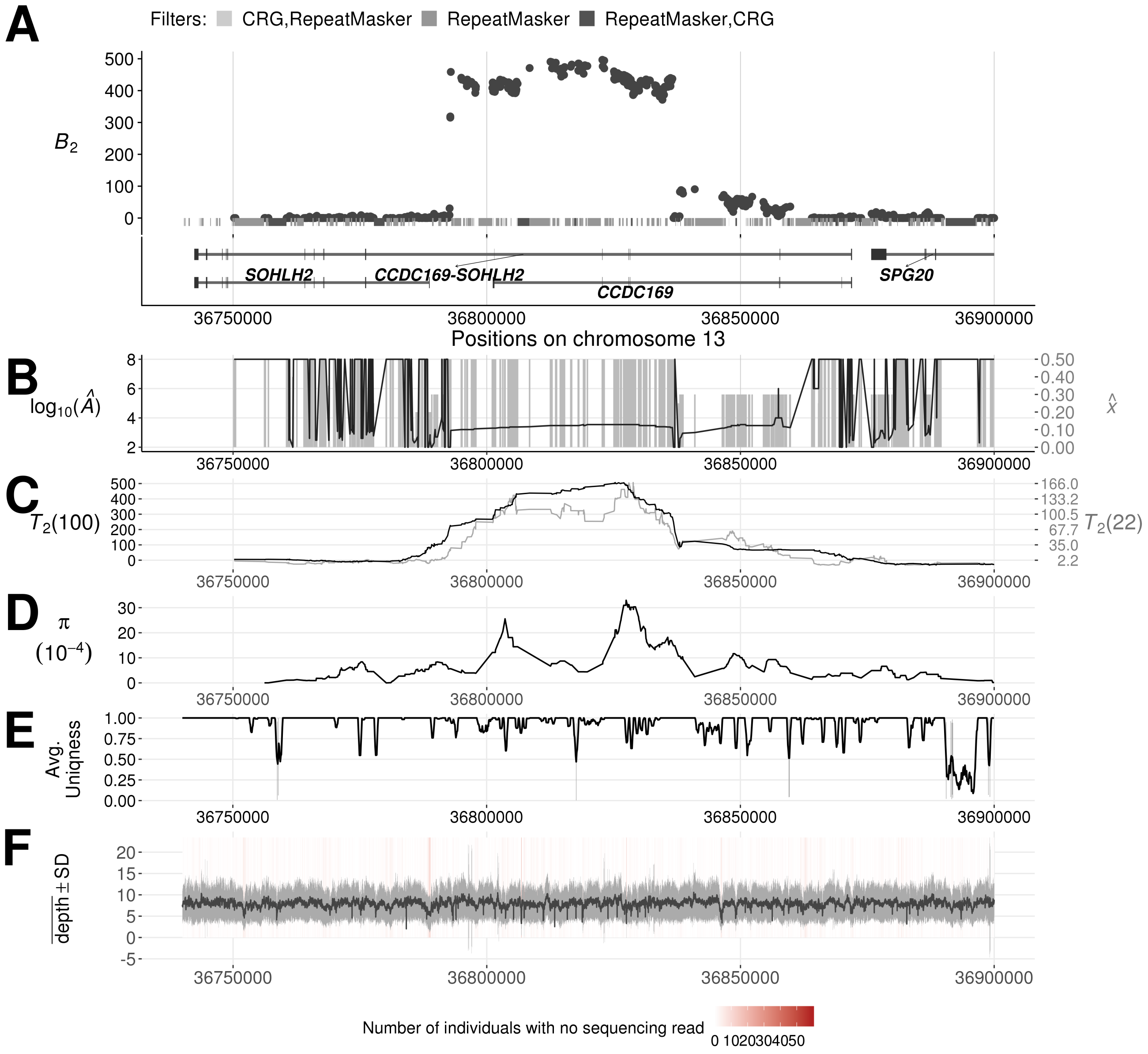


Figure S25: Evidence of balancing selection and sequencing quality across the *CCDC169-SOHLH2* region in the CEU population. (A) $B_2$ scores across the region, with gray bars indicating the regions removed by filters. (B) The optimal equilibrium frequency $\widehat{x}$ (gray bars) and $\log_{10}(\widehat{A})$ (black lines) for each maximized likelihood ratio. (C) $T_2$ scores across the region, with the black and gray lines respectively stand for the scans using 100 and 22 informative sites on either side of each test site. (D) Nucleotide diversity $\pi$ computed from every window of length five kb across the region. (E) The 35 nucleotide sequence uniqueness, indicating mappability, averaged over windows of length 500 nucleotides across the region.

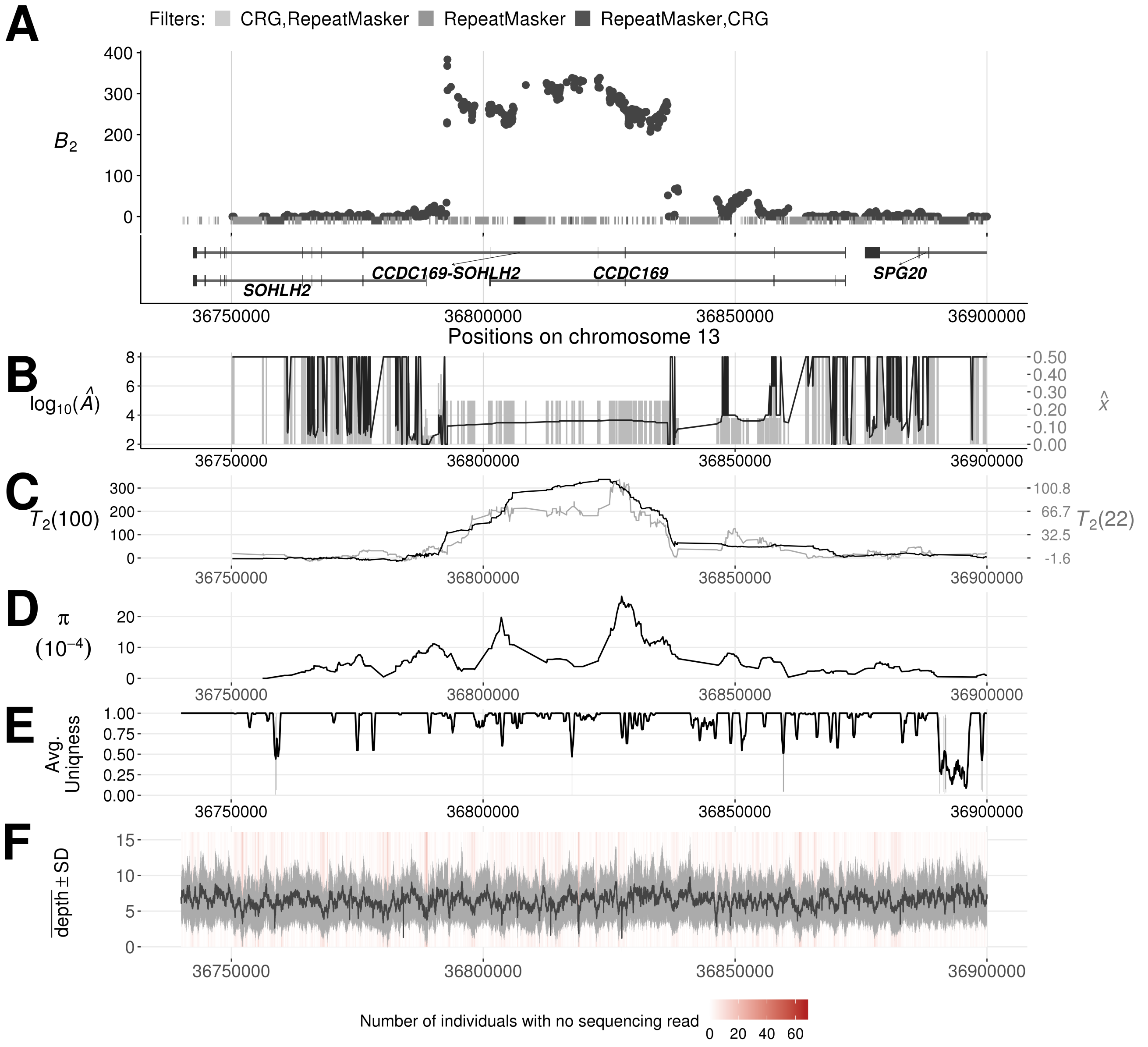


Figure S26: Evidence of balancing selection and sequencing quality across the *CCDC169-SOHLH2* region in the YRI population. (A) $B_2$ scores across the region, with gray bars indicating the regions removed by filters. (B) The optimal equilibrium frequency $\widehat{x}$ (gray bars) and $\log_{10}(\widehat{A})$ (black lines) for each maximized likelihood ratio. (C) $T_2$ scores across the region, with the black and gray lines respectively stand for the scans using 100 and 22 informative sites on either side of each test site. (D) Nucleotide diversity $\pi$ computed from every window of length five kb across the region. (E) The 35 nucleotide sequence uniqueness, indicating mappability, averaged over windows of length 500 nucleotides across the region.

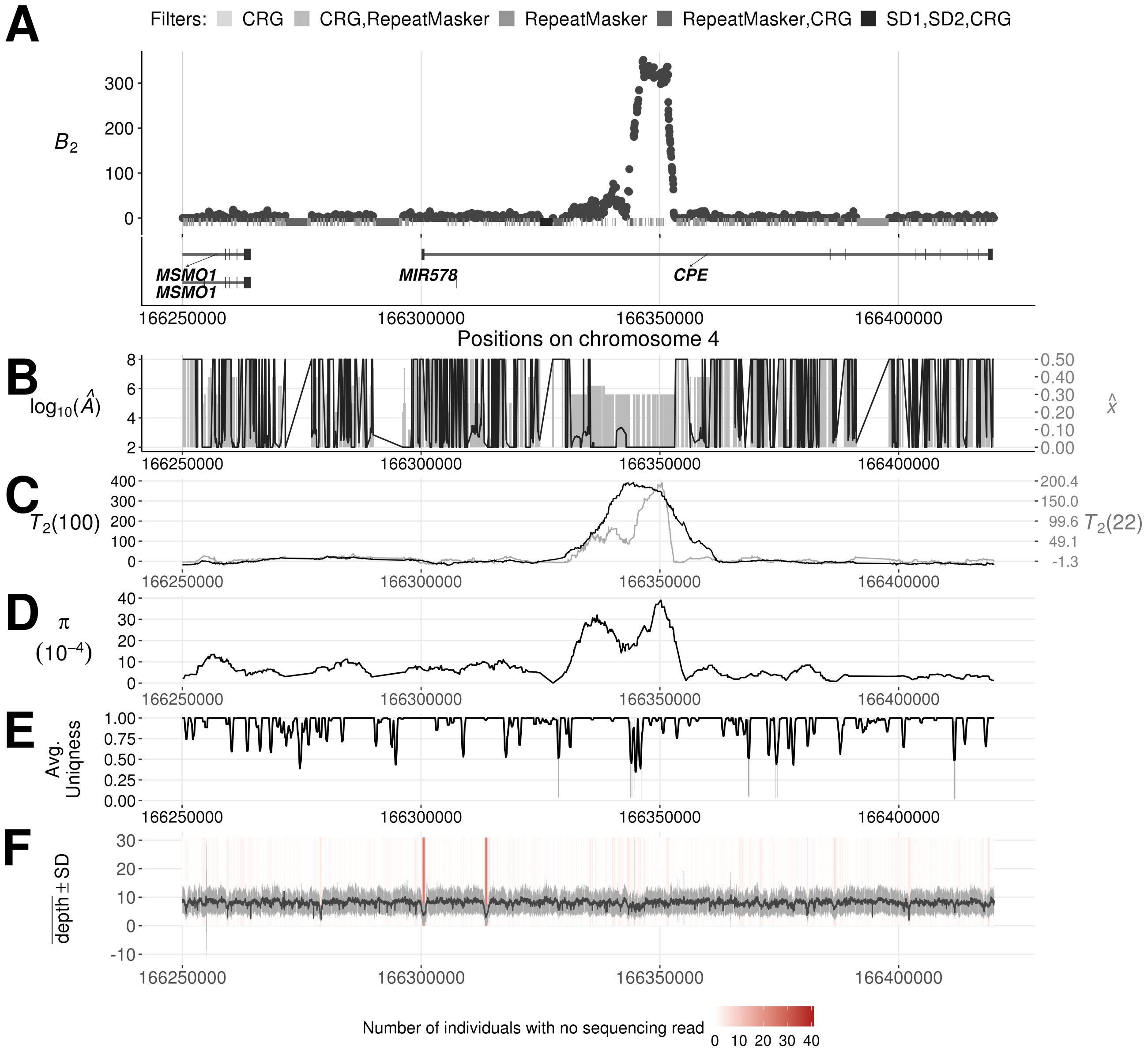


Figure S27: Evidence of balancing selection and sequencing quality around the *MYOM2* gene in the CEU population. (A) $B_2$ scores across the region, with gray bars indicating the regions removed by filters. (B) The optimal equilibrium frequency $\widehat{x}$ (gray bars) and $\log_{10}(\widehat{A})$ (black lines) for each maximized likelihood ratio. (C) $T_2$ scores across the region, with the black and gray lines respectively stand for the scans using 100 and 22 informative sites on either side of each test site. (D) Nucleotide diversity $\pi$ computed from every window of length five kb across the region. (E) The 35 nucleotide sequence uniqueness, indicating mappability, averaged over windows of length 500 nucleotides across the region.

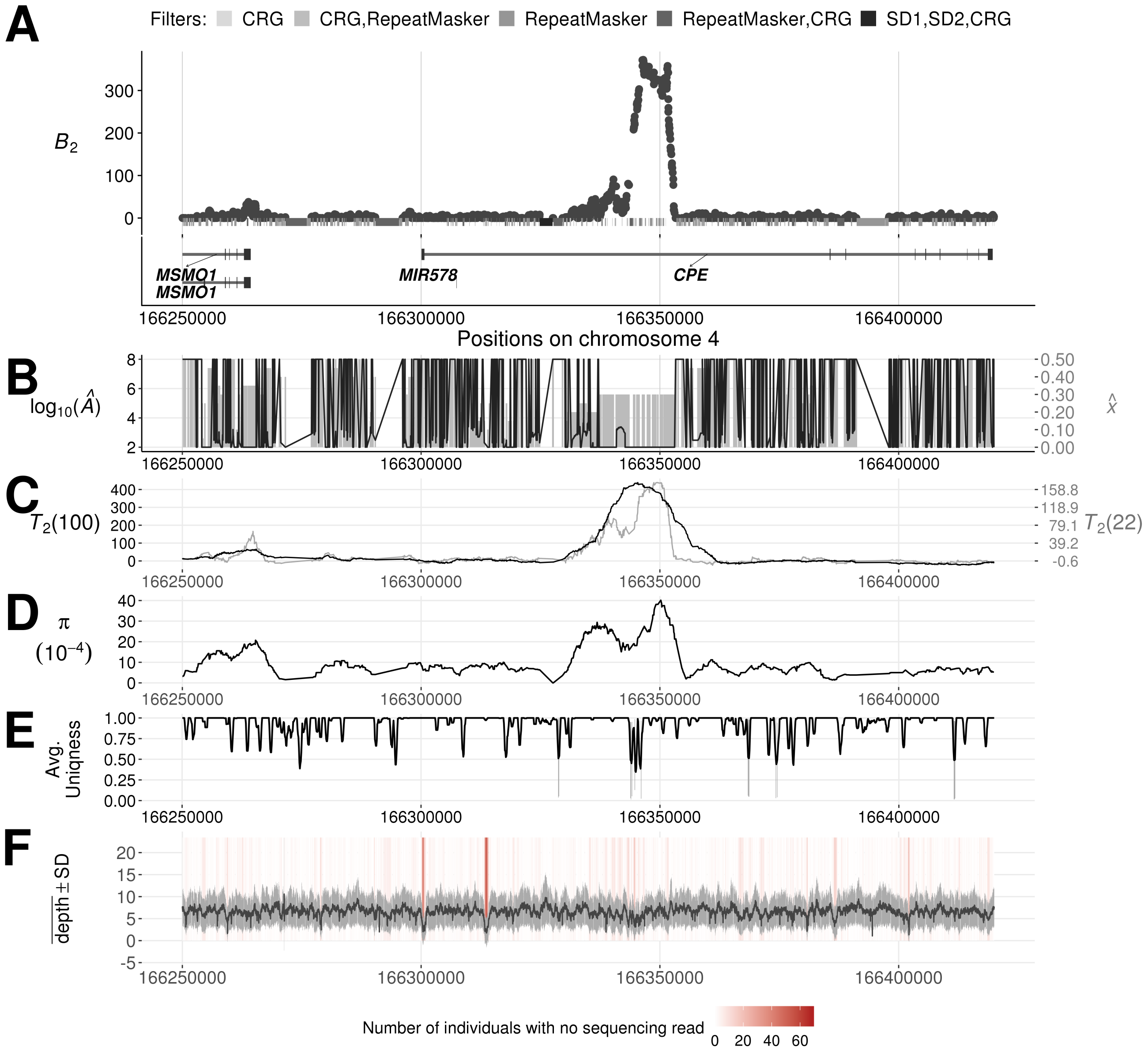


Figure S28: Evidence of balancing selection and sequencing quality around the *CPE* gene in the YRI population. (A) $B_2$ scores across the region, with gray bars indicating the regions removed by filters. (B) The optimal equilibrium frequency $\widehat{x}$ (gray bars) and $\log_{10}(\widehat{A})$ (black lines) for each maximized likelihood ratio. (C) $T_2$ scores across the region, with the black and gray lines respectively stand for the scans using 100 and 22 informative sites on either side of each test site. (D) Nucleotide diversity $\pi$ computed from every window of length five kb across the region. (E) The 35 nucleotide sequence uniqueness, indicating mappability, averaged over windows of length 500 nucleotides across the region.

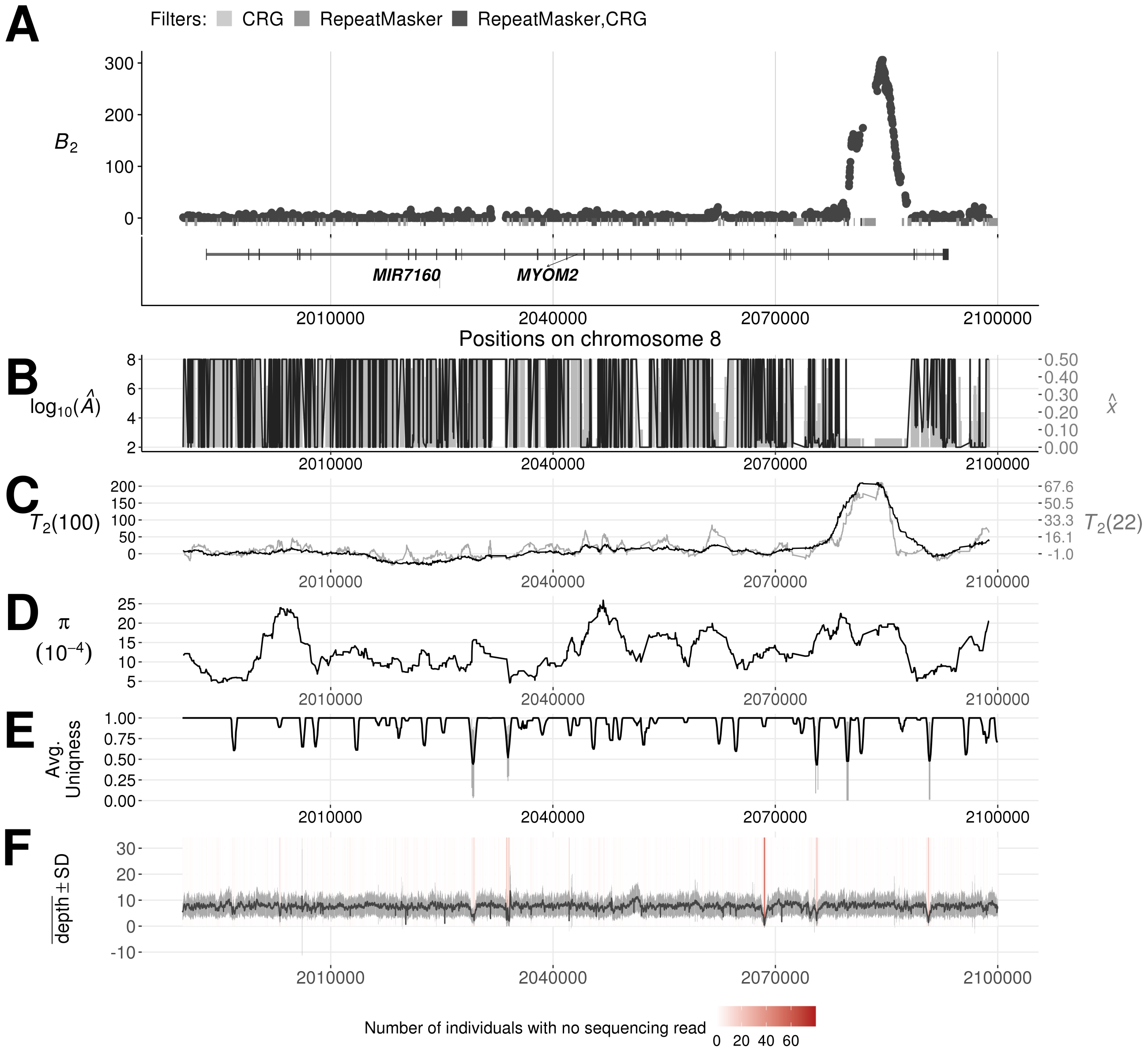


Figure S29: Evidence of balancing selection and sequencing quality around the *CPE* gene in the CEU population. (A) $B_2$ scores across the region, with gray bars indicating the regions removed by filters. (B) The optimal equilibrium frequency $\widehat{x}$ (gray bars) and $\log_{10}(\widehat{A})$ (black lines) for each maximized likelihood ratio. (C) $T_2$ scores across the region, with the black and gray lines respectively stand for the scans using 100 and 22 informative sites on either side of each test site. (D) Nucleotide diversity $\pi$ computed from every window of length five kb across the region. (E) The 35 nucleotide sequence uniqueness, indicating mappability, averaged over windows of length 500 nucleotides across the region.

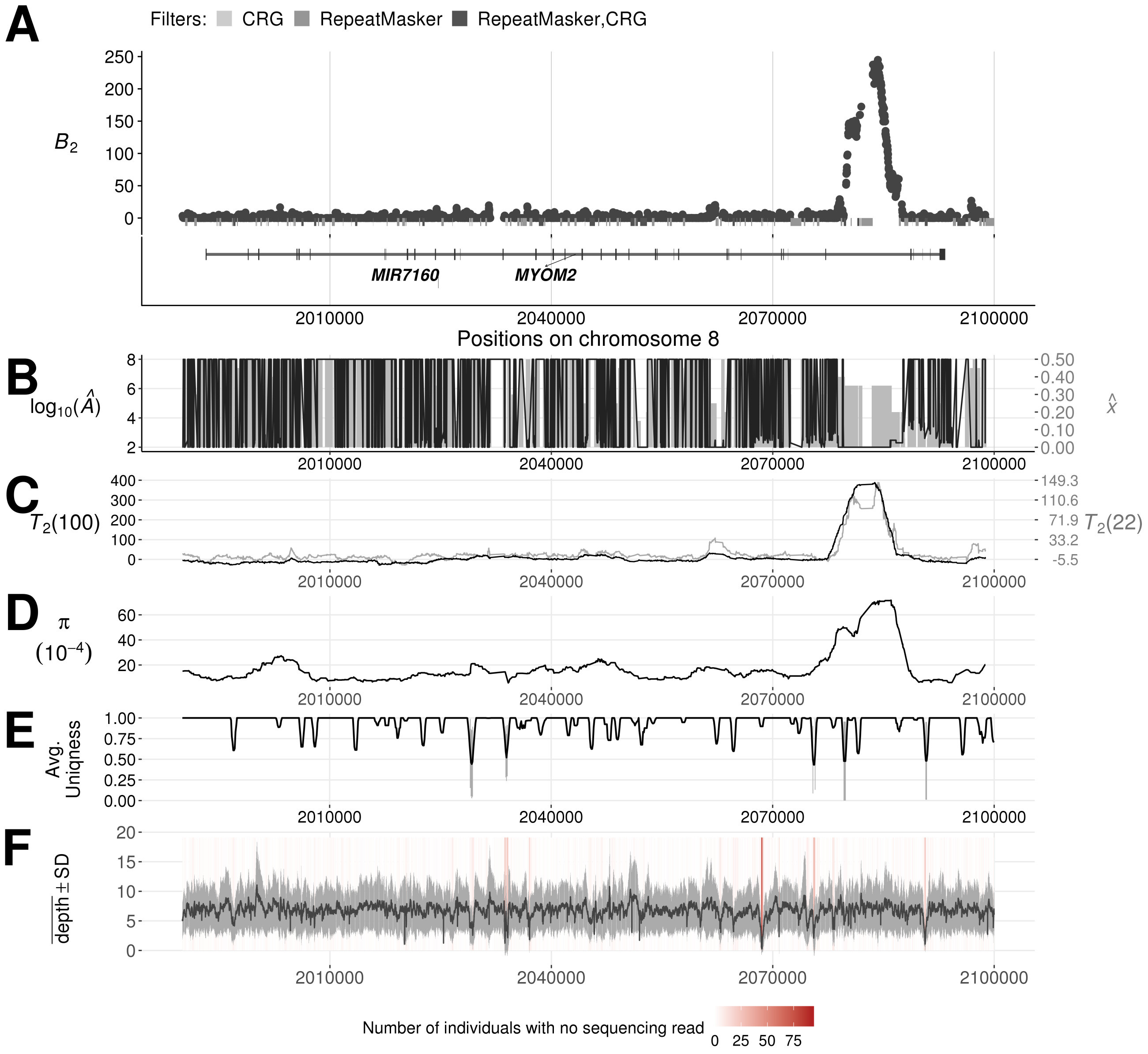


Figure S30: Evidence of balancing selection and sequencing quality around the *MYOM2* gene in the YRI population. (A) $B_2$ scores across the region, with gray bars indicating the regions removed by filters. (B) The optimal equilibrium frequency $\widehat{x}$ (gray bars) and $\log_{10}(\widehat{A})$ (black lines) for each maximized likelihood ratio. (C) $T_2$ scores across the region, with the black and gray lines respectively stand for the scans using 100 and 22 informative sites on either side of each test site. (D) Nucleotide diversity $\pi$ computed from every window of length five kb across the region. (E) The 35 nucleotide sequence uniqueness, indicating mappability, averaged over windows of length 500 nucleotides across the region.

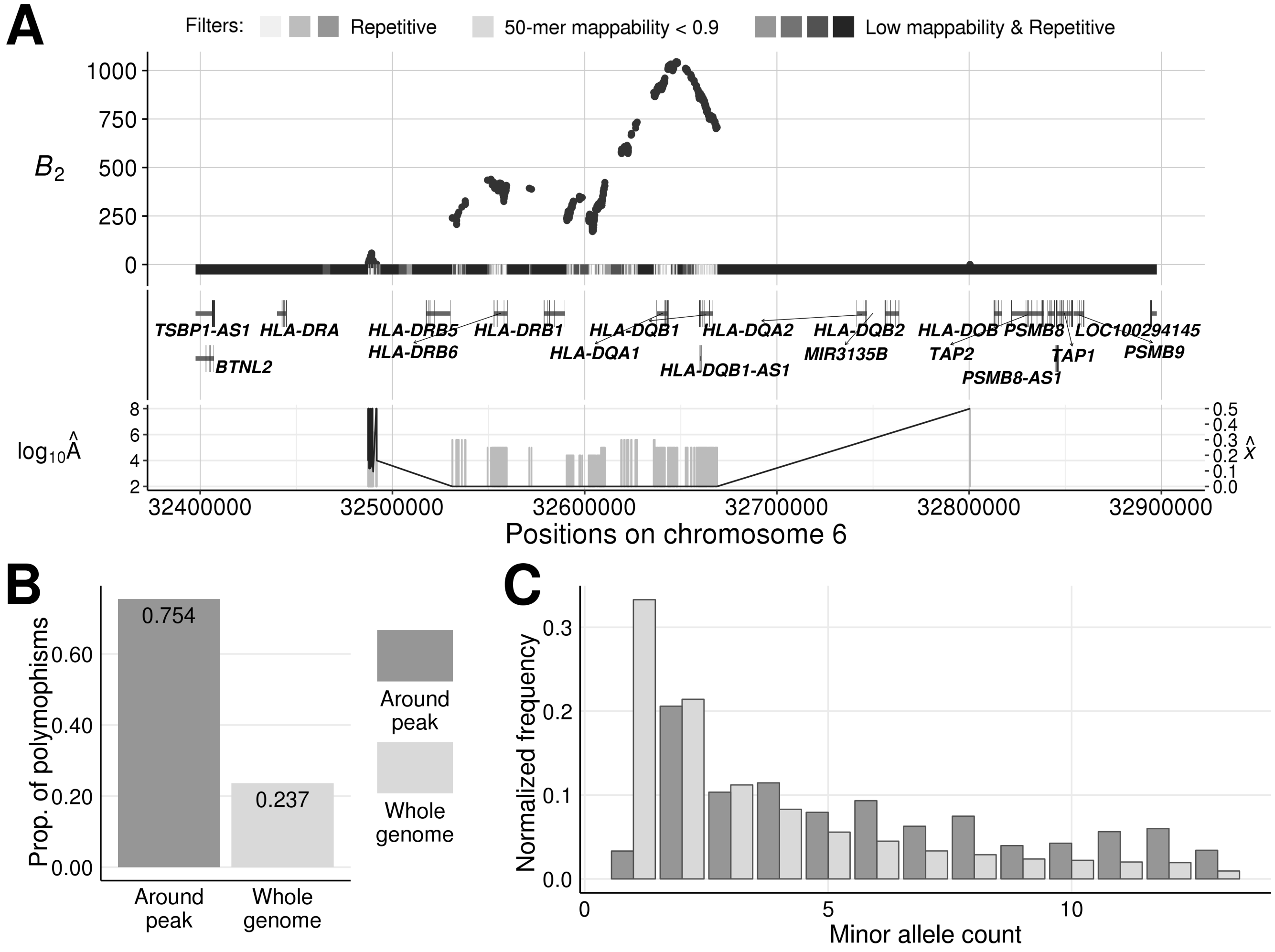


Figure S31: Evidence for balancing selection on *HLA-DQ* genes in bonobos. (A) $B_2$ scores across the genomic region on chromosome 6 surrounding the *HLA-DQA1* and *HLA-DQB1* genes. The gray bars directly under the $B_2$ scores represent the masked regions, as well as the features in these regions. The darker the shade, the greater number of types of repetitive sequences (*e.g.*, RepeatMasker mask, segmental duplication, simple repeats, or interrupted repeats) overlapping the region. Vertical gray bars below display the estimated equilibrium minor allele frequency $\widehat{x}$ for each maximum likelihood ratio $B_2$, and the black line traces the value for the respective inferred footprint size $\log_{10}(\widehat{A})$. (B) Proportion of informative sites that are polymorphic in the 500 kb region centered on the peak compared with the whole-genome average. (C) Minor allele frequency distribution in the 500 kb region centered on the peak compared with the whole-genome average.

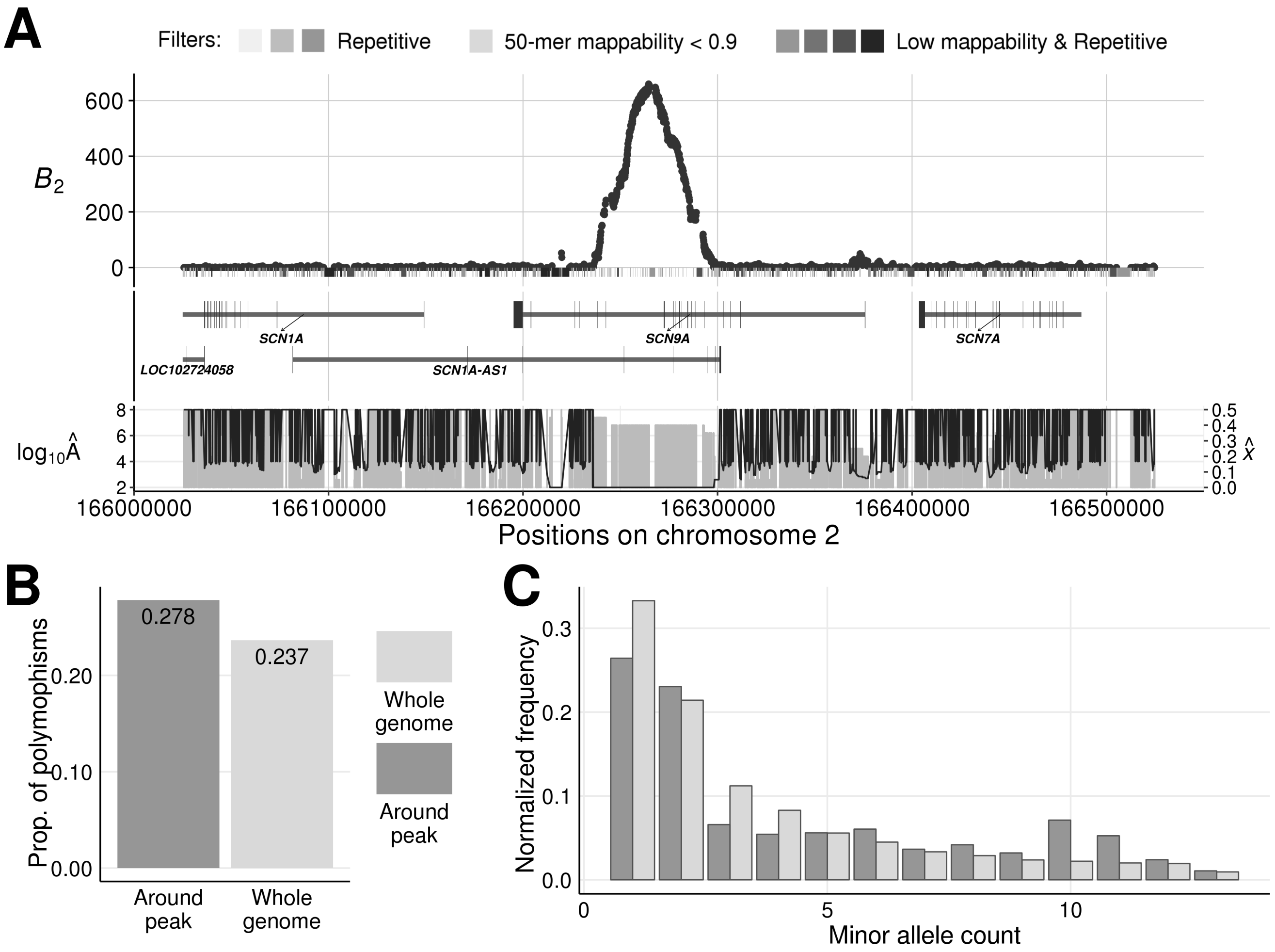


Figure S32: Evidence of balancing selection on *SCN9A*. (A) $B_2$ scores across the 500 kb genomic region on chromosome 2 centered on the peak. The gray bars directly under the $B_2$ scores represent the masked regions, as well as the features in these regions. The darker the shade, the greater number of types of repetitive sequences (*e.g.*, RepeatMasker mask, segmental duplication, simple repeats, or interrupted repeats) overlapping the region. Vertical gray bars below display the estimated equilibrium minor allele frequency $\widehat{x}$ for each maximum likelihood ratio $B_2$, and the black line traces the value for the respective inferred footprint size $\log_{10}(\widehat{A})$. (B) Proportion of informative sites that are polymorphic in the 500 kb region centered on the peak compared with the whole-genome average. (C) Minor allele frequency distribution in the 500 kb region centered on the peak compared with the whole-genome average.

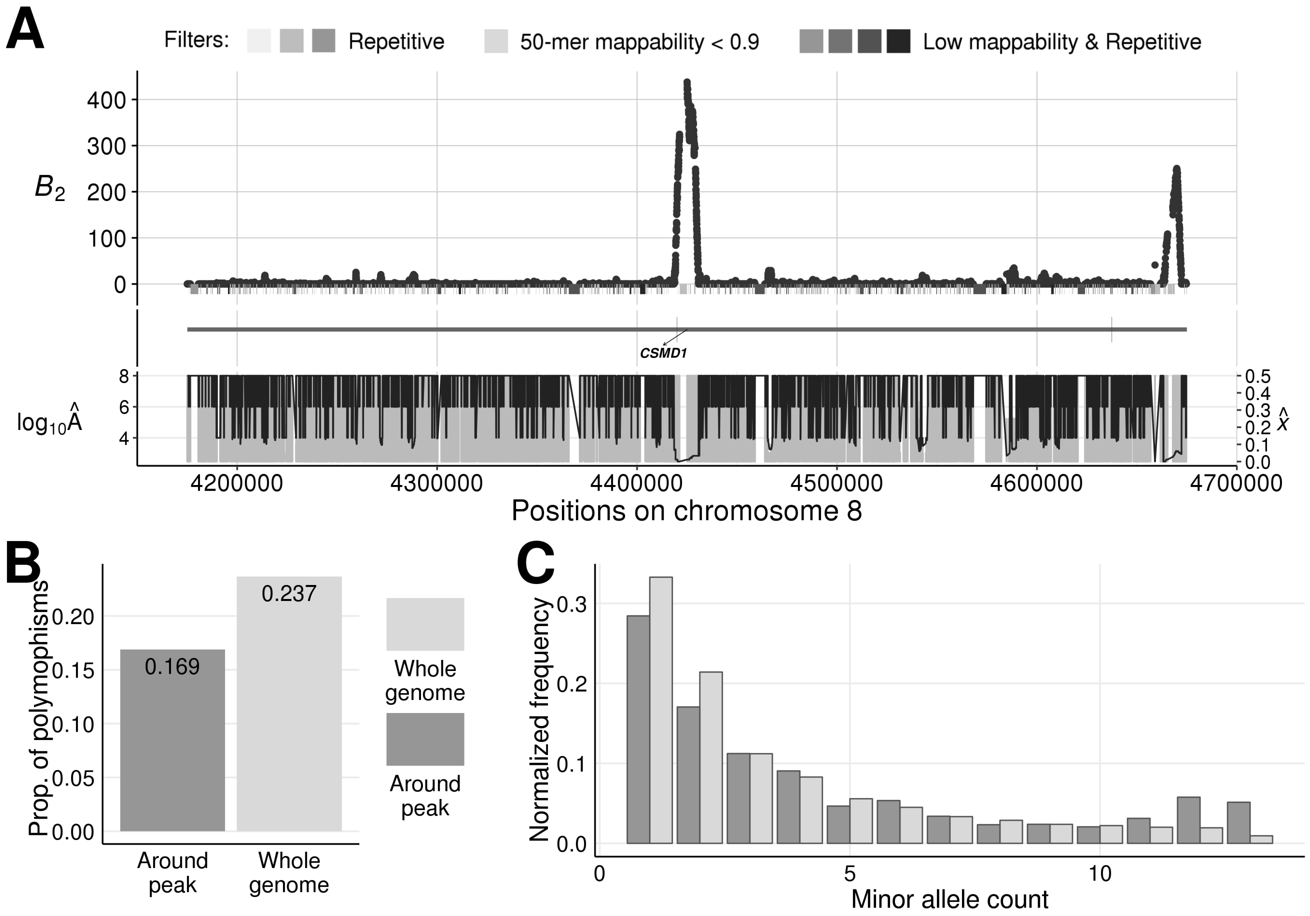


Figure S33: Evidence of balancing selection on *CSMD1*. (A) $B_2$ scores across the 500 kb genomic region on chromosome 8 centered on the peak. The gray bars directly under the $B_2$ scores represent the masked regions, as well as the features in these regions. The darker the shade, the greater number of types of repetitive sequences (*e.g.*, RepeatMasker mask, segmental duplication, simple repeats, or interrupted repeats) overlapping the region. Vertical gray bars below display the estimated equilibrium minor allele frequency $\widehat{x}$ for each maximum likelihood ratio $B_2$, and the black line traces the value for the respective inferred footprint size $\log_{10}(\widehat{A})$. (B) Proportion of informative sites that are polymorphic in the 500 kb region centered on the peak compared with the whole-genome average. (C) Minor allele frequency distribution in the 500 kb region centered on the peak compared with the whole-genome average.

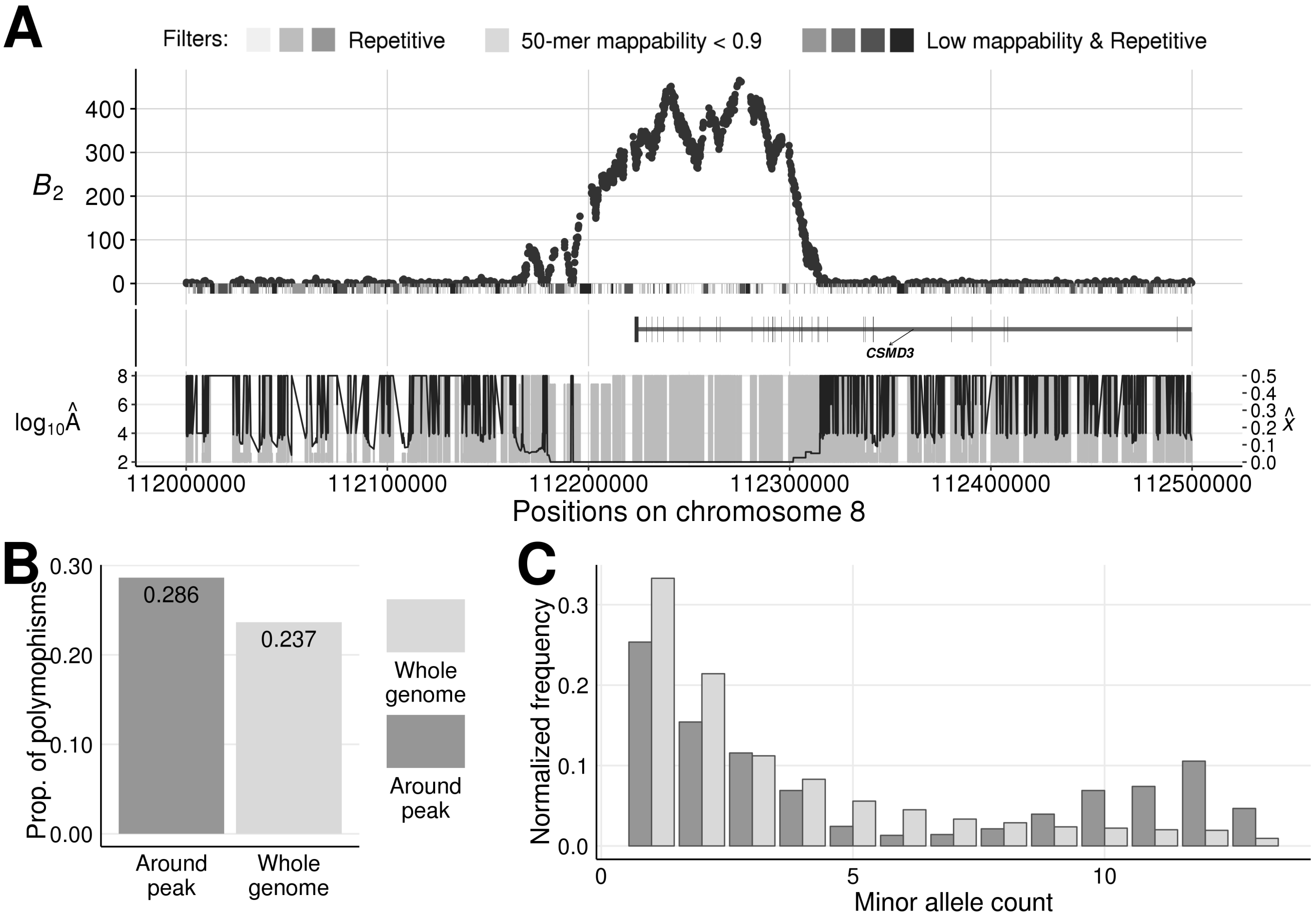


Figure S34: Evidence of balancing selection on *CSMD3*. (A) $B_2$ scores across the 500 kb genomic region on chromosome 8 centered on the peak. The gray bars directly under the $B_2$ scores represent the masked regions, as well as the features in these regions. The darker the shade, the greater number of types of repetitive sequences (*e.g.*, RepeatMasker mask, segmental duplication, simple repeats, or interrupted repeats) overlapping the region. Vertical gray bars below display the estimated equilibrium minor allele frequency $\widehat{x}$ for each maximum likelihood ratio $B_2$, and the black line traces the value for the respective inferred footprint size $\log_{10}(\widehat{A})$. (B) Proportion of informative sites that are polymorphic in the 500 kb region centered on the peak compared with the whole-genome average. (C) Minor allele frequency distribution in the 500 kb region centered on the peak compared with the whole-genome average.

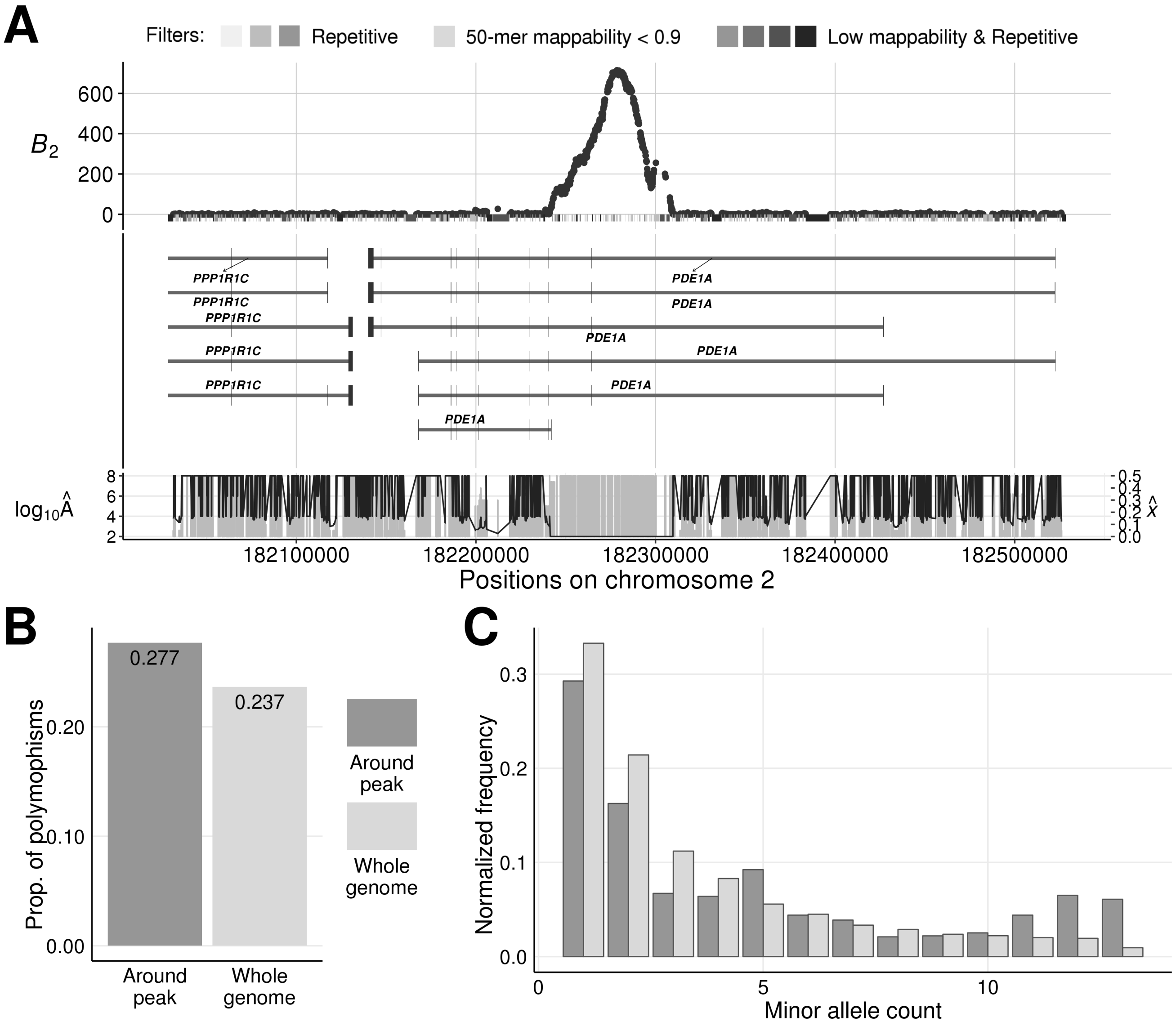


Figure S35: Evidence of balancing selection on *PDE1A*. (A) $B_2$ scores across the 500 kb genomic region on chromosome 2 centered on the peak. The gray bars directly under the $B_2$ scores represent the masked regions, as well as the features in these regions. The darker the shade, the greater number of types of repetitive sequences (*e.g.*, RepeatMasker mask, segmental duplication, simple repeats, or interrupted repeats) overlapping the region. Vertical gray bars below display the estimated equilibrium minor allele frequency $\widehat{x}$ for each maximum likelihood ratio $B_2$, and the black line traces the value for the respective inferred footprint size $\log_{10}(\widehat{A})$. (B) Proportion of informative sites that are polymorphic in the 500 kb region centered on the peak compared with the whole-genome average. (C) Minor allele frequency distribution in the 500 kb region centered on the peak compared with the whole-genome average.

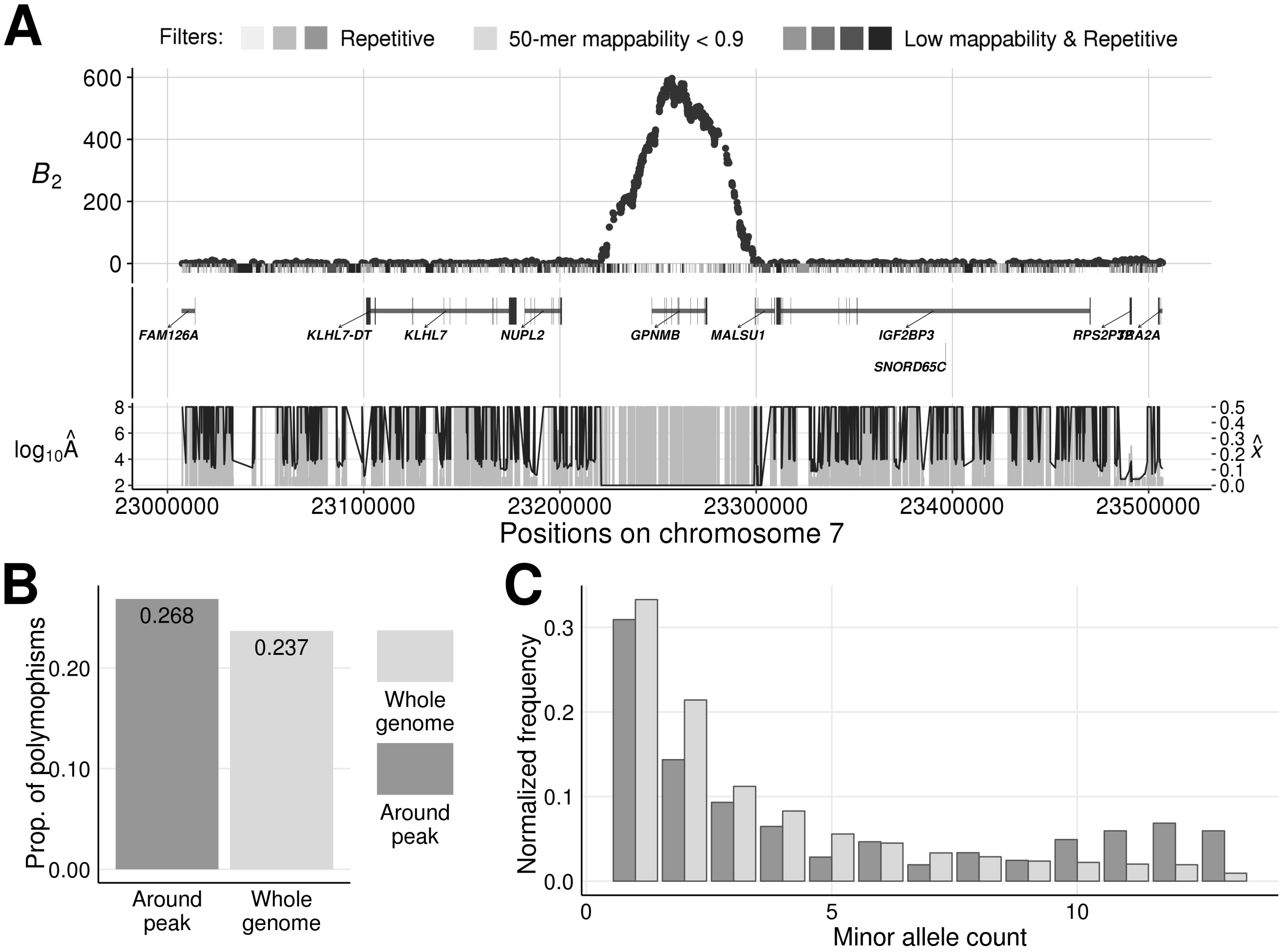


Figure S36: Evidence of balancing selection on *GPNMB*. (A) $B_2$ scores across the 500 kb genomic region on chromosome 7 centered on the peak. The gray bars directly under the $B_2$ scores represent the masked regions, as well as the features in these regions. The darker the shade, the greater number of types of repetitive sequences (*e.g.*, RepeatMasker mask, segmental duplication, simple repeats, or interrupted repeats) overlapping the region. Vertical gray bars below display the estimated equilibrium minor allele frequency $\widehat{x}$ for each maximum likelihood ratio $B_2$, and the black line traces the value for the respective inferred footprint size $\log_{10}(\widehat{A})$. (B) Proportion of informative sites that are polymorphic in the 500 kb region centered on the peak compared with the whole-genome average. (C) Minor allele frequency distribution in the 500 kb region centered on the peak compared with the whole-genome average.

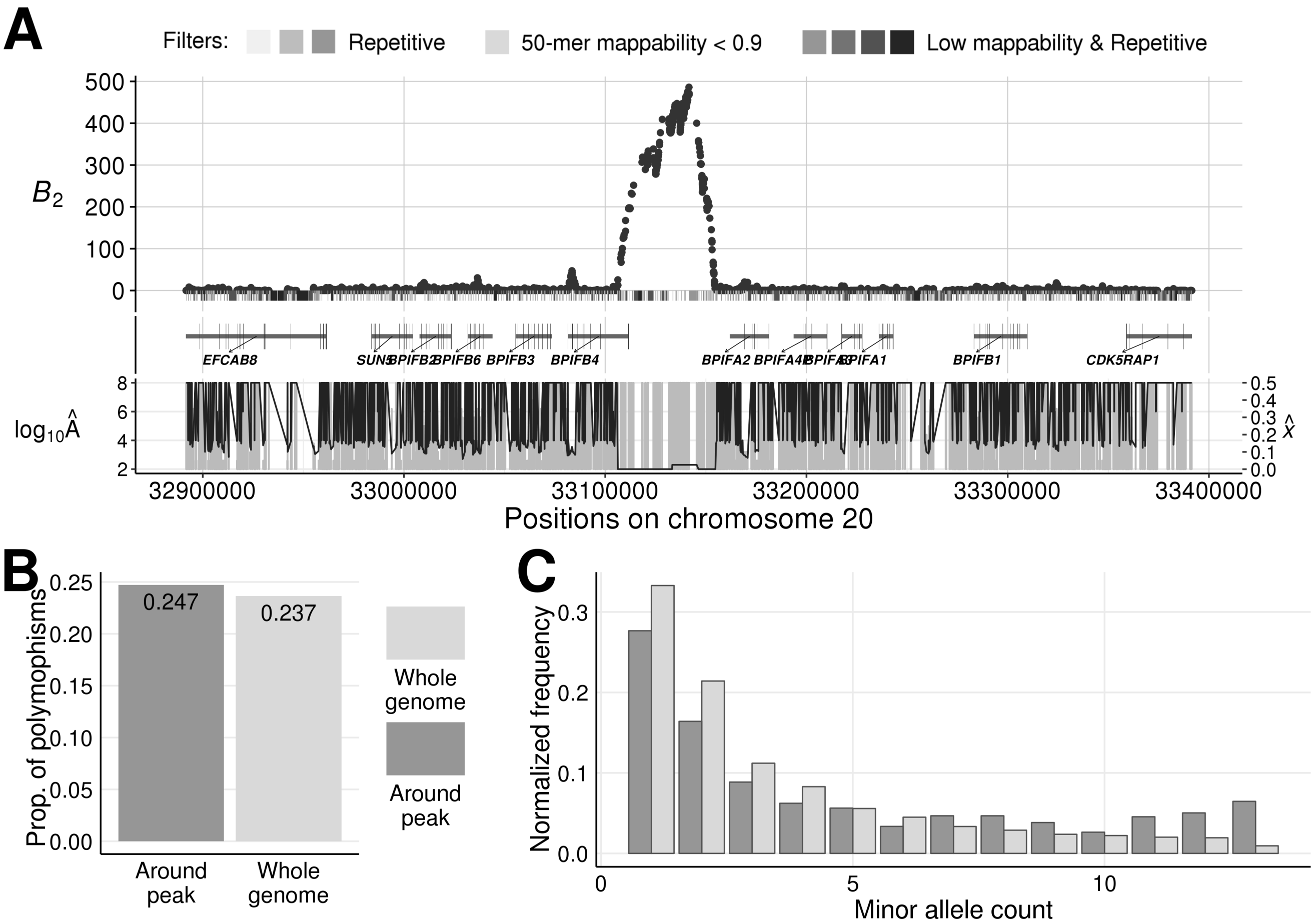


Figure S37: Evidence of balancing selection on the *BPIFB2*/*BPIFA4* intergenic region. (A) $B_2$ scores across the 500 kb genomic region on chromosome 20 centered on the peak. The gray bars directly under the $B_2$ scores represent the masked regions, as well as the features in these regions. The darker the shade, the greater number of types of repetitive sequences (*e.g.*, RepeatMasker mask, segmental duplication, simple repeats, or interrupted repeats) overlapping the region. Vertical gray bars below display the estimated equilibrium minor allele frequency $\widehat{x}$ for each maximum likelihood ratio $B_2$, and the black line traces the value for the respective inferred footprint size $\log_{10}(\widehat{A})$. (B) Proportion of informative sites that are polymorphic in the 500 kb region centered on the peak compared with the whole-genome average. (C) Minor allele frequency distribution in the 500 kb region centered on the peak compared with the whole-genome average.

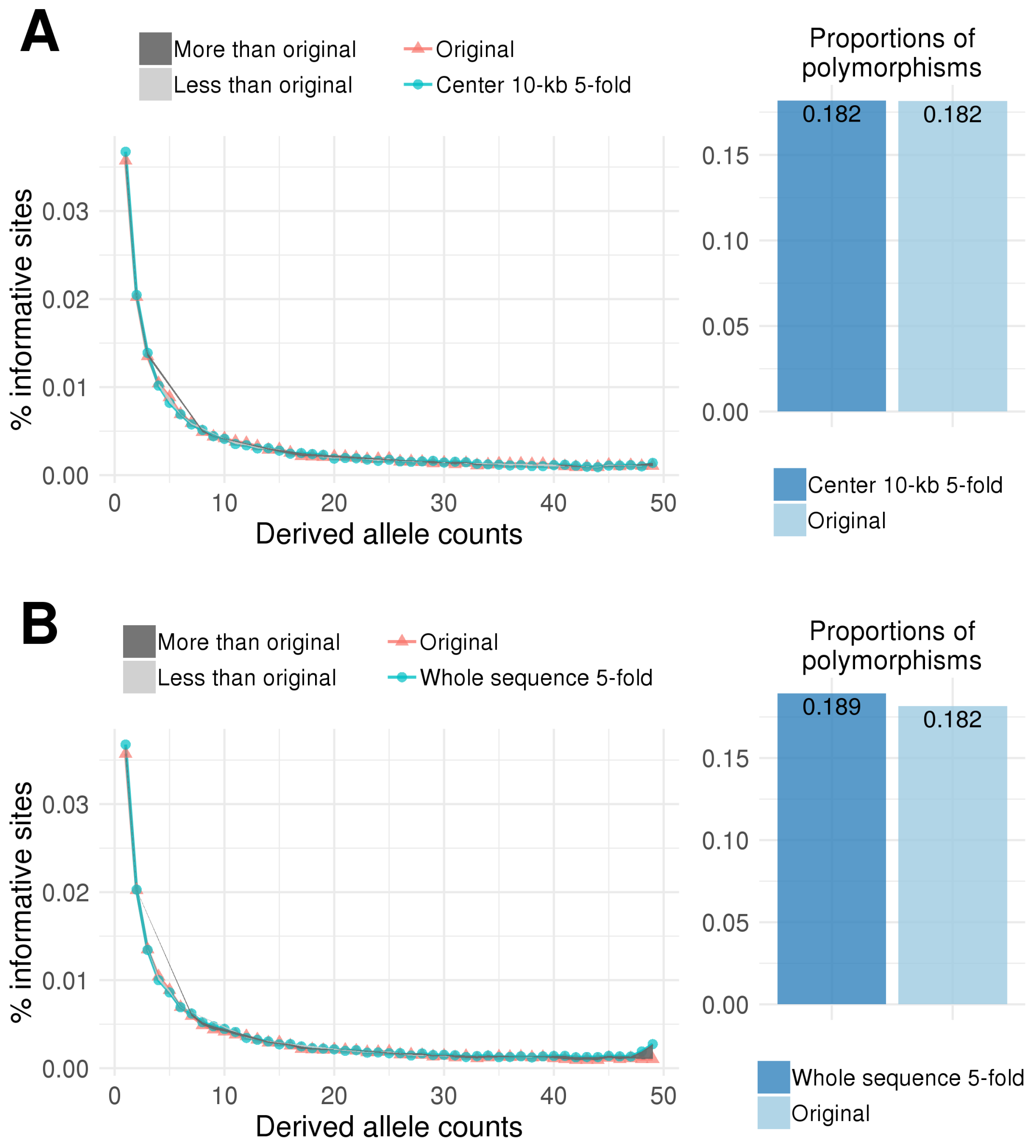


Figure S38: Derived site frequency spectra and proportions of polymorphic sites for (A) sequences with a central 10 kb mutational hotspot of rate $5\mu$, and for (B) sequences with an elevated mutation rate of $5\mu$ across the entire sequence, each compared to sequences with the original mutation rate $\mu$.

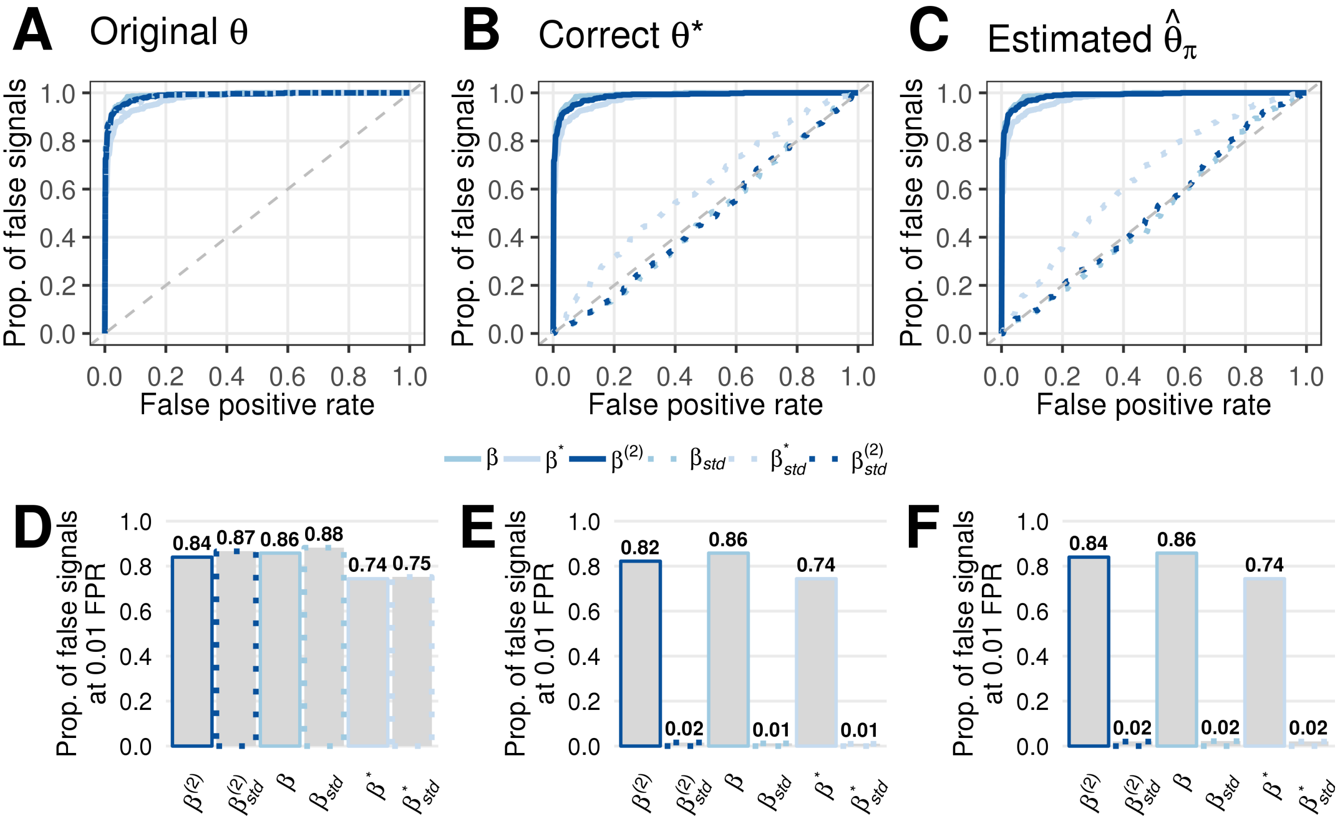


Figure S39: Robustness of $\beta$ statistics on sequences with five-fold mutation rate when provided with different $\theta$ parameters. (A-C) Proportions of false signals reported by each $\beta$ statistic variant as a function of the false positive rate (FPR), and (D-F) proportions of false signals at a 1% FPR when the population-scaled mutation rate $\theta = 4N\mu$ provided tot eh $\beta$ statistics is (A, D) the original un-elevated rate $\theta$, (B, E) the true mutation rate $\theta^\star = 5\theta$, or (C, F) the mean pairwise sequence difference $\widehat{\theta}_\pi$ estimated from the given simulated replicate.

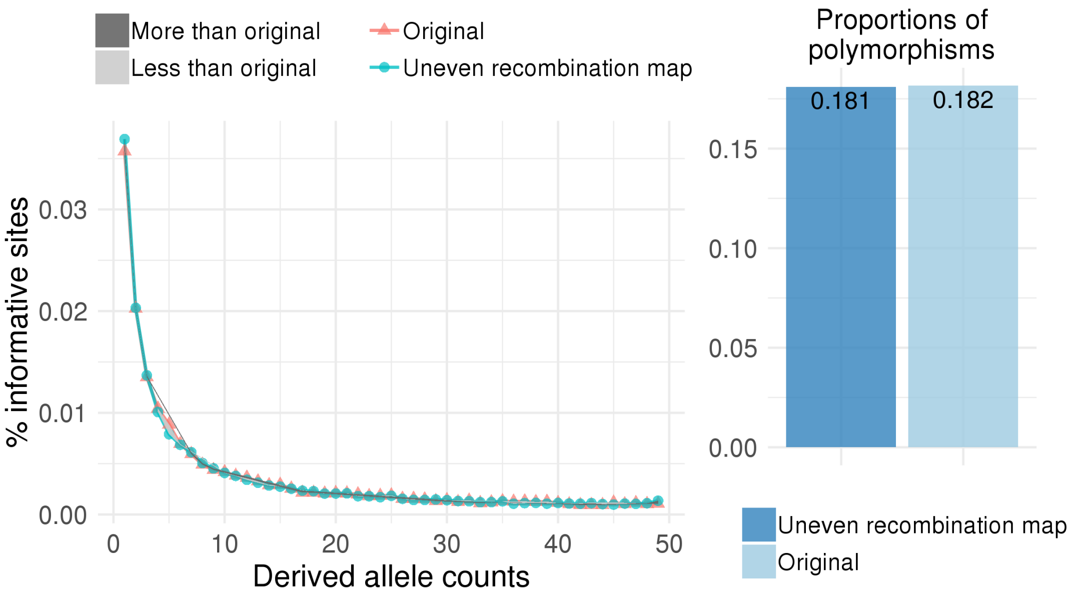


Figure S40: Derived allele frequency spectra and proportions of polymorphic sites for sequences evolving with an uneven recombination map relative to those evolving with a uniform recombination map.

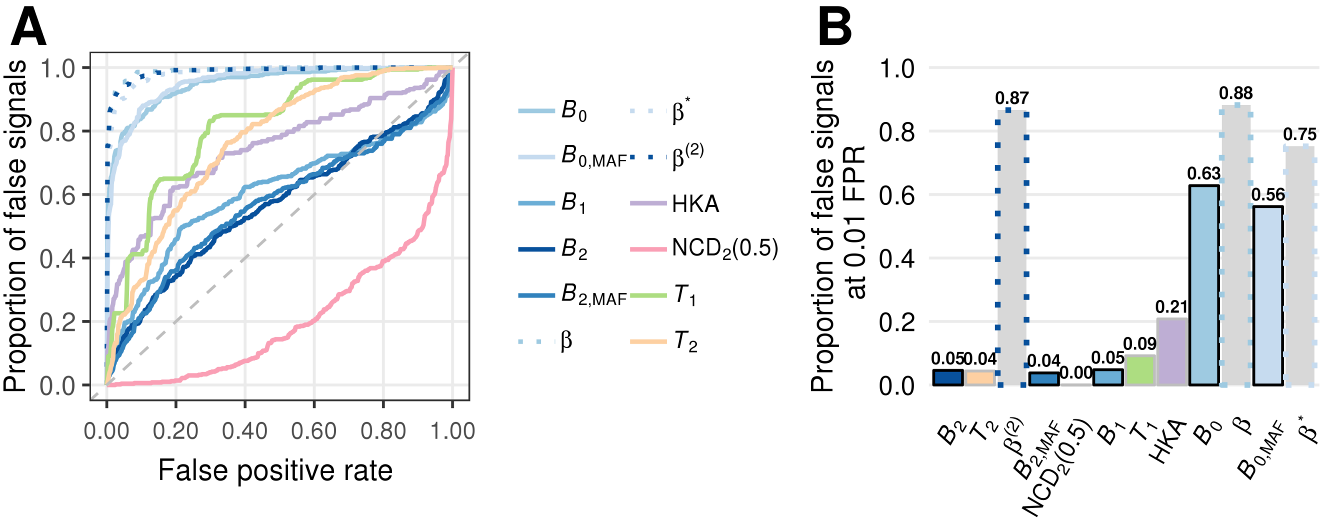


Figure S41: Robustness of $B$ statistics when $B_1$ and $B_2$ variants adopt identical site-based windows as $T_1$ and $T_2$, and $B_0$ variants adopt identical length-based windows as $\beta$ statistics. (A) The proportion of false signals as a function of false positive rates (FPR) for each statisitc. (B) The proportion of false signal at 0.01 FPR for each statistic.

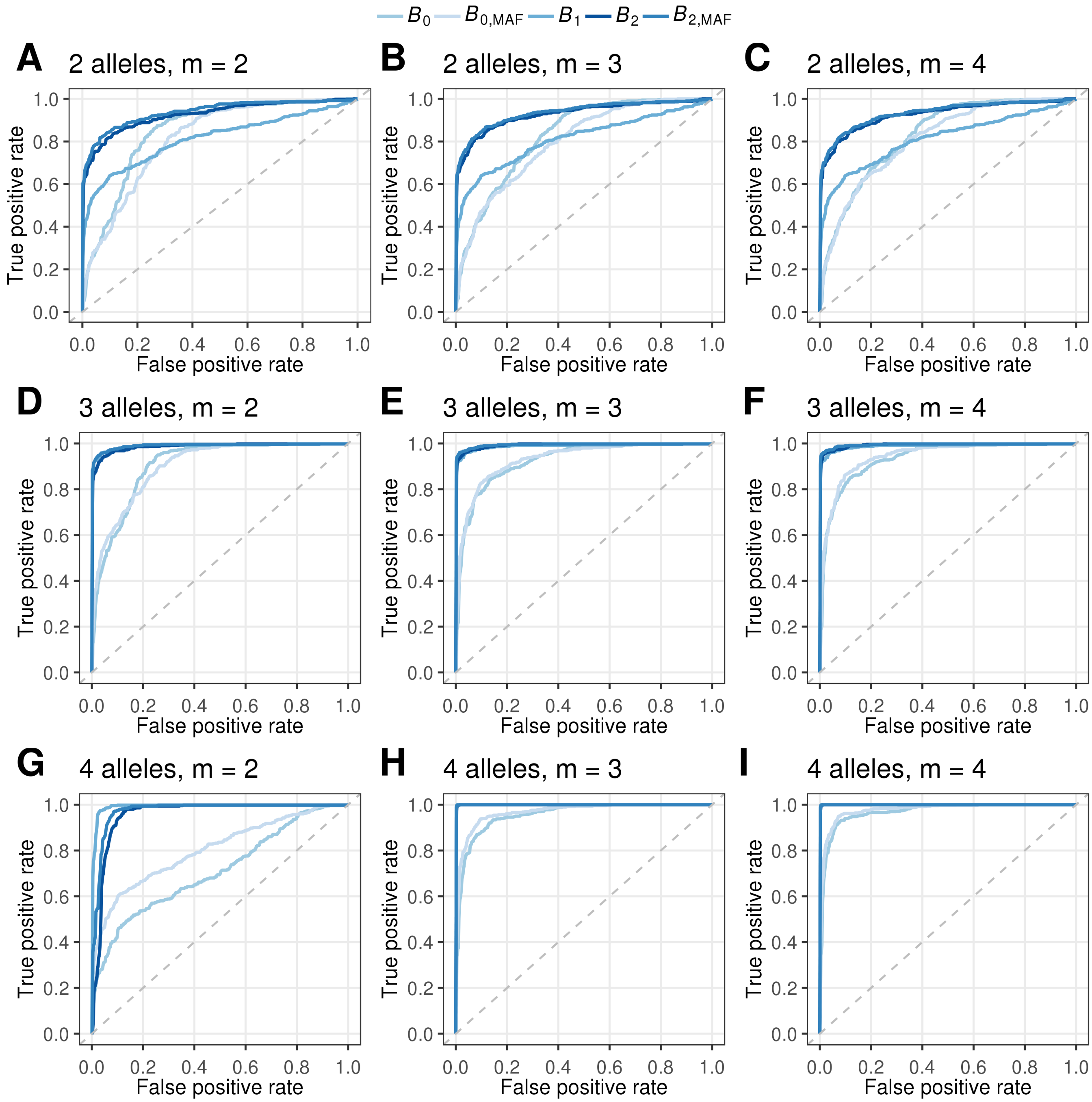


Figure S42: Receiver operating characteristic curves for multi-allelic $B$ statistics when adopting different $m$, the presumed number of balanced alleles. Mutations with selection coefficients $s = 0.001$ with $h = 20$ were introduced 500,000 generations before sampling, with (A-C) two, (D-F) three, or (G-I) four distinct alleles balanced in the population.

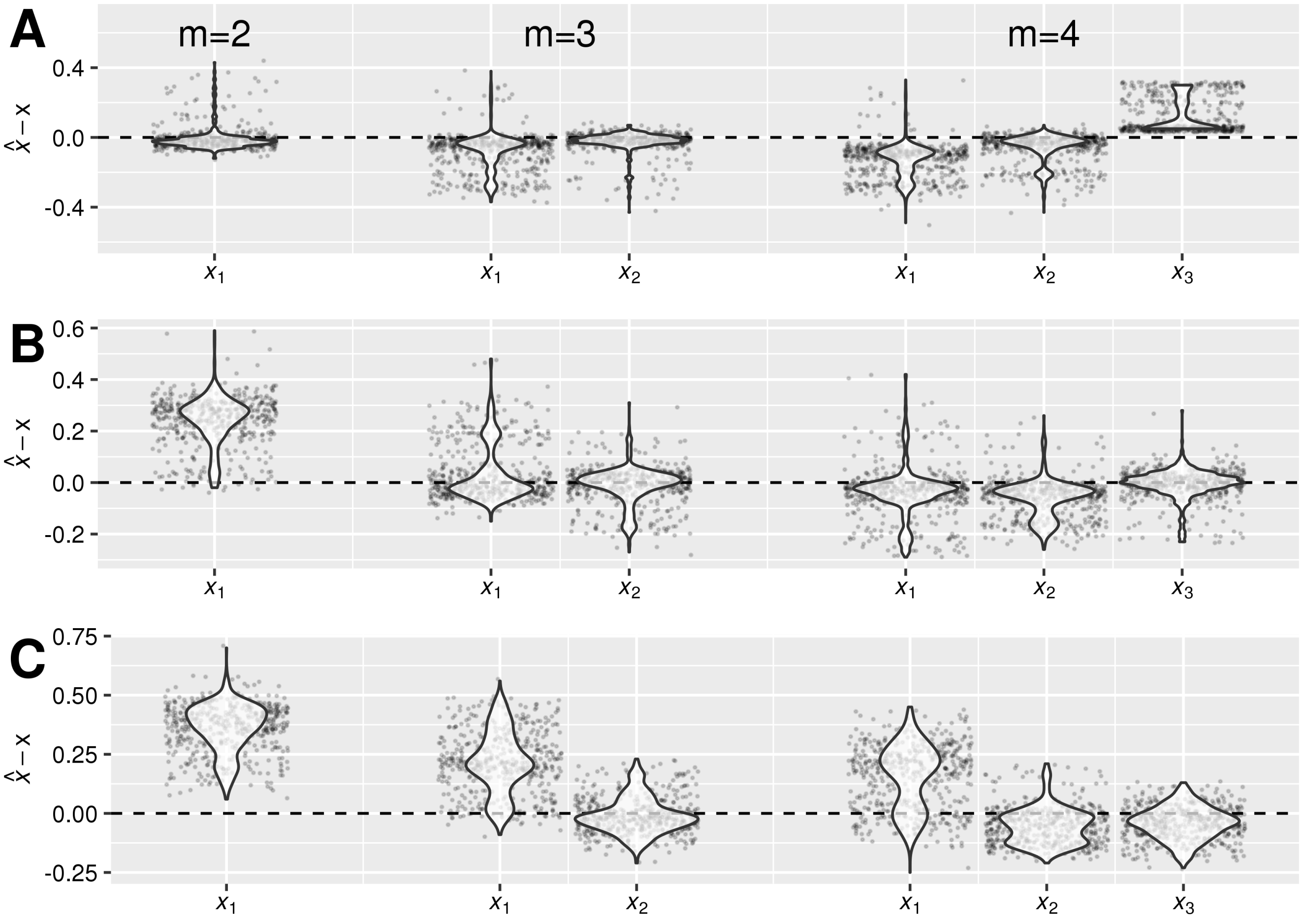


Figure S43: Differences between the optimal equilibrium frequencies ($\hat{x}$) and true allele frequencies of the site under selection reported by $B_2$ statistic when the number of balanced alleles $m$ is assumed to be two, three, or four. Panels represent simulations with (A) two, (B) three, or (C) four distinct alleles balancing at the selected locus.

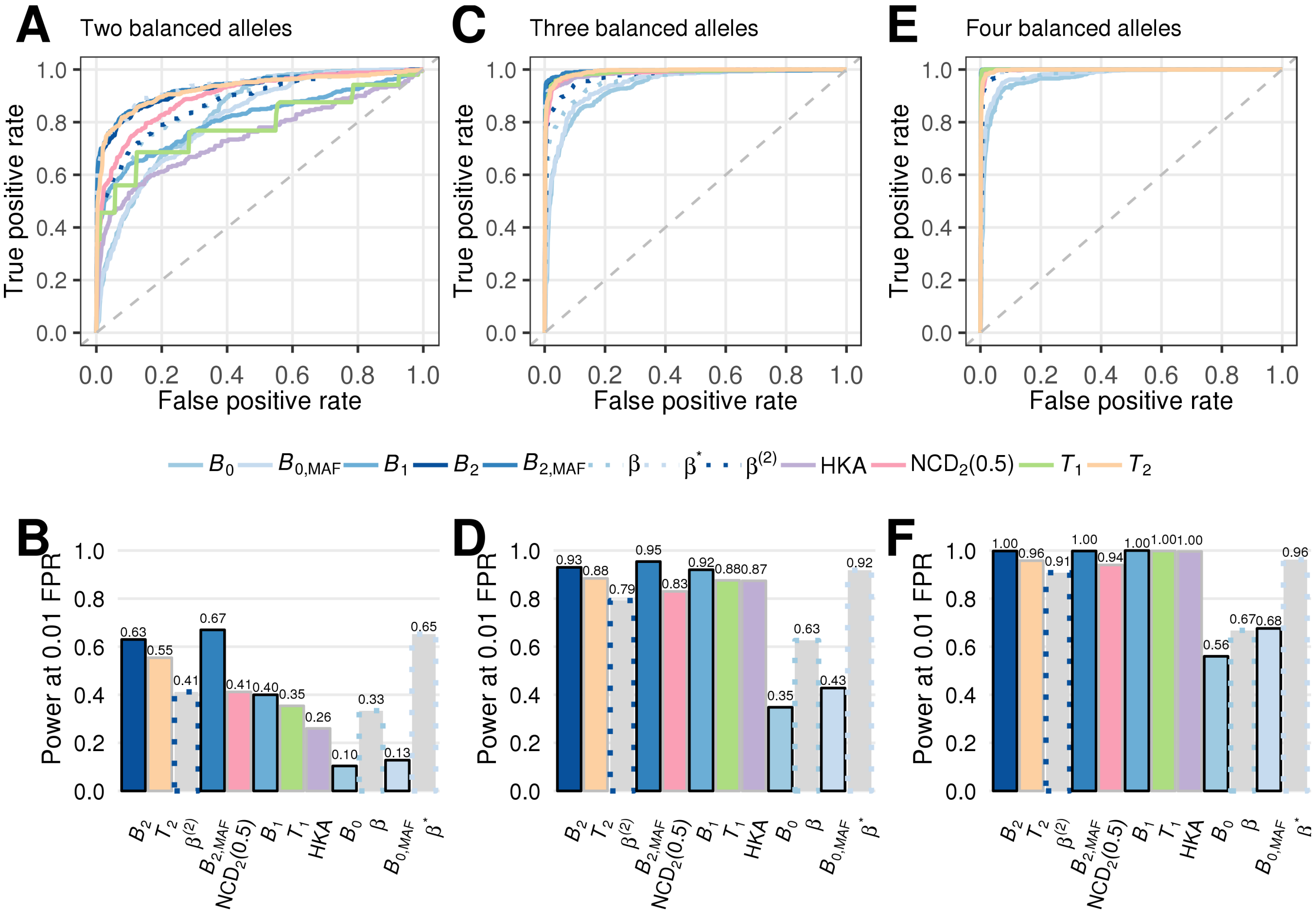


Figure S44: Powers of each statistic to discover genomic regions carrying (A, B) two, (C, D) three, or (E, F) four distinct balanced alleles with $s = 0.001$ and $h = 20$. The $B$ statistics assume $m = 4$. (A, C, E) Receiver operating characteristic curves for each statistic under each scenario. (B, D, F) Powers at a 1% false positive rate (FPR) for each statistic under each of the three scenarios.

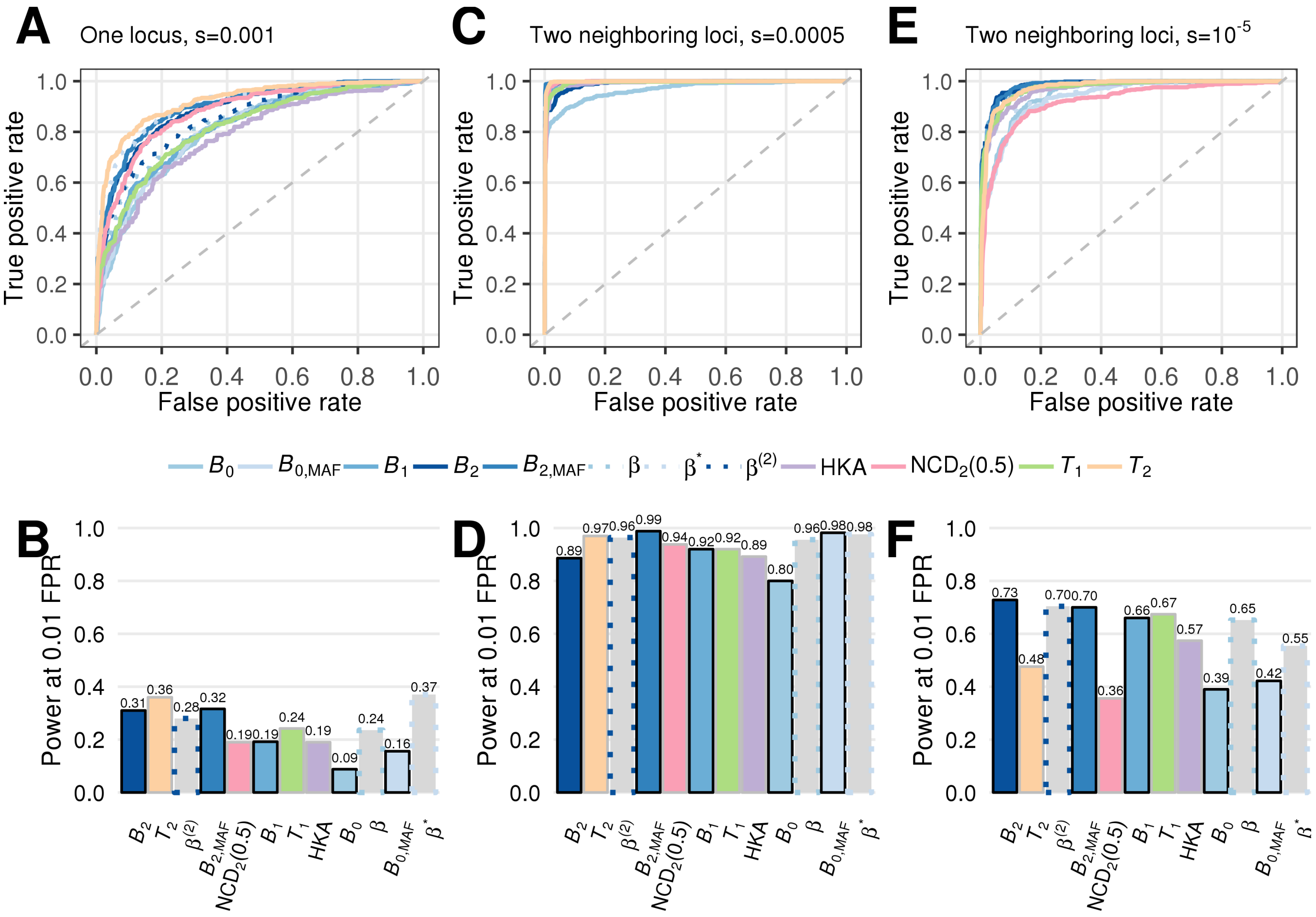


Figure S45: Powers of each statistic to discover genomic regions carrying balanced alleles at (A, B) one locus with $s = 0.001$ and $h = 20$, at (C, D) two neighboring loci 10 kb apart with $s = 0.0005$ and $h = 20$, and at (E, F) two loci 10 kb apart with $s = 10^{-5}$ and $h = 20$. (A, C, E) Receiver operating characteristic curves for each statistic under each scenario. (B, D, F) Powers at a 1% false positive rate (FPR) for each statistic under each of the three scenarios.

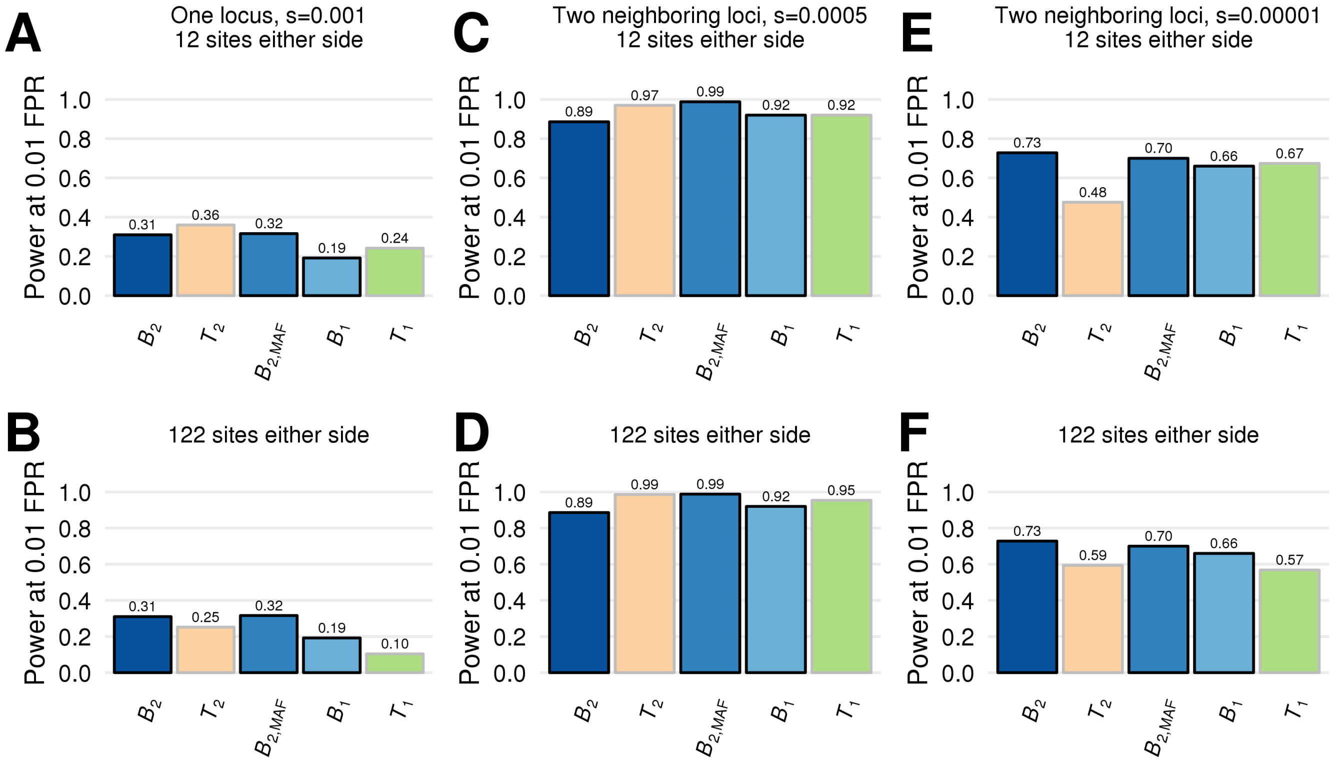


Figure S46: Power at a 1% false positive rate (FPR) of $T$ statistics and their analogous $B$ statistics to discover genomic regions carrying balanced alleles at (A, B) one locus with $s = 0.001$ and $h = 20$, at (C, D) two neighboring loci 10 kb apart with $s = 0.0005$ and $h = 20$, and at (E, F) two loci 10 kb apart with $s = 10^{-5}$ and $h = 20$. $B$ statistics, and $T$ statistics consider (A, C, E) 12 informative sites or (B, D, F) informative sites on either side of each test site.
